# Host-interactor screens of *Phytophthora infestans* RXLR proteins reveal vesicle trafficking as a major effector-targeted process

**DOI:** 10.1101/2020.09.24.308585

**Authors:** Benjamin Petre, Mauricio P. Contreras, Tolga O. Bozkurt, Martin H. Schattat, Jan Sklenar, Sebastian Schornack, Ahmed Abd-El-Haliem, Roger Castells-Graells, Rosa Lozano-Duran, Yasin F. Dagdas, Frank L. H. Menke, Alexandra M. E. Jones, Jack H. Vossen, Silke Robatzek, Sophien Kamoun, Joe Win

**Author notes:** These authors contributed equally to this work (Benjamin Petre, Mauricio P. Contreras).

## Abstract

Pathogens modulate plant cell structure and function by secreting effectors into host tissues. Effectors typically function by associating with host molecules and modulating their activities. This study aimed to identify the host processes targeted by the RXLR class of host-translocated effectors of the potato blight pathogen *Phytophthora infestans.* To this end, we performed an *in planta* protein-protein interaction screen by transiently expressing *P. infestans* RXLR effectors in *Nicotiana benthamiana* leaves followed by co-immunoprecipitation (co-IP) and liquid chromatography tandem mass spectrometry (LC-MS/MS). This screen generated an effector-host protein interactome matrix of 59 *P. infestans* RXLR effectors x 586 *N. benthamiana* proteins. Classification of the host interactors into putative functional categories revealed over 35 biological processes possibly targeted by *P. infestans.* We further characterized the PexRD12/31 family of RXLR-WY effectors, which associate and co-localize with components of the vesicle trafficking machinery. One member of this family, PexRD31, increased the number of FYVE positive vesicles in *N. benthamiana* cells. FYVE positive vesicles also accumulated in leaf cells near *P. infestans* hyphae, indicating that the pathogen may enhance endosomal trafficking during infection. We anticipate that the interactome dataset we generated will serve as a useful community resource for functional studies of *P. infestans* effectors and of effector-targeted host processes.

## INTRODUCTION

Plant pathogens reprogram host cells to their advantage to establish successful infections (Dodds and Rathjen, 2010). Understanding how pathogens manipulate their hosts will advance our knowledge of infection processes and help us develop disease resistant crops (Alfano, 2009; Gawehns *et al.*, 2013; van Schie and Takken, 2014). Pathogens modulate plant cell structure and function by secreting effectors into host tissues. Effectors generally operate by binding or enzymatically modifying host molecules. These host molecules are classified in two major functional classes: (i) ‘targets’ directly modified by effectors to modulate host processes; (ii) ‘helpers’ or ‘facilitators’ co-opted by effectors to execute their activities, for instance to traffic within host cells (Win *et al.*, 2012b). Some effectors can be recognized by host immune receptors, consequently triggering an immune response. Such effectors are said to have an avirulence (AVR) activity, as their recognition often results in loss of virulence of the pathogen in hosts carrying a matching immune receptor (Dodds and Rathjen, 2010). Despite major advances in effector biology, our understanding of effector-targeted processes remains fragmentary and generally centered on suppression of innate immunity (Zheng *et al.*, 2014; Sharpee and Dean, 2016; Toruño *et al.*, 2016; Ren *et al.*, 2019).

Effectors modulate a variety of pathways and therefore can be leveraged as molecular probes to reveal novel and important processes in host cells (Wei *et al.*, 2012; Win *et al.*, 2012b; Lee *et al.*, 2013; Toruño *et al.*, 2016). Pathogen effectors generally associate with host protein complexes to function (Asai and Shirasu, 2015; Boevink *et al.*, 2016). Identifying these host proteins is the first step in unravelling the host processes modulated by effectors. To this end, over the last decade effector biologists have performed large-scale effector-host protein interaction screens using yeast two-hybrid or *in planta* coimmunoprecipitation/tandem mass spectrometry (coIP/MS) assays (Mukhtar *et al.*, 2011; Weßling *et al.*, 2014; Petre *et al.*, 2015; Petre *et al.*, 2016). While yeast-two hybrid identifies one-to-one protein associations, coIP/MS assays tend to reveal multiple host proteins that associate with a given effector in protein complexes (Win *et al.*, 2011; Petre *et al.*, 2017). This feature makes coIP/MS assays particularly suitable for identifying processes targeted by effectors. Moreover, the increasing availability of plant genome sequences allows for improved proteome predictions that greatly aid IP/MS based effector interactor searches.

The Irish potato famine pathogen *Phytophthora infestans* (oomycete, Peronosporales) is a major threat to potato and tomato crops worldwide (Fisher *et al.*, 2012; Kamoun *et al.*, 2015; Derevnina *et al.*, 2016). Peronosporales species secrete a superfamily of effector proteins known as RXLR effectors; named after the conserved Arginine-any amino acid-Leucine-Arginine motif that follows the signal peptide of the proteins (Jiang *et al.*, 2008; Win *et al.*, 2012a). The *P. infestans* genome harbors over 500 predicted RXLR effectors that are grouped into ~150 families and tend to exhibit sequence and expression polymorphisms between pathogen strains (Haas *et al.*, 2009; Cooke *et al.*, 2012; Yoshida *et al.*, 2013; Pais *et al.*, 2018). RXLR effectors are modular proteins with the N-terminal signal peptide and RXLR region mediating secretion and translocation into host cells, and the C-terminal end, often defined by the WY/LWY fold, carrying the effector biochemical activity (Whisson *et al.*, 2007; Win *et al.*, 2012a; He *et al*., 2019). To date, over a dozen *P. infestans* RXLR effectors have been functionally characterized for their virulence activities (Rovenich *et al.*, 2014; Anderson *et al.*, 2015; Du *et al.*, 2015; Wang *et al.*, 2015; Boevink *et al.*, 2016; Dagdas *et al.*, 2016; Yang *et al.*, 2016; Turnbull *et al.*, 2017; Wang *et al.*, 2018). These effectors bind and alter the stability, activity, or subcellular localization of a diversity of host proteins, including proteases, kinases, phosphatases, transcription factors, ubiquitin ligases, RNA binding proteins, autophagy-related proteins, and vesicular trafficking-associated proteins. The emerging picture is that *P. infestans* RXLR effectors modulate multiple host processes to the pathogen’s benefit. Nonetheless, systematic interactome screens between *P. infestans* RXLR effectors and host proteins have not been reported to date.

Many plant pathogenic and symbiotic microbes produce specialized hyphae that invade host cells but remain enveloped by newly synthesized host-derived membranes (Oliveira-Garcia and Valent, 2015; Ivanov *et al.*, 2019; Bozkurt and Kamoun, 2020). Processes taking place at this host-pathogen interface are thought to have a major impact on the outcome of the interaction. The mechanisms underlying the biogenesis and functions of host–microbe interfaces remain poorly understood although recent advances have been made (Bozkurt and Kamoun, 2020; Rausche *et al.*, 2020). Well-studied examples of these specialized hyphae are haustoria formed by *P. infestans* and other filamentous pathogens. During infection, haustoria enter the plant cell cavity and become surrounded by the plant-derived extrahaustorial membrane (EHM). *P. infestans* effectors secreted at this pathogen-host interface are thought to reprogram multiple processes in the invaded (haustoriated) host cells (Bozkurt *et al.*, 2012; Dagdas *et al.*, 2016; Dagdas *et al.*, 2018). In addition, four *P. infestans* RXLR effectors, AVRblb2, AVR1, AVR2, and PexRD54, focally accumulate around haustoria when they are heterologously expressed in host cells (Bozkurt *et al.*, 2011; Saunders *et al.*, 2012; Dagdas *et al.*, 2018; Wang *et al.*, 2018). These findings, along with related cell biology and biochemical studies, indicate that *P. infestans* massively reprograms host membrane trafficking during infection (Du *et al.*, 2015; Tomczynska *et al.*, 2018). Membrane trafficking perturbations of haustoriated cells include the rerouting to the haustorial interface of components of the endocytic pathways (Lu *et al.*, 2012; Bozkurt *et al.*, 2015), autophagy machinery (Dagdas *et al.*, 2018; Pandey *et al.*, 2020) and chloroplasts (Toufexi *et al.*, 2019).

This study aims at identifying the host plant proteins that are associated with a representative set of *P. infestans* RXLR effectors to generate a pathogen-host protein interactome and gain an overview of the diversity of effector-targeted processes in this pathosystem. To achieve our aim, we performed an *in planta* coIP/MS screen in the model plant *Nicotiana benthamiana,* which has emerged as an optimal experimental system for cell biology and biochemistry studies of *P. infestans-host* interactions (Bozkurt and Kamoun 2020). In addition, *N. benthamiana* is particularly appropriate for large-scale coIP/MS screens thanks to rapid transient protein expression using *Agrobacterium tumefaciens* (agroinfiltration) and the availability of a genome sequence that improves MS proteome predictions (Goodin *et al*., 2008; Derevnina *et al.*, 2019; Zess *et al.*, 2019). Using this approach, we generated a host protein-pathogen effector interactome matrix of 59 *P. infestans* RXLR effectors x 586 *N. benthamiana* proteins. This interactome revealed 35 candidate effector-targeted processes in *N. benthamiana.* We further characterized the PexRD12/31 family of RXLR-WY effectors, which associate and co-localize with components of the vesicle trafficking machinery. One member of this family, PexRD31, increased the number of 2xFYVE-GFP labelled vesicles in *N. benthamiana* cells. We also noted that *P. infestans* alters the number and the distribution of 2xFYVE-GFP vesicles in *N. benthamiana* leaves, indicating that the pathogen dramatically alters host endosomal trafficking during infection.

## RESULTS

### *In planta* coimmunoprecipitation/mass spectrometry (coIP/MS) assays reveal host proteins associated with *Phytophthora infestans* RXLR effectors

To identify host protein complexes associated with representative *P. infestans* RXLR effectors, we performed a protein-protein interaction screen *in planta* using a pipeline similar to the one we previously described for candidate effector proteins of the poplar leaf rust fungus (Petre *et al.*, 2015). We first selected 66 effectors from 14 prominent RXLR families, which are enriched in effectors that are transcriptionally induced during plant infection and carrying AVR activities (Table 1) (Oh *et al.*, 2009; Vleeshouwers *et al.*, 2011). We then expressed N-terminal FLAG-tagged effector proteins individually in *N. benthamiana* leaf cells by agroinfiltration and performed anti-FLAG coIP/MS to identify host protein complexes associated with each effector (Figure S1, see methods for details). We evaluated the expression of the effector fusions by analyzing immunoprecipitated protein mixtures using sodium dodecyl sulphate-polyacrylamide gel electrophoresis (SDS-PAGE) coupled with Colloidal Coomassie Blue (CCB) staining (Figure S2). Out of 66 effectors, 59 accumulated to adequate levels for coIP and identification by mass spectrometry. Spectral searches against the *N. benthamiana* proteome revealed 967 host protein interactors for the 59 effectors (Dataset 1).

**Table 1.**
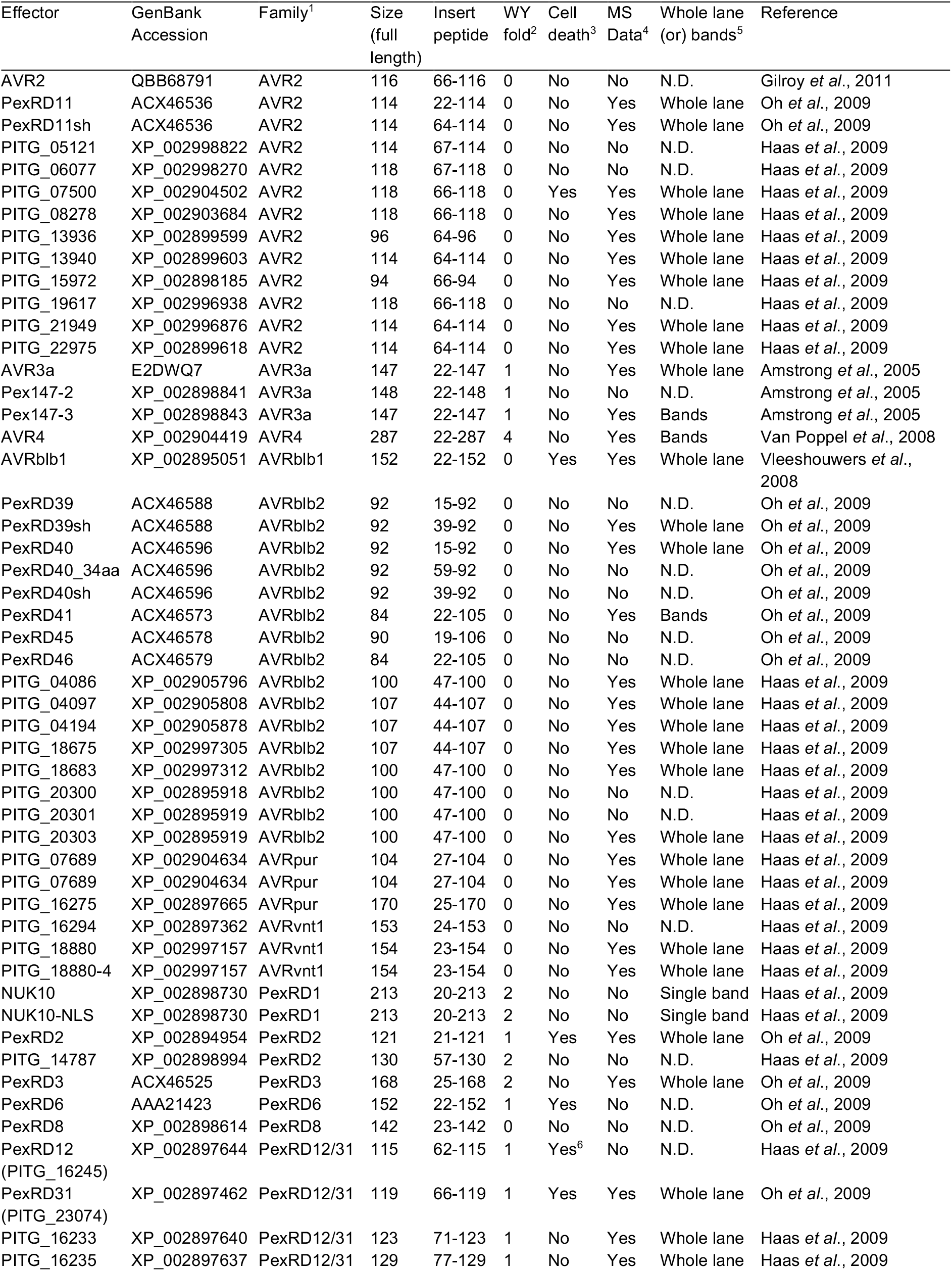
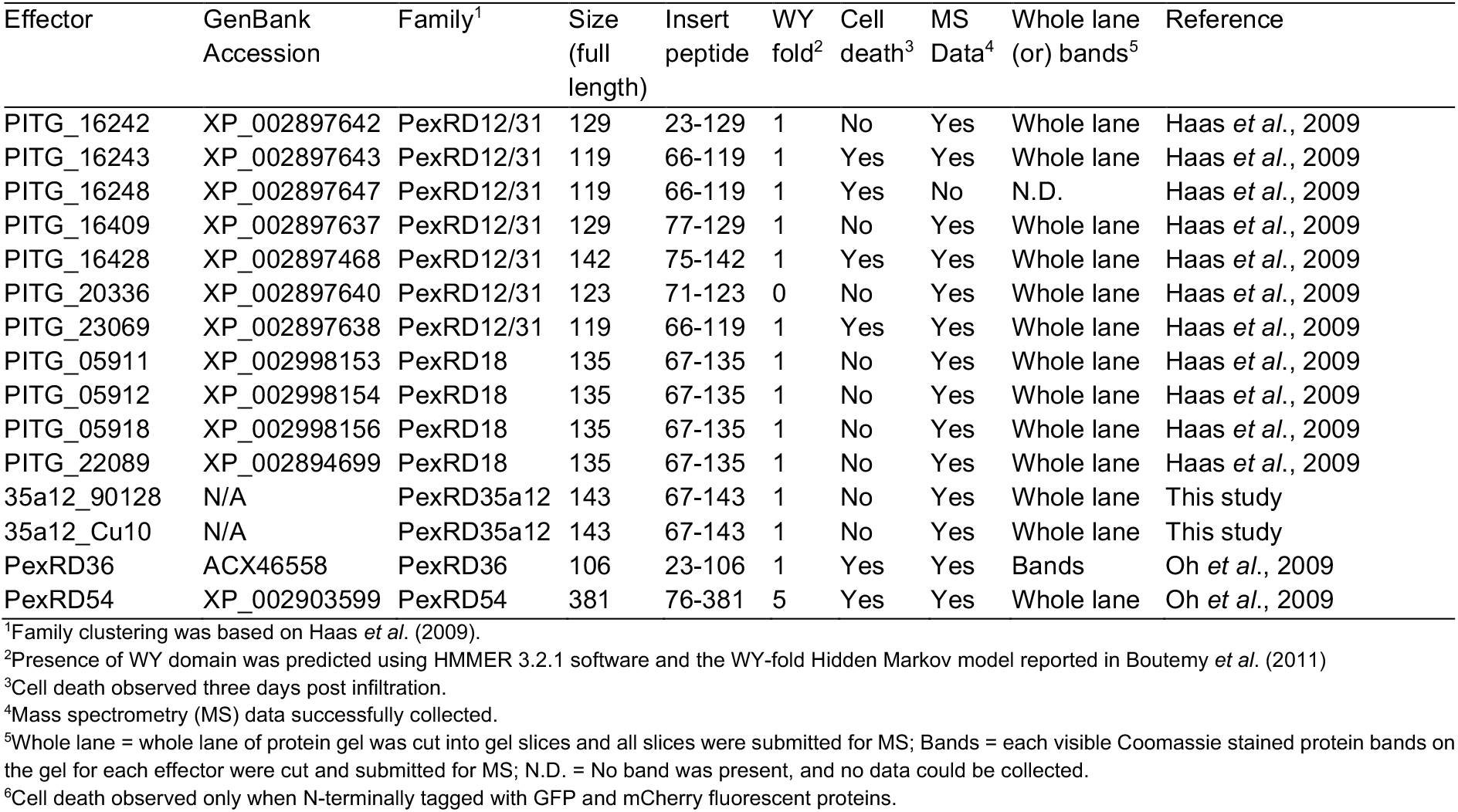
*Phytophthora infestans* RXLR effectors used in co-immunoprecipitation study

Next, we deconvoluted the 59 x 967 matrix to facilitate the interpretation of the results. Firstly, given that multiple RXLR effectors are related and analyses of individual effectors would skew the results towards the largest families, we merged the data for the 59 individual effectors into 14 sequence-related families (Dataset 2). In total, 7 effectors remained as singletons whereas the others grouped into families that range in size from 2 to 16 (AVRblb2 being the largest family) (Table 1). Secondly, to further reduce the matrix complexity, we clustered host protein into groups having at least 80% identity over at least 80% of length. This yielded a total of 586 host proteins, which were represented by a single protein entry to make a consolidated interactome matrix (Dataset 2). We used this interactome network for further analyses. On average, a given effector family associated with 105 host proteins (range 5 to 348), with the PexRD12/31 family associating with the largest number of proteins (Figure 1A). *N. benthamiana* interactor proteins associated in average with 2.5 effector families (Figure 1B).

**Figure 1.**
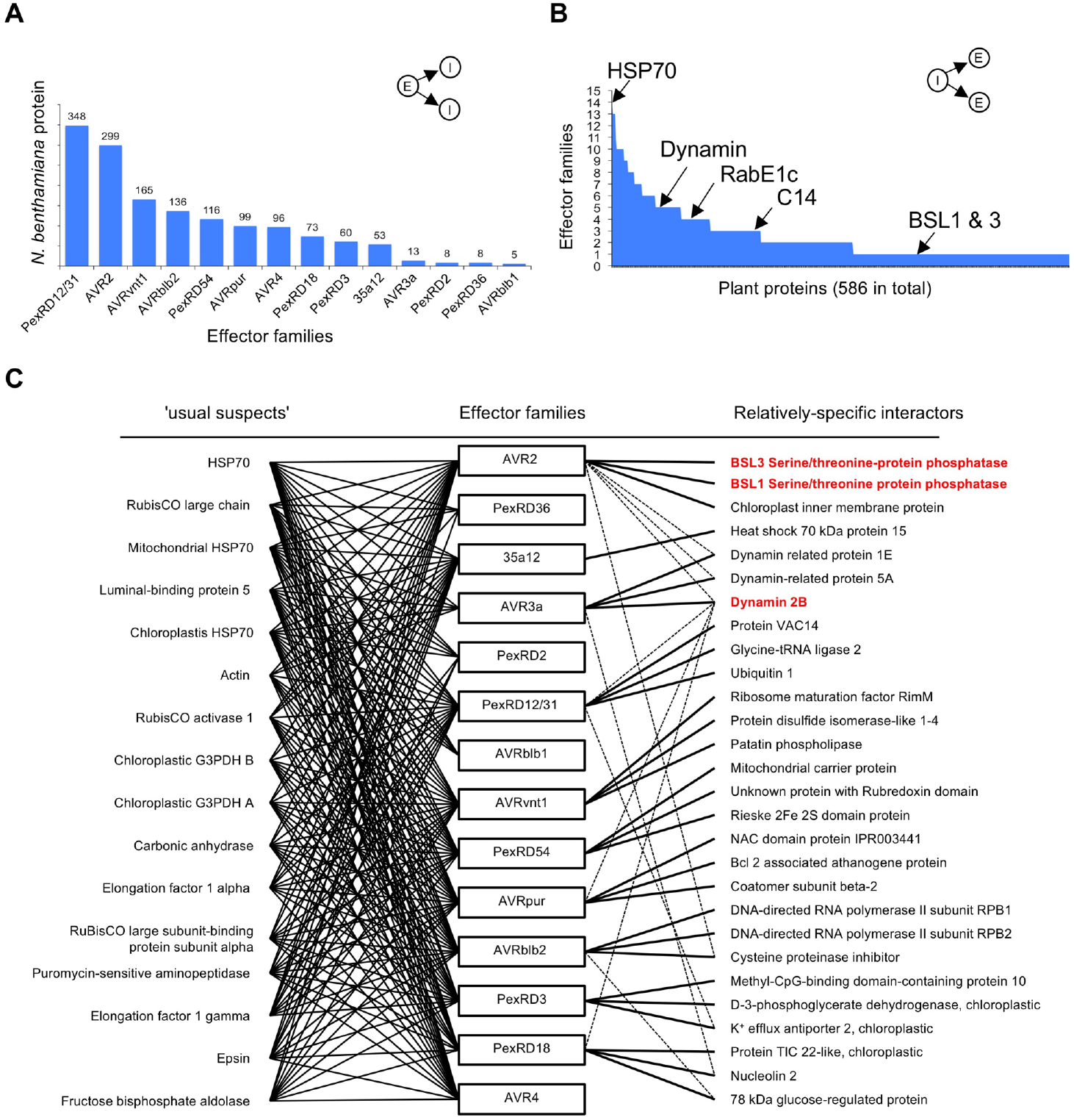
Overview of the *Phytophthora infestans* RXLR effector x *Nicotiana benthamiana* protein interactome. **(A**) Number of plant proteins that associate with each of the tested 14 RXLR effector families of *P. infestans.* E = effector family; I = host interactor. **(B)** Number of effector families that associate with each *N. benthamiana* protein. For example, HSP70 associated with all 14 effector families while previously experimentally validated interactors BSL1, BSL3, C14, dynamin, and RabE1c associated with one to five effector families. **(C)** Interactome network of most promiscuous *N. benthamiana* proteins (also referred to as the ‘usual suspects’, shown on the left) and a selection of relatively-specific interactors (on the right) for each effector family. The usual suspects include plant proteins associated with more than nine different effector families. The relatively-specific set includes top three *N. benthamiana* proteins identified in coIP/MS from each effector family by following criteria: have at least three peptides hits; have the maximum peptide hits in coIP/MS of the effector family they associate within the data set; and show association with the least number (up to five) of effector family. Effector families PexRD36, PexRD2, AVRblb1 and AVR4 did not show association with plant proteins that fulfil the aforementioned criteria. The dotted lines indicate associations that do not have the maximum number of peptide hits in coIP/MS within the dataset. The plant proteins shown in red bold text have been experimentally validated for their association with respective effector families.

### *Phytophthora infestans* RXLR effector families associate with host protein complexes that include previously validated interactors

One limitation of affinity purification-mass spectrometry assays is that they tend to yield false positives, notably abundant and sticky proteins (Mellacheruvu *et al*., 2013; Petre *et al*., 2015). Given that the same false positives tend to appear in independent experiments, interactors that are specific to a smaller subset of bait proteins are more likely to be biologically relevant. We therefore flagged the most promiscuous interactors that associated with >9 effector families as likely representing false positives (Figure 1C). These 16 “usual suspect” interactors overlap with the list previously reported by Petre *et al.* (2015) for *N. benthamiana* interactors of candidate effectors from the poplar rust fungus.

Five host interactors of *P. infestans* RXLR effectors in our interactome matrix have been previously validated (Table S1). The cysteine protease RD21a (C14) associates with members of the AVRblb2 family (Bozkurt *et al.*, 2011; Wang *et al.*, 2015), the BSU1-like serine/threonine protein phosphatases BSL1 and BSL3 associate with members of the AVR2 family (Saunders *et al.*, 2012), dynamin 2B associates with the AVR3a family (Chaparro-Garcia *et al.*, 2015) and Ras-related protein RabE1c (also known as Rab8a) associates with PexRD54 (Pandey *et al.*, 2020) (Table S1, Figure 1C). These validated host interactors associated with anywhere from 1 to 5 effector families in our experiments (Table S1). Therefore, we reasoned that host interactors showing association with five or less effector families would likely represent relatively specific interactors. Using this cut-off, we identified a set of 497 near-specific host interactors in our matrix (Dataset 2). We conclude that the interactome matrix we generated is likely to contain biologically relevant host protein-effector associations and warrants further investigation.

### Interactome network analyses reveal putative host processes targeted by *Phytophthora infestans* RXLR effector

The host interactors of the RXLR effectors can serve as proxies to identify candidate host processes targeted by *P. infestans.* To this end, we categorized the 586 host interactors according to their Gene Ontology (GO) terms and built a GO-based interactome network from protein-to-protein binary interactions (Figure 2A, Table S2). Nodes represent plant proteins or effector families and edges represent the interaction between two proteins. Classification of the host interactors into putative functional GO categories revealed about 35 biological processes that are potentially targeted by effectors (Table S2). The network revealed that 50% of the interactions involved host proteins from three main GO categories: translation, metabolic process, and photosynthesis. Some of these may be false positives. For example, we anticipated that proteins involved in translation would associate with effectors during *in planta* expression and may be difficult to study as bona fide effector targets.

**Figure 2.**
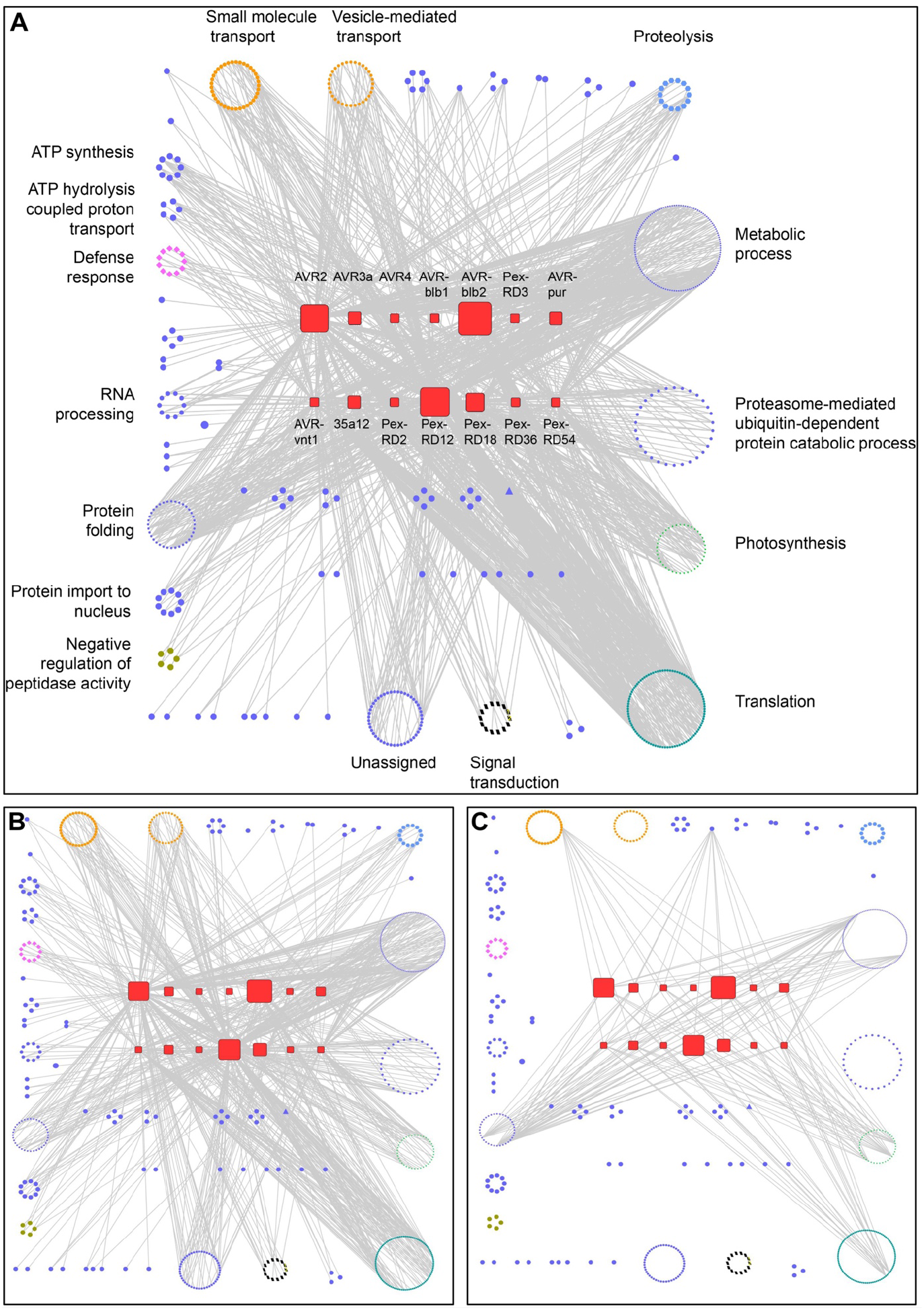
Interactome network analyses reveal host biological processes likely targeted by *Phytophthora infestans* RXLR effectors. **(A)** General network overview of the association between the RXLR effector families and the host interactors grouped according to their predicted biological processes using Gene Ontology (GO) terms. **(B)** Same as (A), with only the relatively specific associations (i.e. host interactors associated with < 5 effector families) depicted. **(C)** Same as (A), with only the redundant associations (i.e. host interactors associated with > 9 effector families) depicted.

To clarify the network topology, we generated sub-networks by mapping the 497 near-specific host interactors that associated with less than five effector families (Figure 2B). This sub-network revealed that edges from host proteins involved in some of the plant processes converged on a particular effector family. For example, the edges from plant proteins involved in ‘small molecule transport’ and ‘vesicle-mediated transport’ processes converged on the PexRD12/31 family of effectors; the edges from proteins assigned to the ‘protein import to nucleus’ process converged on the AVRvnt1 family; and the edges from proteins in ‘signal transduction’ and ‘proteolysis’ processes converged on the AVR2 family (Figure 2B, Table S2). To visualize these connections, we generated subnetworks for selected processes (Figure S3). These subnetworks revealed more detailed associations between individual host proteins involved in the biological process and effector families. For example, 24 out of the 32 members of the ‘vesicle mediated transport’ process associated with the PexRD12/31 family, and seven out of nine members of the ‘protein import to nucleus’ associated with the AVRvnt1 family. The observation that multiple proteins assigned to the same host biological process associated predominantly with single effector families suggests that these host processes constitute targets of these effectors.

We also generated an interaction subnetwork for the 16 most promiscuous interactors that showed association with more than nine effector families (Figure 2C). This subnetwork revealed that the promiscuous host interactors are distributed across different biological processes with edges radiating towards different effector families in an indiscriminate manner, suggesting that these interactions are non-specific. Therefore, we conclude that the GO-based interactome network analysis was useful in flagging host processes likely targeted by *P. infestans* RXLR effectors as well as identifying potential non-specific interactions.

### PexRD12/31 proteins form a complex family of *Phytophthora infestans* RXLR-WY effectors

Given the prominence of vesicle mediated trafficking as a target of the PexRD12/31 family and considering that this large effector family has not been previously functionally characterized, we decided to further characterize this family. We reasoned that PexRD12/31 effectors could serve as useful probes to shed further light on how *P. infestans* manipulates host vesicular trafficking. The PexRD12/31 family comprises 19 predicted members in the genome of *P. infestans* strain T30-4 (Haas *et al.*, 2009). Two family members, PexRD12 (PITG_16245) and PexRD31 (PITG_23074), were previously reported in *P. infestans* isolate 88069 and included in *in planta* screens for AVR activities (Vleeshouwers *et al.*, 2008; Oh *et al.*, 2009). PexRD12 is recognized by *Rpi-chc1,* a *Solanum chacoense* resistance gene against *P. infestans* (Vossen *et al.*, 2017). Sequence analyses of the 19 predicted genes revealed that two members (PITG_23230 and PITG_20336) lack the effector domain and one divergent member (PITG_09577) lacks the signature RXLR-EER motif. To evaluate the sequence diversity of the remaining 16 family members, we compared the amino acid sequence of their effector domains (Figure 3). Phylogenetic analyses grouped 15 of the 16 members into four clades supported by bootstrap values over 90%, while one divergent member (PITG_16428) contained a 10 amino acid insertion in the effector domain and did not cluster with any of the other proteins (Figure 3). The effector domains of the canonical family members comprise 62 amino acids and all 16 proteins contained a single predicted WY-fold based on HMMER searches (E value < 1.8^-07^) (Boutemy *et al.*, 2011). Proteins within each of the four clades displayed an average pairwise amino acid identity of over 95% in their effector domains. We conclude that the PexRD12/31 family of *P. infestans* RXLR-WY effectors is structured into four clades of highly similar proteins with PITG_16428 as a fifth divergent member of the family.

**Figure 3.**
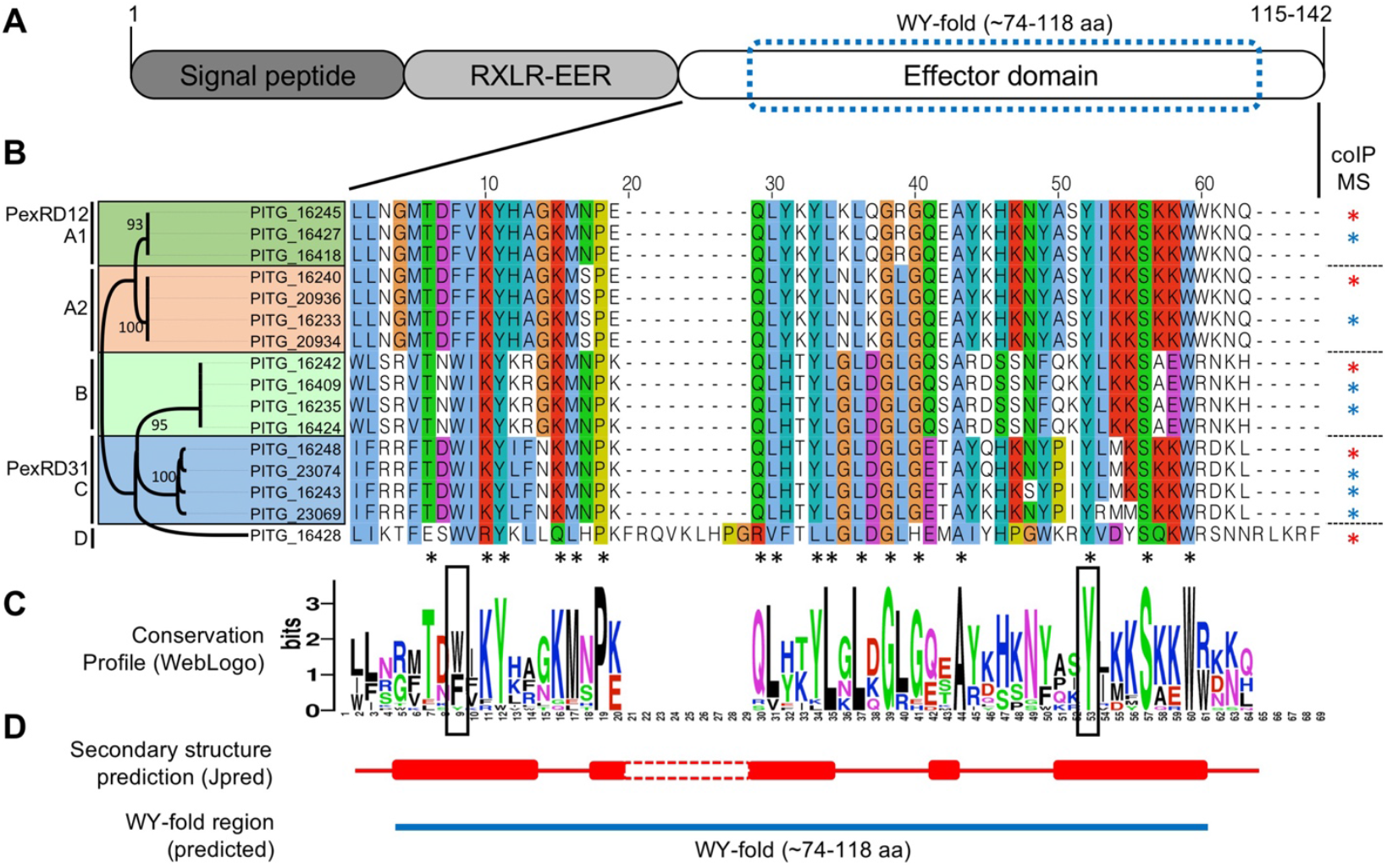
*Phytophthora infestans* PexRD12/31 family of RXLR effectors group into four distinct classes. **(A)** Schematic diagram of the canonical protein domain organization of PexRD12/31 proteins. Numbers indicate minimal and maximal peptide length of the effectors. **(B)** Phylogenetic tree and amino acid alignment of the C-terminal domain of the 16 members of the PexRD12/31 family. The family consists of four distinct groups (A1, A2, B and C) and PITG_16428 on its own in D. The phylogenetic relationship within the PexRD12/31 family members was inferred by using the Maximum Likelihood method and JTT matrix-based model implemented in MEGA X. Simplified version of the tree with the highest log likelihood (−561.03) is shown. The percentage of trees in which the associated taxa clustered together is shown next to the branches. All positions containing gaps and missing data were eliminated. There was a total of 54 positions in the final dataset. Amino acid residues are colored according to the ClustalX scheme. The black asterisks indicate the amino acid residues that are conserved in ≥ 90% of the sequences. The blue asterisks indicate effectors used for the first coIP/MS screen; the red asterisks indicate effectors used for both the coIP/MS screen and follow-up experiments. **(C)** WebLogo conservation profile of the sequence alignment shown in B. **(D)** Jpred4-predicted alpha helices and predicted WY-fold in PexRD12/31 family members. Solid red cylinders indicate predicted helices based on the sequences of PITG_16424 and PITG_23074. The predicted WY-fold region is indicated by a blue bar.

### PexRD12/31 effectors consistently associate with vesicle trafficking-related host proteins in independent coimmunoprecipitation/mass spectrometry experiments

To further explore the association between PexRD12/31 effectors and host proteins implicated in vesicle trafficking, we fused the effectors to green fluorescent protein (GFP) and used them in additional coIP/MS experiments. We selected one PexRD12/31 family representative from each of the four clades: PexRD12, PexRD31, PITG_16242, and PITG_16428 (Figure 3). We expressed the GFP effector fusion proteins in *N. benthamiana* leaf cells by agroinfiltration and performed coIP/MS using anti-GFP antibodies under the stringent conditions described in the methods (Dataset 3). All the fusion proteins could be detected after coIP by either SDS-PAGE Coomassie blue staining, mass spectrometry, or both, indicating effective expression and immunoprecipitation (Figure S4). These assays yielded 24 out of the 30 vesicle trafficking-related host proteins that we previously identified in the anti-FLAG coIP/MS screen (Table S3). Among these 24 proteins, 16 associated with no more than four bait proteins throughout our laboratory dataset of over a hundred independent coIP/MS assays (Dataset 1, Dataset 2) (Petre *et al.*, 2015; Petre *et al.*, 2016). We conclude that these 16 *N. benthamiana* proteins are relatively-specific interactors of PexRD12/31 effectors in coIP/MS assays.

### PexRD12/31 effectors associate with *Nicotiana benthamiana* R-SNARE protein of the VAMP72 family

Vesicle associated membrane proteins (VAMPs, also known as R-SNAREs) are components of secretory vesicles and endosomes that mediate vesicle fusion. VAMPs have been implicated in plant pathogen response (Collins *et al.*, 2003; Kwon *et al.*, 2008; Yun and Kwon, 2017) but are not known to be targeted or associated with pathogen effectors. Our FLAG coIP/MS assays identified a protein annotated as vesicle associated membrane protein (VAMP) 7B (Sol Genomics Network (SGN) accession NbS00022342g0004) by two peptide hits in Mascot searches from IP of effector PITG_16428, a member of PexRD12/31 family (Figure S5). In addition, other PexRD12/31 effectors were also shown to consistently and specifically associated with VAMP 7B in the GFP coIP/MS assays (Table S3). VAMPs occur as multiple paralogs in *N. benthamiana* (Figure S5). To further characterize the sequence of NbS00022342g0004 we performed BLASTP (BLAST 2.9.0+) (Altschul *et al.*, 1997) searches against the *A. thaliana* proteome version Araport11 in which VAMPs are comprehensively annotated (The Arabidopsis Information Resource, https://www.arabidopsis.org). The top hit was AT1G04760, annotated as AtVAMP726, with an E value of 2^-126^ and a score of 357. To determine the relationship between NbS00022342g0004 and *A. thaliana* VAMPs, we extracted all members of AtVAMP72 family (AtVAMP721-727) from Araport11 and 14 VAMP72-like amino acid sequences from the proteome of *N. benthamiana* (Sol Genomics Network, version 0.44). Phylogenetic analyses of these sequences revealed that NbS00022342g0004 does not have a clear ortholog in *A. thaliana* within the VAMP72-family (Figure S5) and therefore we decided to refer to it from here on as NbVAMP72x.

To further evaluate whether PexRD12/31 effectors physically associate with NbVAMP72x *in planta*, we combined coIP and immunoblot assays. We co-expressed eight FLAG tagged PexRD12/31 effectors with GFP-NbVAMP72x in *N. benthamiana* leaf cells and performed immunoprecipitations with anti-FLAG and anti-GFP antibodies bound to agarose beads. These coIP assays revealed that seven tested PexRD12/31 effectors associated with NbVAMP72x *in planta* (Figure 4). One effector, PITG_16427, did not associate with NbVAMP72x. However, PITG_16427 did not accumulate to detectable levels in the input samples indicating that the protein is not stable *in planta* and making the IP assay inconclusive (Figure 4). The negative control FLAG-AVR3a, which did not associate with NbVAMP72x in the coIP/MS screens described earlier, also did not associate with NbVAMP72x in the coIP/immunoblot experiments. We conclude that members of the PexRD12/31 family associate with NbVAMP72x *in planta*.

**Figure 4.**
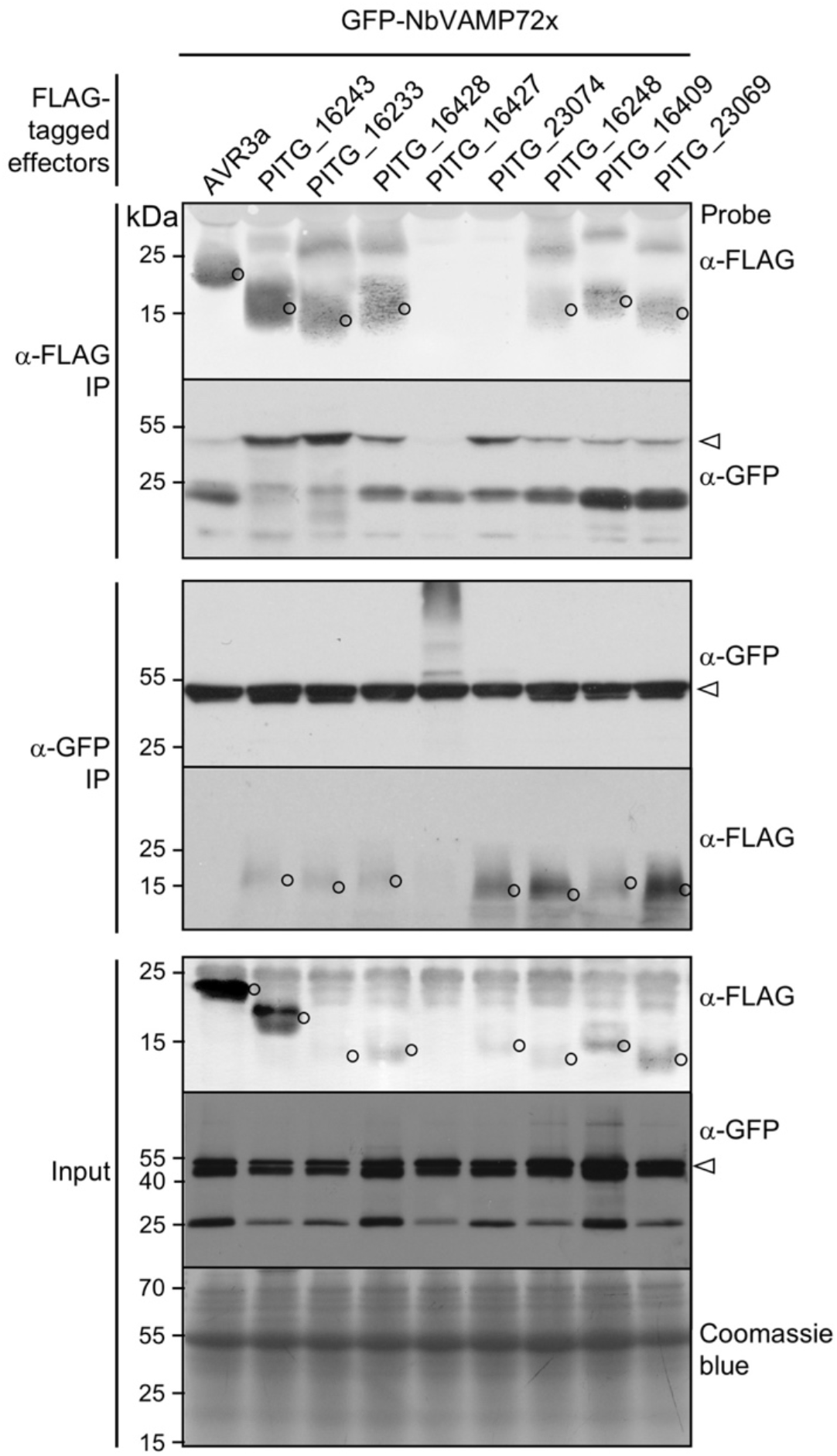
PexRD12/31 effectors associate with NbVAMP72x *in planta.* Immunoblots show FLAG-tagged PexRD12/31 family effector fusion proteins coimmunoprecipitate with GFP-tagged NbVAMP72x transiently co-expressed in *N. benthamiana.* Approximate molecular weights of the proteins are shown on the left. Open arrowhead shows the expected size of the NbVAMP72x bands and open circles show the expected sizes of the effector bands. IP = immunoprecipitation.

### PexRD12/31 effectors mainly accumulate at the host plasma membrane

To determine the subcellular localization of PexRD12/31 effectors in plant cells, we expressed the GFP N-terminally tagged PexRD12, PexRD31, PITG16242, and PITG16428 described above in *N. benthamiana* leaf cells by agroinfiltration and performed live cell imaging by confocal microscopy. GFP-tagged PexRD31, PITG16242, and PITG16428 produced informative fluorescent signals that distributed mainly at the cell periphery (Figure S6). In addition, the GFP-PexRD31 fluorescent signal accumulated sharply around the nucleus and in small and mobile puncta. GFP-PexRD12 was not informative as it produced a weak fluorescent signal and triggered a necrotic response in leaves that interfered with the imaging.

To evaluate the robustness of these observations, we performed similar live cell imaging experiments with effectors N-terminally tagged with the fluorescent protein mCherry (Figure S6). We observed similar patterns for PexRD31, PITG16242, and PITG16428 with the fluorescence signal accumulating mainly at the cell periphery. Here too, mCherry-PexRD12 triggered a necrotic response with no clear fluorescent signal. Interestingly, as observed with the GFP fusion, mCherry-PexRD31 labelled the nuclear periphery and mobile cytosolic puncta in addition to the cell periphery, and we documented these mobile signals in a movie (Movie S1).

To determine whether PexRD31, PITG16242, and PITG16428 accumulate at the plasma membrane, we co-expressed the mCherry fusion proteins with the plasma membrane marker protein YFP-Rem1.3 in *N. benthamiana* using agroinfiltration (Bozkurt *et al.*, 2015) and contrasted them to co-expression with cytosolic free GFP (Figure 5). All three effectors produced a sharp fluorescent signal on the outside edge of the cytosol that overlapped with the YFP-Rem1.3 fluorescent signal and showed no detectable background in the cytosol (Figure 5).

**Figure 5.**
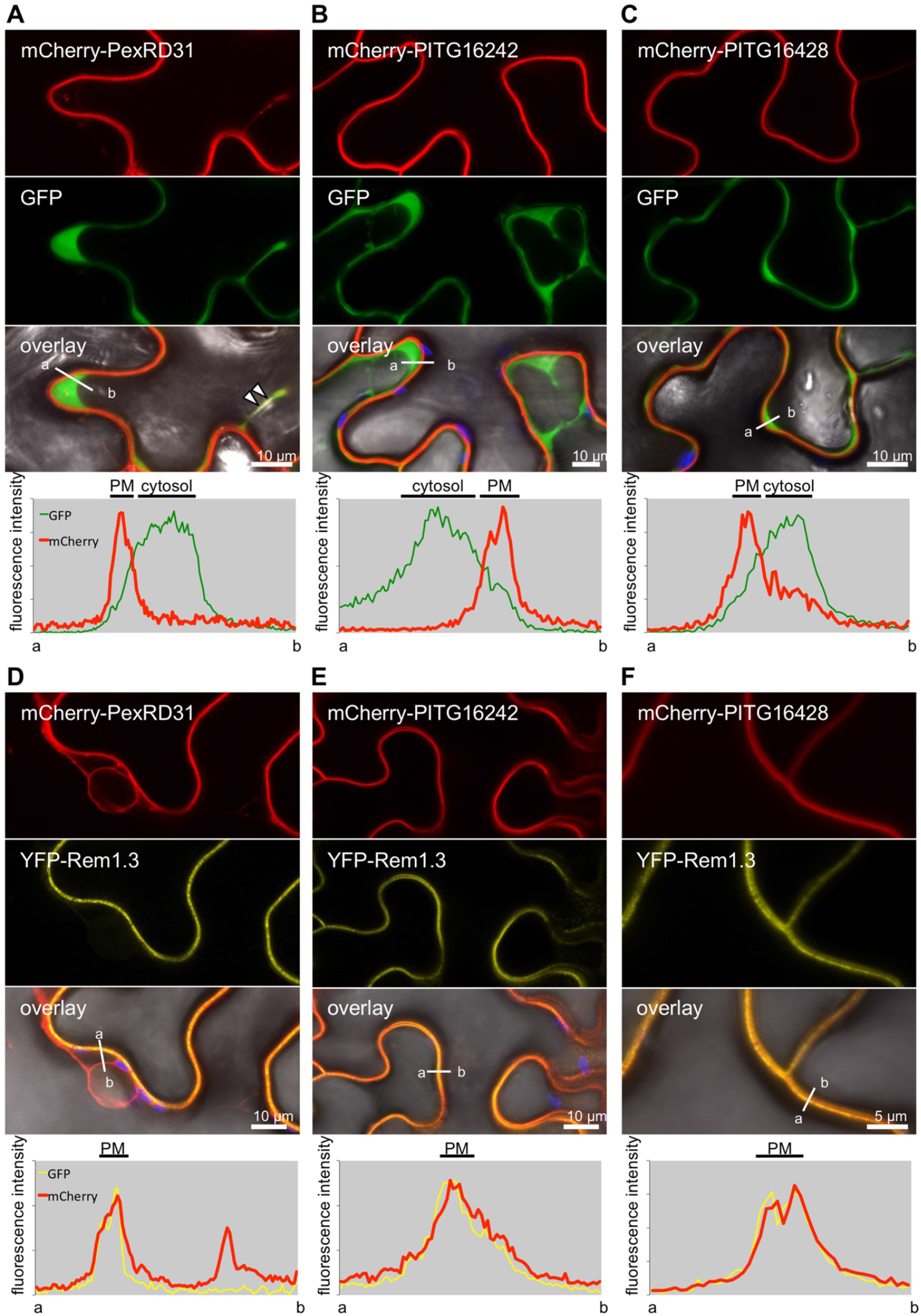
PexRD12/31 effectors accumulate at the host plasma membrane. mCherry-PexRD31, mCherry-PITG16242 and mCherry-PITG16428 were co-expressed with free green fluorescent protein (GFP) or with the plasma membrane (PM) marker YFP-Rem1.3 via agroinfiltration in *N. benthamiana* leaves. Live-cell imaging was performed with a laser-scanning confocal microscope three days after infiltration. **(A-F)** Confocal microscopy of *N. benthamiana* leaf epidermal cells co-expressing mCherry-PexRD31, mCherry-PITG16242 or mCherry-PITG16428 with free GFP (A-C) or PM marker YFP-Rem1.3 (D-F). All three effectors show PM localization, while mCherry-PexRD31 also accumulated in cytosolic puncta (white arrowheads). Images are single optical sections of 0.8 μm. The overlay panel combines GFP (A-C) or YFP (D-F), mCherry, chlorophyll, and bright field images. The lower panels show relative fluorescence intensity plots of the GFP (A-C, green sharp line) or YFP (D-F, yellow sharp line) and the mCherry (red thick line) signals along the line from a to b depicted in the corresponding overlay panels.

We conclude that the three effectors accumulate at the plasma membrane when expressed in *N. benthamiana* cells. We also noted that the mCherry-PexRD31 mobile puncta reported above overlapped with the cytosolic GFP signal. In summary, we conclude that all three effectors accumulate at the plasma membrane, and that PexRD31 also localizes to mobile cytosolic bodies.

### PexRD31 localizes to RabC1-positive mobile vesicles of the post-Golgi/endosomal network

We further investigated the localization of PexRD31 cytosolic mobile puncta by co-expressing its GFP or mCherry fusions with a set of eleven fluorescent proteins that mark different plant cell endomembrane compartments and organelles. These include secretory vesicles (NbVAMP72x-mRFP), peroxysomes (GFP-PTS1), Golgi (MAN11-49-GFP), phosphatidylinositol-3-phospate (PI3P)-positive vesicles (2xFYVE-GFP), endoplasmic reticulum (ER, WAK2_SP_-GFP-HDEL), autophagosomes (GFP-ATG8C), early and late endosomes (mRFP-ARA7), late endosomes and multi vesicular bodies (ARA6-mRFP), exocyst-positive organelle (EXPO, Exo70E2-GFP), mitochondria (COX41-29-GFP), and post-Golgi/endosomal network (YFP-RabC1) (Nelson *et al.*, 2007; Voigt, 2008; Heard *et al.*, 2015; Dagdas *et al.*, 2016). Among these, we noted a distinct overlap in the fluorescent signals produced by mCherry-PexRD31 and YFP-RabC1 (Figure 6). These overlapping signals marked mobile puncta that are probably cytosolic vesicles associated with post-Golgi endosomal elements (Geldner *et al*., 2009). These observations were specific to PexRD31 since another member of the PexRD12/31 family, PITG16242, did not co-localize with YFP-RabC1 (Figure 6C). The colocalization between PexRD31 and RabC1 was also specific in our experiments, as PexRD31 fluorescent signals did not overlap with any of the other ten markers (Figure S7). Notably, aside from colocalization with RabC1, PexRD31 signals did not overlap with other markers associated with the post-Golgi endosomal network, such as early and late endosomes, secretory vesicles and Golgi bodies (Figure S7). We conclude that PexRD31 specifically localizes to RabC1-positive mobile vesicles and that this localization is specific to PexRD31.

**Figure 6.**
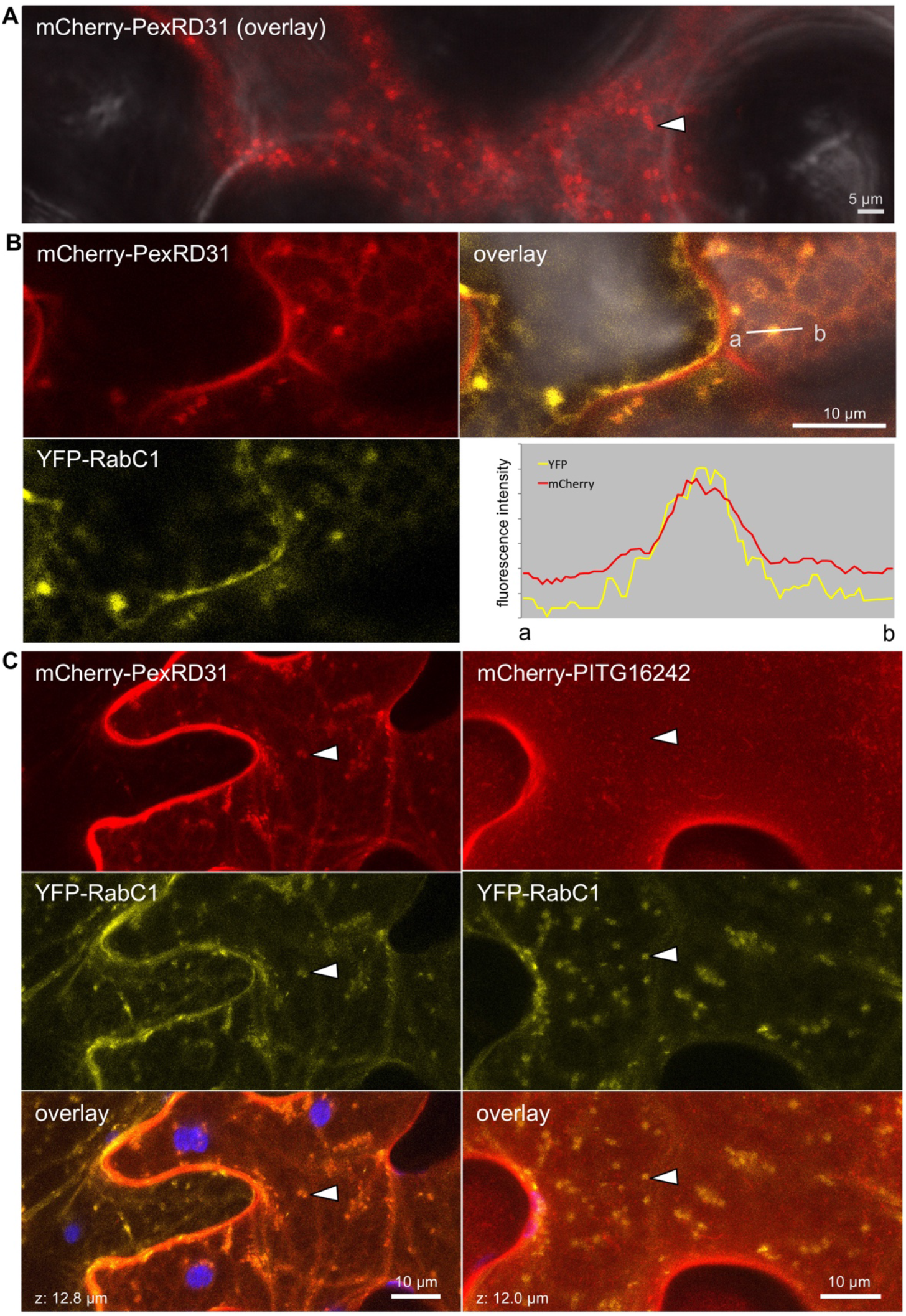
PexRD31 accumulates at RabC1-positive mobile vesicles. **(A)** Confocal microscopy of *N. benthamiana* leaf epidermal cells expressing mCherry-PexRD31. The panel shows an overlay of the mCherry and bright field channels. It corresponds to the first image of Movie S1. **(B)** Confocal microscopy of *N. benthamiana* leaf epidermal cells co-expressing mCherry-PexRD31 and YFP-RabC1. The right-hand side panel shows relative fluorescence intensity plots of the YFP and the mCherry along the line from a to b depicted in the corresponding overlay panel. **(C)** Confocal microscopy of *N. benthamiana* leaf epidermal cells co-expressing mCherry-PexRD31 or mCherry-PITG16242 with YFP-RabC1. In all cases proteins were expressed in leaf cells by agroinfiltration. Live-cell imaging was performed with a laser-scanning confocal microscope three days after infiltration. Images are single optical sections of 0.8 μm or maximal projections of 10 optical sections (max. z-stack: 12.8 μm). The overlay panels combine YFP, mCherry, and chlorophyll channels.

### PexRD12/31 effectors accumulate at haustoria in *Phytophthora infestans-* infected *Nicotiana benthamiana* cells

We previously reported that several endomembrane compartments are directed towards *P. infestans* haustoria during infection (Bozkurt *et al.*, 2012; Bozkurt *et al.*, 2015; Dagdas *et al.*, 2018; Bozkurt and Kamoun, 2020). In addition, some effectors display perihaustorial localization accumulating around haustoria in infected plant cells (Bozkurt *et al.*, 2011; Saunders *et al.*, 2012; Dagdas *et al.*, 2018; Wang *et al.*, 2018). To investigate whether this applies to the PexRD12/31 family, we determined the subcellular localization of the mCherry fusions of PexRD31, PITG16242, and PITG16428 in haustoriated cells of *N. benthamiana* (Figure 7) using established protocols and methods (Bozkurt *et al.*, 2011). All three proteins produced sharp fluorescent signals around haustoria indicating that they are perihaustorial effectors and accumulate at the haustorial interface when expressed in infected haustoriated plant cells (100% of haustoria imaged, N>60) (Figure 7A-C).

**Figure 7.**
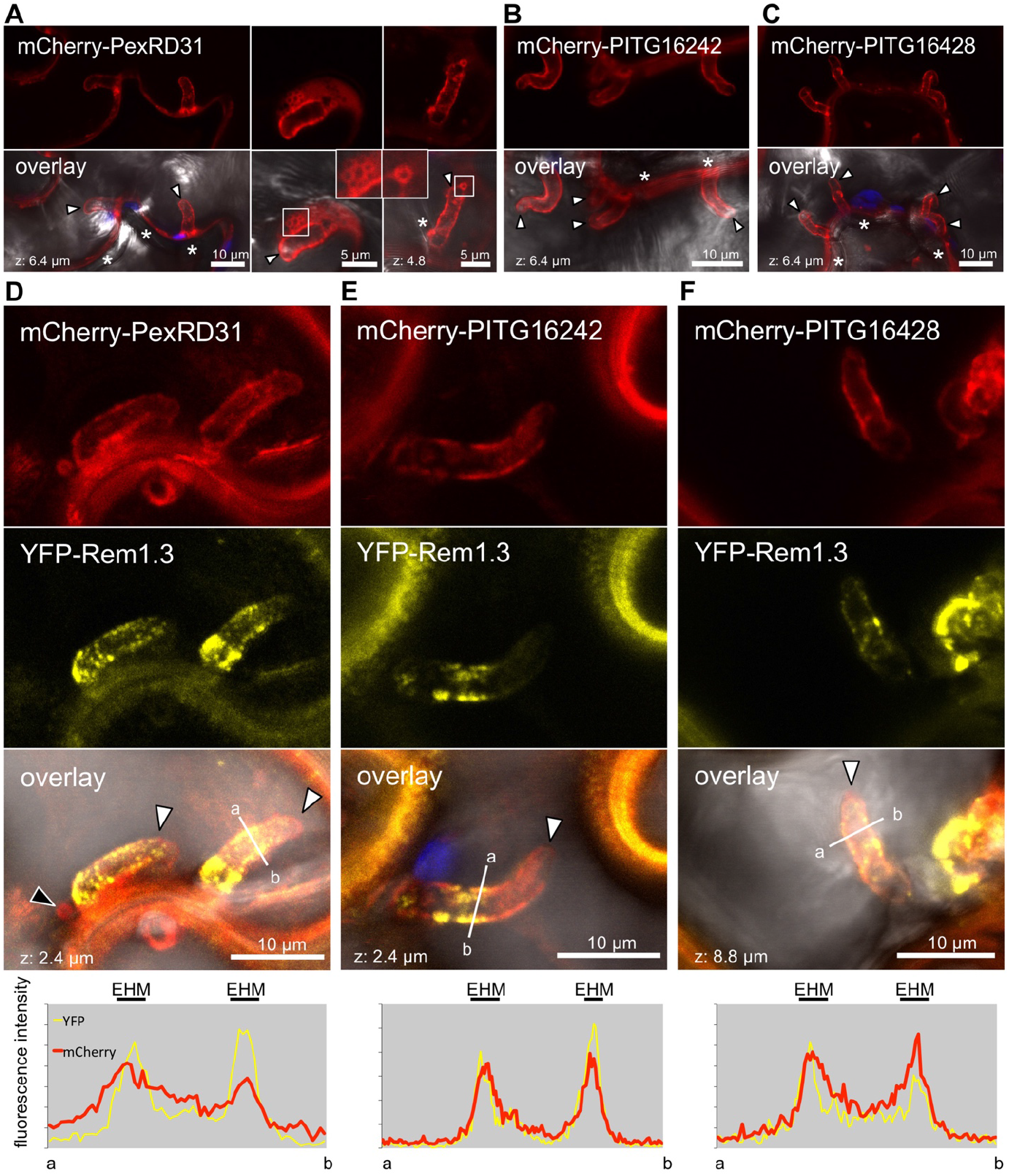
PexRD12/31 effectors accumulate around haustoria. mCherry-PexRD31, mCherry-PITG16242 or mCherry-PITG16428 were expressed on their own or co-expressed with YFP-Rem1.3 in leaf cells by agroinfiltration, and leaves were drop inoculated with zoospores of *P. infestans* isolate 88069 three hours after agroinfiltration. Live-cell imaging was performed with a laser-scanning confocal microscope three days after infection. **(A)** Confocal microscopy of *P. infestans* infected *N. benthamiana* leaf epidermal cells expressing mCherry-PexRD31. mCherry-PexRD31 around haustoria and at perihaustorial vesicles. The two panels on the right-hand side show mCherry-labelled vesicles in contact with haustoria. The inserts in the overlay panels show a close-up of these vesicles. **(B and C)** Confocal microscopy of *P. infestans* infected *N. benthamiana* leaf epidermal cells expressing mCherry-PITG16242 and mCherry-PITG16428, respectively. Both effectors accumulated around haustoria. **(D)** Confocal microscopy of *P. infestans* infected *N. benthamiana* leaf epidermal cells co-expressing mCherry-PexRD31 and PM and EHM marker YFP-Rem1.3. The black arrowhead indicates a PexRD31-labelled vesicle in close proximity to the leaf haustorium. **(E and F)** Confocal microscopy of *P. infestans* infected *N. benthamiana* leaf epidermal cells co-expressing mCherry-PITG16242 and mCherry-PITG16428 (respectively) with PM and EHM marker YFP-Rem1.3. Images are single optical sections of 0.8 μm or maximal projections of 11 optical sections (max. z-stack of 8.8 μm). The overlay panels combine YFP (D-F), mCherry, chlorophyll, and bright field images. White arrowheads indicate haustoria. For (A-C), white asterisks indicate extracellular hyphae, which can be out of focus. For (D-F), lower panel show relative fluorescence intensity plots of the YFP and the mCherry along the line from a to b depicted in the corresponding overlay panel.

In plants infected with *Phytophthora,* the host side of the haustorial interface is comprised of the extrahaustorial membrane (EHM), a thin layer of cytosol and the tonoplast that tends to be in close proximity to the EHM (Bozkurt *et al.*, 2012). To further determine where the PexRD12/31 effectors accumulate, we co-expressed in *N. benthamiana* their mCherry fusions with YFP-Rem1.3, a marker that sharply labels EHM microdomains (Bozkurt *et al.*, 2014). The fluorescent signal produced by the three effectors overlapped with the YFP-Rem1.3 signal indicating that these effectors accumulate at the EHM (Figure 7D-F).

Experiments with *P. infestans*-infected *N. benthamiana* tissue enabled us to visualize the mCherry-PexRD31 puncta in haustoriated cells. Some of these puncta were immobile and were in proximity to the haustorial interface, enabling us to image them at higher resolution. This revealed that mCherry-PexRD31 produced a sharp circular fluorescent signal that can be observed near haustoria (N>20) (Figure 7). We conclude that PexRD31 accumulates at vesicle-like structures and probably localizes on the outside of these vesicles. Consistent with our previous observations with non-infected *N. benthamiana* cells (Figure 5, Figure 6), PITG16242 and PITG16428 differed from PexRD31 and did not label vesicle-like structures. Altogether, these data indicate that PexRD31, PITG16242, and PITG16428 accumulate at the EHM around *P. infestans* haustoria. In addition, PexRD31 accumulates in vesicle-like structures in proximity to the haustorial interface.

### PexRD31 increases the number of FYVE-labelled endosomes in *Nicotiana benthamiana* cells

During co-expression experiments with the FYVE marker (2xFYVE-GFP), we noted that PexRD31 appears to increase the number of GFP-labelled endosomes. We further investigated this observation by using agroinfiltration to express the mCherry fusions to PexRD31, PITG16242, and PITG16428 in leaves of transgenic *N. benthamiana* expressing 2xFYVE-GFP (Figure 8A-B). In independent experiments, mCherry-PexRD31 increased the number of FYVE-labelled endosomes by about 40% compared to mCherry and the other two effectors (n=40).

**Figure 8.**
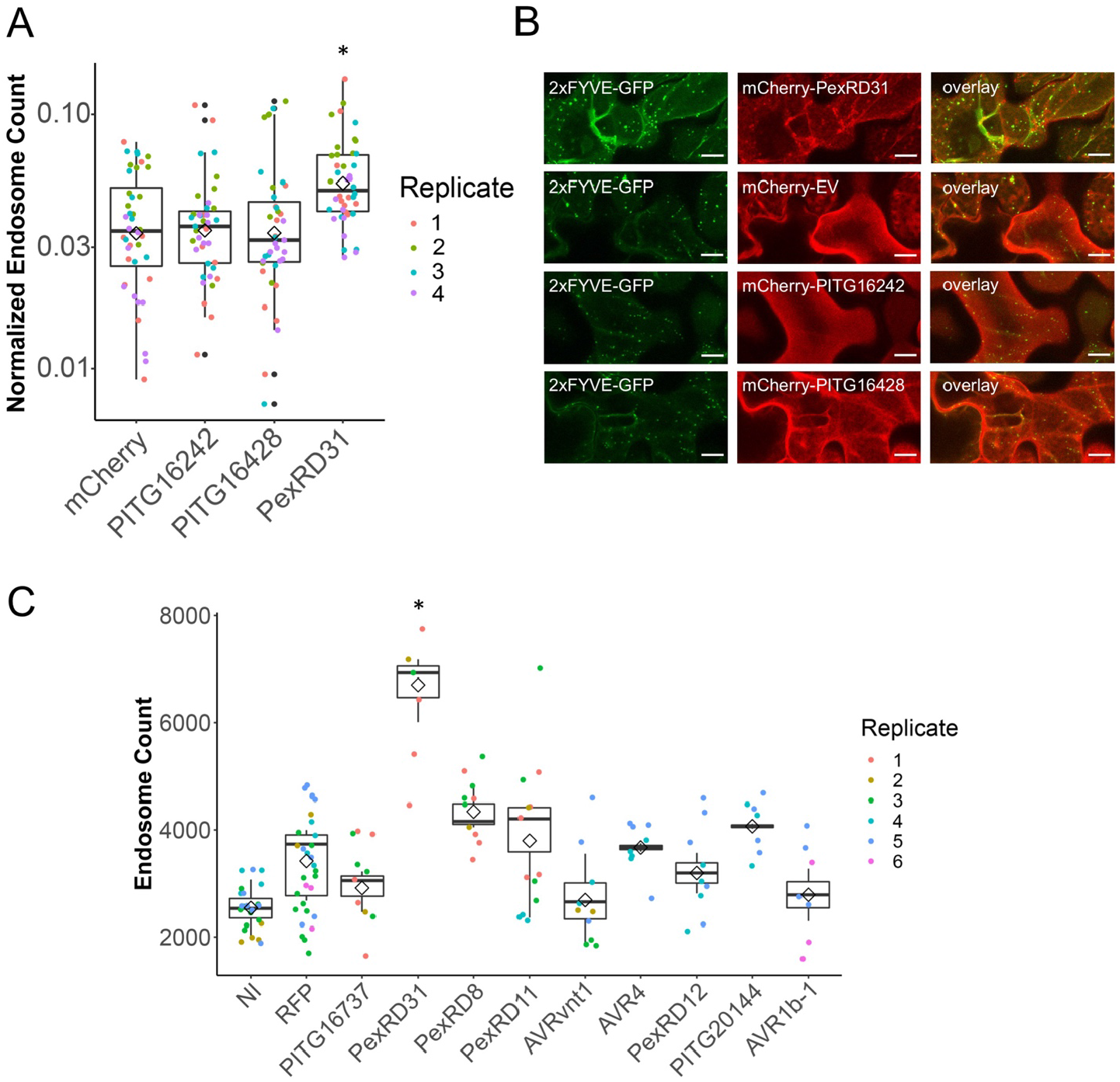
PexRD31 increases the number of FYVE labelled endosomes in *Nicotiana benthamiana* cells. **(A)** Categorical scatterplots with superimposed boxplots displaying the number of 2xFYVE-GFP endosomes per μm^2^ of cytoplasm in cells expressing mCherry-PexRD31, mCherry-PITG_16242, mCherry-PITG16428, or mCherry. The number of 2xFYVE-GFP-labelled endosomes was significantly enhanced by the expression of mCherry-PexRD31 (* = p<0.01). The data are representative of 40 maximum projection images from 4 biological replicates. Each replicate consists of 10 individual Z-stacks of 17 slices per treatment. Images were taken 3 days post infiltration. **(B)** Representative images of transient expression of mCherry-PexRD31, mCherry-EV, mCherry-PITG_16242 and mCherry-PITG_16428 in leaf epidermal cells of transgenic *N. benthamiana* lines expressing 2xFYVE-GFP. Scale bar = 10μm. **(C)** Categorical scatterplots with super-imposed boxplots displaying the number of 2xFYVE-GFP endosomes per field of view in the presence of different FLAG-tagged effectors, RFP, or with no agroinfiltration (NI). 2xFYVE-GFP endosomes were significantly enhanced by the expression of PexRD31 (* = p<0.01) with respect to the NI and RFP controls and the other effectors assayed. None of the other effectors assayed significantly increased the number of 2xFYVE-GFP-labelled endosome numbers. We obtained 2, 3 or 4 biological replicates per treatment, with each replicate consisting of 4 individual z-stacks of 12 slices each per treatment. The data consist of averaged counts obtained from individual slices conforming each z-stack. Images were acquired 3 days post-infiltration.

We also compared PexRD31 with nine *Phytophthora* RXLR effectors selected from our collection, which allowed to estimate the degree to which the observed activity is specific. These assays were done with a different expression vector system and experimental setup to determine the robustness of the effect. We performed agroinfiltration to express the FLAG tagged effector constructs used earlier in the coIP experiments in the 2xFYVE-GFP transgenic *N. benthamiana* plants. PexRD31 significantly increased the number of 2xFYVE-GFP labelled endosomes, resulting in an increase of 40 to 90% compared to the all other effectors and 90% compared to the negative control (Figure 8C). PexRD31 was the only effector to show a statistically significant increase with respect to the RFP control. We conclude that PexRD31 specifically alters vesicle trafficking by increasing the number of PI3P endosomes labelled by the FYVE marker.

### FYVE-labelled endosomes accumulate in *Nicotiana benthamiana* tissue colonized by *Phytophthora infestans*

To determine whether the effect of PexRD31 on FYVE-labelled endosomes can be recapitulated with the pathogen, we examined the number and distribution of FYVE-labelled endosomes during *P. infestans* infection of *N. benthamiana* leaves. To this end, we inoculated *N. benthamiana* plants expressing 2xFYVE-GFP with *P. infestans* 88069td, a transgenic strain expressing the red fluorescent marker tandem dimer RFP (tdTomato) (Whisson *et al.*, 2007; Giannakopoulou *et al.*, 2014). We evaluated 2xFYVE-GFP fluorescence signals relative to the RFP fluorescence and focused our imaging on the edge of the disease lesions where haustoria are produced and which corresponds to the biotrophic phase of infection (van West *et al.*, 1998; Lee and Rose, 2010). Using low magnification confocal microscopy, we noted that leaf areas biotrophically colonized by *P. infestans* display bright and large GFP puncta; such puncta were absent in non-colonized areas (Figure 9A). *N. benthamiana* cells with GFP puncta signals were in direct contact with *P. infestans* hyphae, whereas the non-colonized areas surrounding the lesions showed hardly any GFP puncta signals. 2xFYVE-GFP puncta were mobile, with a maximal diameter of 2μm (Figure 9B, Movie S2). We compared these patterns to *P. infestans* infections of *N. benthamiana* stable transgenics expressing free GFP (marking the nucleus and the cytosol) or a YFP-RabG3c fusion (marking late endosomes and the tonoplast). In both cases, there was no evident alteration of the distribution of the fluorescence signals in tissue colonized by *P. infestans* (Figure S8), indicating that *P. infestans* specifically perturbs 2xFYVE-GFP-positive compartments during the biotrophic colonization of *N. benthamiana.*

**Figure 9.**
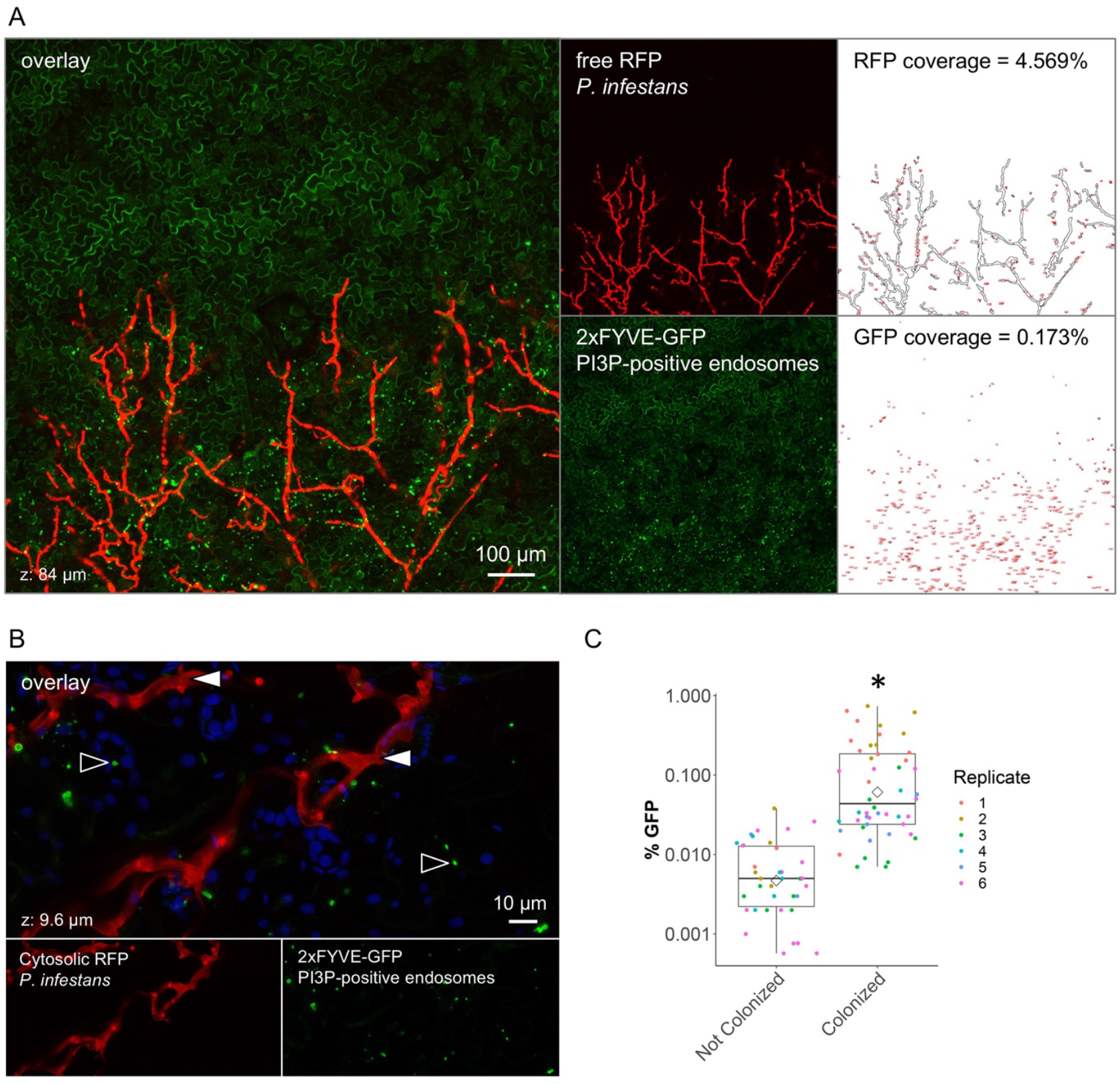
FYVE-labelled endosomes accumulate in leaf cells colonized by *Phytophthora infestans.* **(A)** Low magnification live cell imaging of a 2xFYVE-GFP fusion (marker of PI3P-positive endosomes) in *N. benthamiana* leaf cells colonized by *P. infestans* isolate 88069td. The right-hand side panels show processed images used for the quantification of the image area displaying an RFP (upper panel) or a GFP puncta (lower panel) fluorescent signal, expressed as a percentage of the total image area. Images are maximal projections of 21 optical sections (z-stack of 84 μm). **(B)** High magnification live cell imaging of a 2xFYVE-GFP fusion in *N. benthamiana* leaf cells colonized by *P. infestans* isolate 88069td. In the overlay image, black-lined white arrows indicate *P. infestans,* and white-lined black arrows indicate FYVE-labelled puncta. Images are maximal projections of 12 optical sections (z-stack of 9.6 μm). **(C)** Categorical scatterplots with superimposed boxplots showing the semi-automated quantification of GFP puncta coverage from confocal microscopy images corresponding to leaf areas biotrophically colonized by *P. infestans* (‘Colonized’) or leaf areas without *P. infestans* (Not Colonized). (* = p<0.01) Leaves of stable transgenic *N. benthamiana* plants were drop inoculated by zoospores of *P. infestans* isolate 88069td. Live-cell imaging was performed with a laser-scanning confocal microscope three days after inoculation. The overlay panels combine (A) GFP and RFP channels, or (B) GFP, RFP, and chlorophyll channels.

To further evaluate the correlation between the presence of *P. infestans* hyphae and the formation of FYVE-labelled puncta, we performed a blind confocal microscopy image acquisition analysis. We acquired 13 z-stacks of 11 images over leaf tissues colonized by *P. infestans* 88069td and surrounding non-colonized areas (see methods). Next, we developed and applied an automatic analytical pipeline using the ImageJ package Fiji (Schindelin *et al.*, 2012) to quantify *P. infestans* red fluorescence and punctate GFP-FYVE fluorescence signals. This experiment further showed that 2xFYVE-GFP-positive puncta signals increased in *N. benthamiana* tissue during the biotrophic stage of *P. infestans* infection compared to the non-colonized areas of leaf tissue (Figure 9C). This set of experiments indicate that *P. infestans* alters host endosome trafficking during biotrophic colonization.

## DISCUSSION

A major research aim in the field of molecular plant pathology is to unravel the activities of effectors in order to understand how pathogens successfully colonize their hosts. The rationale behind this project is based on the view that effectors can serve as molecular probes to identify the host processes that pertain to pathogen interactions. We applied an effectoromics pipeline centered on an *in vivo* proteomics-based protein-protein interaction screen. This screen revealed that ~50 *P. infestans* RXLR effectors associate *in planta* with ~580 unique *N. benthamiana* proteins representing as many as 35 biological processes. We conclude that *P. infestans* RXLR effectors target multiple processes in their host plants; and our study provides a broad overview of these effector-targeted processes.

We previously reported *in planta* proteomics screens for interactors of candidate effectors of the rust fungal pathogens *Melampsora larici-populina* and *Puccinia striiformis* f sp *tritici* (Petre *et al.*, 2015; Petre *et al.*, 2016). Our study also complements large-scale yeast-two-hybrid screens for effector interactors of the oomycete plant pathogen *Hyaloperonospora arabidopsidis,* the bacterium *Pseudomonas syringae* and the powdery mildew fungus *Golovinomyces orontii* (Mukhtar *et al.*, 2011; Weßling *et al.*, 2014). As previously discussed (Petre *et al.*, 2015; Petre *et al.*, 2016), *in planta* protein-protein interaction assays have both advantages and disadvantages compared to the more commonly used yeast-two hybrid assay. Given that the coIP assay takes place *in vivo* in plant tissue, the host proteins are expressed in a biologically relevant molecular context and cellular compartment. Another difference is that yeast two-hybrid interactions are presumably binary, detecting one-on-one protein interactions, whereas the proteins we identified by coIP/MS may not directly bind the effector but could instead associate in a multi-protein complex. Indeed, our observation that functionally related plant proteins tend to associate with a given effector family increases the probability that the targeted host complex or process is physiologically relevant (Figure 2).

One drawback of the coIP method is that a direct host interactor may be missed even when relevant associated proteins are recovered. This could be due to several factors, such as abundance of the interacting plant proteins, absence of a suitable tryptic peptide detectable by mass spectrometry, or absence of the protein sequence in the annotated database used for the mass spectra searches. Also, given that mass spectrometry sampling of peptides is random, a target protein may be missed by chance. One example is the *P. infestans* RXLR-WY effector PexRD54, which binds with high affinity to the autophagy protein ATG8CL (Dagdas *et al.*, 2016; Maqbool *et al.*, 2016). Although ATG8CL was missing from the initial coIP/MS screen it was picked up in two out of four subsequent experiments (Dagdas *et al.*, 2016). This might be due to the low molecular weight of ATG8CL (~14 kDa), which may not have produced enough peptides for MS detection. Interestingly, we flagged ATG8C as likely to directly bind PexRD54 because their interaction survived stringent binding conditions in contrast to other candidate interactors (Dagdas *et al.*, 2016). Rab8a, another interactor identified in the original PexRD54 coIP/MS experiment, was recently reported as a genuine target of PexRD54 that associates with this effector independently of ATG8CL binding (Pandey *et al.*, 2020). The PexRD54 work illustrates the importance of replication and follow-up experiments to the coIP/MS. As with any biochemical assay, there is a benefit to experiment with different assay conditions, such as binding stringency.

A subset of the host proteins we identified can be tagged as “usual suspects” that are unlikely to form biologically relevant interactions with the effectors (Petre *et al.*, 2015; Petre *et al.*, 2016). Nonetheless, 497 of the host interactors associated with no more than 4 out of the 14 effector families (Dataset 2). These relatively specific interactors are more likely to be biologically relevant compared to the more promiscuous proteins, which could include genuine “hubs” but are probably enriched in false positives (Petre *et al.*, 2015; Petre *et al.*, 2016). In total, six of the interactions we identified have been reported and analyzed in earlier studies (Table S1, Figure 1C). Indeed, as our follow-up analyses of the PexRD12/31 effector family demonstrate, the interactome network we generated can serve as a launchpad for studies of effector activities and the perturbations they cause in host plant cells. We therefore hope that this interactome dataset will complement previous screens of *P. infestans* RXLR effectors aimed at studying effector localization in the host and PTI suppressing activities (Zheng *et al.*, 2014; Wang *et al.*, 2018) and serve as a useful community resource for functional effector biology studies.

Classification of the host interactors into putative functional categories revealed over 35 biological processes that are candidate effector-targeted processes (Table S2). The diversity of these processes is consistent with the view that *Phytophthora* RXLR effectors have evolved to modulate host pathways as diverse as immune signaling, gene silencing, and selective autophagy (Bos *et al.*, 2010; Qiao *et al.*, 2015; Dagdas *et al.*, 2016). However, the interactome network also illustrates a degree of convergent targeting of particular processes both within and between RXLR effector families. Indeed, even though effectors can converge on a few “hub” host proteins (Song *et al.*, 2009; Mukhtar *et al.*, 2011), one challenge is to understand how multiple effectors act on different steps of a host-targeted pathway (Win *et al.*, 2012b). A classic example of this concept can be observed in plant viruses, which encode several proteins that have evolved different mechanisms to suppress host RNA silencing rather than converging on a single host molecule (Burgyán and Havelda, 2011). In another example, *P. infestans* counteract apoplastic cysteine proteases by inhibiting host protease secretion via the RXLR effector AVRblb2 and by secreting protease inhibitors in the apoplast (Tian *et al.*, 2007; Bozkurt *et al.*, 2011). Indeed, the architecture of the interactome we generated indicates a certain degree of convergence between unrelated *P. infestans* effectors towards certain host processes.

In light of the differences between the protein-protein interaction methods described above, it is relevant to note that we recovered relatively few transcription factors or nuclear proteins compared to yeast two-hybrid screens. This could be explained by the protein extraction protocol we used, which may have yielded a proteome relatively depleted in nuclear proteins (Howden and Huitema, 2012). On the other hand, our interactome was enriched in vesicle trafficking proteins possibly reflecting the value of having the effector baits expressed *in planta* in a cellular context that more closely mirrors dynamic cellular processes, such as endomembrane trafficking.

During *P. infestans* infection, RXLR effectors are thought to be delivered inside haustoriated cells where they orchestrate cellular and molecular reprogramming of these plant cells, notably by modulating membrane trafficking (Bozkurt *et al.*, 2012; Dagdas *et al.*, 2016; Dagdas *et al.*, 2018). However, the mechanisms by which *P. infestans* RXLR effectors and other plant pathogen effectors block or hijack host membrane trafficking remain poorly understood. Several RXLR effectors, such as *P. infestans* AVRblb2 and AVR1, *Phytophthora palmivora* REX3, and *Phytophthora brassicae* RxLR24, prevent secretion of host proteins presumably to counteract focal immune responses in the host plant (Bozkurt *et al*., 2011; Du *et al*., 2015; Evangelisti *et al.*, 2017; Tomczynska *et al.*, 2018). Another *P. infestans* effector, PexRD54, stimulates and co-opts plant membrane trafficking by associating with the plant autophagy machinery at polarized foci in the haustorial interface (Dagdas *et al.*, 2018). PexRD54 was also recently proposed to connect small GTPase Rab8a vesicles with the autophagy machinery to stimulate formation of autophagosomes at the pathogen interface (Pandey *et al.*, 2020). In our host-interactor screen, we found that ten effector families associated with 32 proteins implicated in vesicle mediated transport (Figure 2, Figure S3, Table S2). These results, along with our follow-up observation that members of the PexRD12/31 family associate with R-SNARE VAMP72 proteins and accumulate at the haustorial interface, further point to vesicular trafficking as a major target of oomycete effectors.

PexRD12/31 effectors add to a growing list of perihaustorial effectors that are associated with polarized markers at the *P. infestans* haustorial interface (Bozkurt and Kamoun, 2020). We also discovered that the PexRD12/31 family of RXLR-WY effectors associate with 16 vesicle-trafficking *N. benthamiana* proteins in a relatively specific manner (Table S3). However, at this stage we did not determine which of these host proteins, including NbVAMP72x, directly bind the effectors. It remains possible that the PexRD12/31 effectors have the same direct host interactor and that the additional proteins discovered by coIP/MS are part of a multiprotein complex targeted by these effectors. Nevertheless, PexRD12/31 effectors accumulate at the host plasma membrane and, therefore, could potentially co-localize with vesicle trafficking components at this subcellular location. In addition, the examined effectors focally accumulate at the EHM suggesting that they are associated with the dynamic membrane trafficking processes that take place in the infected haustoriated cells. PexRD31 stood out compared to the other examined effectors by labelling mobile vesicle-like puncta; and these PexRD31 vesicle-like structures were visualized in contact with the EHM (Figures 7-9). PexRD31 co-localized with the post-Golgi endosomal marker RabC1, but, unfortunately, the biology of RabC1 is unknown (Geldner *et al.*, 2009). Recently, VAPYRIN, a protein required for the intracellular establishment of arbuscular mycorrhizal fungi, was shown to co-localize with RabC1 in small mobile structures (Bapaume *et al.*, 2019). Interestingly, VAPYRIN also interacts with a symbiotic R-SNARE of the VAMP72 family (Bapaume *et al.*, 2019). Therefore, VAPYRIN/RabC1 mobile bodies might mark an endosomal pathway associated with perimicrobial membranes of plant cells colonized by filamentous microbes. Future studies will determine whether the PexRD31 and VAPYRIN bodies form another example of common cellular structures formed by pathogens and symbionts during intracellular colonization of plant cells (Rey and Schornack, 2013).

We found that PexRD31 and *P. infestans* alter the number and the distribution of FYVE-labelled endosomes in leaves of *N. benthamiana*. How would a pathogen benefit by modulating FYVE-positive endosomes? The 2xFYVE-GFP marker labels vesicles enriched in the phosphoinositide lipid PI3P, notably late endosomes and multivesicular bodies of the late endocytic pathway (Robinson *et al.*, 2008; Gao *et al.*, 2014). Our finding that *P. infestans* alters PI3P vesicles is consistent with previous reports that this pathogen perturbs plant endocytic trafficking, notably by redirecting this pathway to the EHM (Bozkurt *et al.*, 2015). Pathogen-induced modifications in host membrane phosphoinositide composition have recently been proposed both as a potential pathogen strategy to promote infection and as a host focal immune response (Rausche *et al.*, 2020). *P. infestans* may deploy effectors such as PexRD31 to boost the late endocytic pathway, help recruit endomembranes for EHM biogenesis and counter host immunity. Another possibility is that PexRD31 blocks PI3P vesicle fusion to host membranes resulting in the accumulation of these vesicles in the cytosol. Further mechanistic studies of the dynamics of the late endocytic pathway during *P. infestans* infection should reveal the biological significance of the observed phenomenon.

*P. infestans* is an aggressive plant pathogen that continues to threaten global food security (Kamoun *et al.*, 2015). Our study adds to the *in planta* effectoromics screens that have been conducted with *P. infestans* RXLR effectors ever since the sequencing of this pathogen genome over 10 years ago (Haas *et al.*, 2009; Oh *et al.*, 2009; Pais *et al.*, 2013; Zheng *et al.*, 2014; Wang *et al.*, 2018). To date, high-throughput screens have focused primarily on assigning AVR activities to effectors, which helped guide the identification and characterization of matching plant immune receptors (Vleeshouwers *et al.*, 2008; Oh *et al.*, 2009; Rietman, 2011). The interactome network resource we have generated complements these earlier screens and should prove to be a valuable platform for functional studies of *P. infestans* effectors and the processes they target.

## MATERIALS AND METHODS

### Biological materials

We used *Escherichia coli* strain DH5α, *Agrobacterium tumefaciens* strain GV3101 (pMP90), and *Nicotiana benthamiana* as previously described (Petre *et al.*, 2015). Transgenic 2xFYVE-GFP *N. benthamiana* were obtained by transforming the plants with binary plasmid pBLTI221-2xFYVE-GFP (Voigt *et al.*, 2005). *P. infestans* isolates 88069 and 88069td, expressing the red fluorescent marker tandem dimer RFP (tdTomato), were grown and used for *N. benthamiana* leaf infection as previously reported (Dagdas *et al.*, 2016).

### Plasmid construction

Molecular cloning and recombinant DNA manipulations were conducted using standard protocols. Primers and coding sequences of the fusion proteins used in this study are indicated in Table S4 and Table S5.

To generate FLAG-tagged protein fusions, we obtained DNA fragments matching the coding sequence of the effector domains by gene synthesis (GenScript, Pistacaway, USA) including *PacI* and *NotI* restriction sites on 5’ and 3’ ends, respectively (Table S5). Through this process, the coding sequences were codon-optimized for expression in *N. benthamiana* and the N-termini (signal peptide and RXLR regions) were replaced by FLAG tag sequences. We then cloned the DNA fragments into the Tobacco mosaic virus–based *A. tumefaciens* binary vector pTRBO (Lindbo, 2007).

To generate fluorescent protein fusions, we obtained the coding sequence of effector domains by PCR amplification from genomic DNA of *P. infestans* isolate T30-4, using primers that included *BbsI* AATG/GCTT-compatible sites. DNA fragments were cloned into the Golden Gate level 0 vector pICSL41308 by digestion/ligation as previously described (Petre *et al.*, 2015; Petre *et al.*, 2017) and verified by sequencing (GATC Biotech, Constance, Germany). DNA fragments were combined into a Golden Gate level 1 binary vector pICSL47742 in order to create a ‘CaMV 35S promoter::fluorescent protein coding sequence::effector domain coding sequence::OCS terminator’ expression unit.

The following fluorescent markers were described previously: mitochondria (COX4_1_-_29_-GFP), peroxysomes (GFP-PTS1), Golgi apparatus (MAN11-49-GFP), and ER (WAK2_SP_-GFP-HDEL) (Nelson *et al.*, 2007); phosphatidylinositol-3-phospate (PI3P)-positive vesicles (2xFYVE-GFP) (Voigt, 2008); autophagosomes (GFP-ATG8C) (Dagdas *et al.*, 2016). We synthesized the coding sequence of *Arabidopsis thaliana* Exo70E2 (At5g61010) (GenScript) and assembled it with fluorescent proteins into plasmid vector pICH86988 as previously described (Petre *et al.*, 2015).

GFP-NbVAMP72x was cloned using Gateway technology (Thermo Fisher Scientific, Waltham, USA) as follows. Coding sequence for NbVAMP72x was amplified by PCR using the primer pair NbVAMP72x_F and NbVAMP72x_R (Table S4), and Pfu taq polymerase (Takara, Mountain View, USA) from *N. benthamiana* cDNA made from total RNA extracted from 4 week-old *N. benthamiana* leaves. PCR was performed for 35 cycles with denaturing at 96 °C for 30 s, annealing at 58 °C for 40 s and extension at 72 °C for 1 min for each cycle, followed by final extension at 72 °C for 10 min. The amplicon was cloned into pENTR (Thermo Fisher) using Topo TA cloning kit (Thermo Fisher). The insert was transferred to the destination vector pK7WGF2 by LR reaction using Gateway LR Clonase II Enzyme mix (Thermo Fisher) in frame with the coding sequence of GFP which was part of the vector. All Gateway cloning was performed following the manufacturer’s instructions.

### Immunoblotting

We performed immunoblot analyses on SDS-PAGE separated proteins as described elsewhere (Oh *et al.*, 2009). We used Monoclonal FLAG M2-alkaline phosphatase antibodies (Sigma-Aldrich, St-Louis, USA) at a 1:10,000 dilution and we developed blots using the AP conjugate substrate kit (Bio-Rad; Hercules, USA). We used polyclonal GFP antibodies (Thermo Fisher) at a 1:4,000 dilution as primary antibody, and anti-rabbit polyclonal antibody conjugated to horseradish peroxidase (Sigma-Aldrich, St-Louis, USA) as a secondary antibody at a 1:12,000 dilution. We detected protein band signals using ECL substrate (Thermo Fisher) following exposure on Amersham Hyperfilm ECL (GE Healthcare, Chicago, USA).

### Coimmunoprecipitation assays

We performed anti-FLAG coIP/MS assays following a protocol previously described (Win *et al.*, 2011). Briefly, the strategy consisted of affinity purifying transiently expressed FLAG epitope tagged effector proteins and identifying the plant proteins associated with purified effectors by mass spectrometry (MS) (Figure S1). FLAG-tagged effector proteins were immunoprecipitated using agarose beads conjugated with anti-FLAG monoclonal antibodies. Bound effectors and associated plant proteins were competitively eluted by 3XFLAG peptides (Sigma-Aldrich, Dorsett, England) and separated by SDS-PAGE. Each lane from the gel was cut into strips, proteins were digested in gel with trypsin, and submitted for protein identification by MS. For spectral searches, we used a protein sequence database composed of two proteome predictions of *N. benthamiana* genome (Bombarely *et al.*, 2012): (i) proteome version 0.44 available at the Sol Genomics Network (SGN, http://solgenomics.net/) and (ii) evidence-based proteome prediction made by The Genome Analysis Centre (TGAC) based on the same genome sequence. Identical sequences were removed from the combined database to create a non-redundant *N. benthamiana* proteome sequence database. Each sequence in the database was annotated with a top BLASTP hit (e-value cutoff 1 x 10^-3^) to SwissProt protein database if the search was successful. This annotated database was used for Mascot searches using the spectral collection from the mass spectrometry. Mascot results were analyzed using Scaffold (Proteome Software Inc., Oregon, USA). Presence of a protein in the analyzed samples were identified by having at least two peptide matches with equal or more than 95% probability and an ion score of equal or more than 40 in their matches. All protein hits fulfilling these criteria were exported from Scaffold program for further analysis. To reduce complexity of the dataset, we grouped the RXLR effectors into families as previously described (Boutemy *et al.*, 2011) based on their effector domains using a Markov clustering algorithm (MCL) (Enright *et al*., 2002). To further reduce the complexity of the dataset, host protein interactors were also clustered if they shared (i) a minimum of 80% identity on 80% of their lengths, or (ii) the exact same annotation. Each cluster was represented by a single identifier for downstream analyses. Each plant protein identified in coIP/MS was annotated with Gene Ontology (GO) terms using Blast2Go program (Götz *et al.*, 2008).

For a subset of samples, we performed anti-GFP coIP/MS assays and subsequent analyses as previously reported (Win *et al.*, 2011; Petre *et al.*, 2015; Petre *et al.*, 2017), using GFP-Trap agarose beads (Chromotek, Munich, Germany), a hybrid mass spectrometer LTQ-Orbitrap XL (Thermo Fisher Scientific, Carlsbad, California, USA) and a nanoflow-UHPLC system (NanoAcquity Waters Corp., Burnsville, Minnesota, USA). However, to be more stringent during the immunoprecipitation process, we used extraction and immunoprecipitation buffers with 0.5 % IGEPAL and 400 mM NaCl. We performed two technical replicates for the trypsin digestion. For the first replicate, we followed the in-gel digestion protocol as previously reported (Petre *et al.*, 2015). For the second replicate, we performed on-beads digestion as described previously (Zess *et al.*, 2019). Both methods yielded similar results (Figure S9, Dataset 3). We merged the total spectrum count values from the two technical replicates for the analyses shown in Table S3.

### Sequence analyses

We obtained amino acid sequences of the PexRD12/31 family effectors from GenBank based on ‘RxlRfam9’ RXLR family reported in Haas *et al.* (2009). We removed identical sequences and pseudogenes from the family and collected 16 sequences in a database for further analysis (Figure 3). We used effector domain sequences (after ‘EER’ residues, Figure 3) for multiple sequence alignment by MAFFT program using “--auto –reorder” options and L-INS-I strategy (Katoh *et al.*, 2005). Multiple sequence alignment was performed with iterative refinement method (<16) with local pairwise alignment information using amino acid substitution matrix BLOSUM62, 1.53, with the amino acids colored according to the ClustalX scheme (Larkin *et al.*, 2007). The phylogenetic relationship within the PexRD12/31 family members was inferred using the Maximum Likelihood method and JTT matrix-based model (Jones *et al.*, 1992) implemented in MEGA X (Kumar *et al.*, 2018) with 1000 bootstraps. Sequence conservation profile was obtained using the WebLogo server (http://weblogo.berkeley.edu) (Crooks *et al*., 2004). Secondary structure prediction was done using Jpred server (http://www.compbio.dundee.ac.uk/jpred) (Drozdetskiy *et al.*, 2015). WY-fold region was predicted based on the hidden Markov model (HMM) reported in Boutemy *et al.* (2011). HMMER v3.1b (Mistry *et al.*, 2013) was used to search the WY-fold HMM in amino acid sequences of PexRD12/31 family of effectors.

The sequence of NbS00022342g0004 was searched against the *A. thaliana* proteome version Araport11 (The Arabidopsis Information Resource (TAIR), https://www.arabidopsis.org) using BLASTP (BLAST 2.9.0+) (Altschul *et al.*, 1997) as provided by TAIR server. To attempt to identify an ortholog of NbS00022342g0004 in *A. thaliana*, we extracted all AtVAMP72 family (AtVAMP721-727) from Araport11 and 14 VAMP72-like amino acid sequences from the proteome of *N. benthamiana* (Sol Genomics Network, version 0.44). Multiple Sequence alignment was performed with MAFFT program using “--auto –reorder” options and L-INS-I strategy (Katoh *et al.*, 2005). Phylogeny was inferred by using the Maximum Likelihood method and JTT matrix-based model implemented in MEGA X (Kumar *et al.*, 2018). Initial tree(s) for the heuristic search were obtained automatically by applying Neighbor-Join and BioNJ algorithms to a matrix of pairwise distances estimated using the JTT model, and then selecting the topology with superior log likelihood value after 1000 bootstraps.

### Laser-scanning confocal microscopy

We collected leaves of *N. benthamiana* three days post agroinfiltration and immediately performed live-cell imaging with a Leica DM6000B/TCS SP5 laser-scanning confocal microscope (Leica microsystems, Bucks, UK), using a 63x water immersion objective as previously described (Petre *et al*., 2015). We used the following settings for excitation/collection of fluorescence: GFP (488/505-525 nm), chlorophyll (488/680-700 nm), and mCherry (561/580-620 nm). We performed image analysis with Fiji (http://fiji.sc/Fiji).

### Quantification of FYVE endosomes in transient assays

To quantify the 2xFYVE-GFP puncta in confocal images acquired from *N. benthamiana* leaf cells transiently expressing different mCherry or RFP-fused constructs, we built a pipeline using the ImageJ package Fiji (Schindelin *et al.*, 2012). We split the channels (GFP, RFP, and bright field) and considered only the GFP channel. We then smoothed the images with the Smooth tool. Third, we performed an auto threshold using the RenyiEntropy parameter. Finally, we counted the puncta per image with the Analyze particles tool, with the following settings: size = 0.07 – 5; circularity = 0.3 – 1. In parallel, we calculated an estimate of the area of cytoplasm using the total GFP signal in the image. To do this, we first applied the gaussian blur tool with a Sigma = 5. We then applied the Threshold function using the Huang white method. After that, we obtained the area of cytoplasm using the Measure tool. The number of GFP puncta calculated in every image was divided by the total area of cytoplasm calculated, obtaining the number of GFP labelled puncta/μm^2^ of cytoplasm. We then exported the quantification results in a spreadsheet and generated the scatterplot using R.

For the quantification of 2xFYVE-GFP puncta in confocal images acquired from *N. benthamiana* leaf cells transiently expressing FLAG-tagged *Phytophthora* effectors and free RFP, a different automated pipeline was employed, as described in (Salomon *et al.*, 2010).

### Quantitative analysis of confocal images during *P. infestans* leaf colonization

To quantify the 2xFYVE-GFP puncta in confocal images acquired on *N. benthamiana* leaf cells colonized by *P. infestans* isolate 88069td, we built a four-step analytical pipeline using the ImageJ package Fiji (Schindelin *et al.*, 2012). First, we split the channels (GFP, RFP, chlorophyll, and bright field) and considered only the GFP channel. Second, we smoothed the images with the Smooth tool. Third, we performed an auto threshold using the RenyiEntropy parameter. Finally, we counted the puncta per image with the Analyze particles tool, with the following settings: size = 0.5 – infinity; circularity = 0.5 – 1. To quantify the presence of *P. infestans* isolate 88069td on *N. benthamiana* leaves, we adapted the first and last steps of the pipeline abovementioned as follow: for the first step, we considered the RFP channel, and for the fourth step, we used the default settings for particle size and circularity, i.e. 0 – infinity and 0 – 1, respectively. We then exported the quantification results in a spreadsheet and generated the scatterplot using R (R Core Team, 2019).

### Data availability and accession numbers

GenBank accession numbers for the effectors used in this study are presented in Table 1. The Solanaceae Genomics Network (https://solgenomics.net) accession numbers for *N. benthamiana* proteins are listed in Dataset 1.

The mass spectrometry proteomics data were deposited to the ProteomeXchange Consortium via the PRIDE (Perez-Riverol *et al.*, 2019) partner repository (https://www.ebi.ac.uk/pride/archive) with the dataset identifier PXD020751 and 10.6019/PXD020751.

## ACKNOWLEDGEMENTS

We are thankful to several colleagues for discussions and ideas. Over the course of this research project, the Kamoun Lab was funded primarily from the Gatsby Charitable Foundation, Biotechnology and Biological Sciences Research Council (BBSRC, UK), and European Research Council (ERC; NGRB and BLASTOFF projects).

## References

Alfano, J.R. (2009). Roadmap for future research on plant pathogen effectors. Molecular plant pathology 10, 805–813.

Altschul, S.F., Madden, T.L., Schäffer, A.A., Zhang, J., Zhang, Z., Miller, W., and Lipman, D.J. (1997). Gapped BLAST and PSI-BLAST: a new generation of protein database search programs. Nucleic Acids Res. 25, 3389–3402.

Anderson, R.G., Deb, D., Fedkenheuer, K., and McDowell, J.M. (2015). Recent progress in RXLR effector research. Molecular Plant-Microbe Interactions 28, 1063–1072.

Asai, S., and Shirasu, K. (2015). Plant cells under siege: plant immune system versus pathogen effectors. Current opinion in plant biology 28, 1–8.

Bapaume, L., Laukamm, S., Darbon, G., Monney, C., Meyenhofer, F., Feddermann, N., Chen, M., and Reinhardt, D. (2019). VAPYRIN marks an endosomal trafficking compartment involved in arbuscular mycorrhizal symbiosis. Frontiers in plant science 10, 666.

Boevink, P.C., Wang, X., McLellan, H., He, Q., Naqvi, S., Armstrong, M.R., Zhang, W., Hein, I., Gilroy, E.M., and Tian, Z. (2016). A Phytophthora infestans RXLR effector targets plant PP1c isoforms that promote late blight disease. Nature communications 7, 1–14.

Bombarely, A., Rosli, H.G., Vrebalov, J., Moffett, P., Mueller, L.A., and Martin, G.B. (2012). A draft genome sequence of Nicotiana benthamiana to enhance molecular plant-microbe biology research. Molecular Plant-Microbe Interactions 25, 1523–1530.

Bos, J.I., Armstrong, M.R., Gilroy, E.M., Boevink, P.C., Hein, I., Taylor, R.M., Zhendong, T., Engelhardt, S., Vetukuri, R.R., and Harrower, B. (2010). Phytophthora infestans effector AVR3a is essential for virulence and manipulates plant immunity by stabilizing host E3 ligase CMPG1. Proceedings of the National Academy of Sciences 107, 9909–9914.

Boutemy, L.S., King, S.R., Win, J., Hughes, R.K., Clarke, T.A., Blumenschein, T.M., Kamoun, S., and Banfield, M.J. (2011). Structures of Phytophthora RXLR effector proteins a conserved but adaptable fold underpins functional diversity. Journal of Biological Chemistry 286, 35834–35842.

Bozkurt, T.O., and Kamoun, S. (2020). The plant–pathogen haustorial interface at a glance. Journal of Cell Science 133.

Bozkurt, T.O., Schornack, S., Banfield, M.J., and Kamoun, S. (2012). Oomycetes, effectors, and all that jazz. Current opinion in plant biology 15, 483–492.

Bozkurt, T.O., Belhaj, K., Dagdas, Y.F., Chaparro-Garcia, A., Wu, C.H., Cano, L.M., and Kamoun, S. (2015). Rerouting of plant late endocytic trafficking toward a pathogen interface. Traffic 16, 204–226.

Bozkurt, T.O., Schornack, S., Win, J., Shindo, T., Ilyas, M., Oliva, R., Cano, L.M., Jones, A.M., Huitema, E., van der Hoorn, R.A., and Kamoun, S. (2011). Phytophthora infestans effector AVRblb2 prevents secretion of a plant immune protease at the haustorial interface. Proceedings of the National Academy of Sciences 108, 20832–20837.

Burgyán, J., and Havelda, Z. (2011). Viral suppressors of RNA silencing. Trends Plant Sci. 16, 265–272.

Chaparro-Garcia, A., Schwizer, S., Sklenar, J., Yoshida, K., Petre, B., Bos, J.I., Schornack, S., Jones, A.M., Bozkurt, T.O., and Kamoun, S. (2015). Phytophthora infestans RXLR-WY effector AVR3a associates with dynamin-related protein 2 required for endocytosis of the plant pattern recognition receptor FLS2. PLoS One 10, e0137071.

Collins, N.C., Thordal-Christensen, H., Lipka, V., Bau, S., Kombrink, E., Qiu, J.-L., Hückelhoven, R., Stein, M., Freialdenhoven, A., and Somerville, S.C. (2003). SNARE-protein-mediated disease resistance at the plant cell wall. Nature 425, 973–977.

Cooke, D.E.L., Cano, L.M., Raffaele, S., Bain, R.A., Cooke, L.R., Etherington, G.J., Deahl, K.L., Farrer, R.A., Gilroy, E.M., Goss, E.M., Grünwald, N.J., Hein, I., MacLean, D., McNicol, J.W., Randall, E., Oliva, R.F., Pel, M.A., Shaw, D.S., Squires, J.N., Taylor, M.C., Vleeshouwers, V.G.A.A., Birch, P.R.J., Lees, A.K., and Kamoun, S. (2012). Genome Analyses of an Aggressive and Invasive Lineage of the Irish Potato Famine Pathogen. PLOS Pathogens 8, e1002940.

Crooks, G.E., Hon, G., Chandonia, J.-M., and Brenner, S.E. (2004). WebLogo: a sequence logo generator. Genome Res. 14, 1188–1190.

Dagdas, Y.F., Pandey, P., Tumtas, Y., Sanguankiattichai, N., Belhaj, K., Duggan, C., Leary, A.Y., Segretin, M.E., Contreras, M.P., Savage, Z., Khandare, V.S., Kamoun, S., and Bozkurt, T.O. (2018). Host autophagy machinery is diverted to the pathogen interface to mediate focal defense responses against the Irish potato famine pathogen. Elife 7.

Dagdas, Y.F., Belhaj, K., Maqbool, A., Chaparro-Garcia, A., Pandey, P., Petre, B., Tabassum, N., Cruz-Mireles, N., Hughes, R.K., Sklenar, J., Win, J., Menke, F., Findlay, K., Banfield, M.J., Kamoun, S., and Bozkurt, T.O. (2016). An effector of the Irish potato famine pathogen antagonizes a host autophagy cargo receptor. Elife 5, e10856.

Derevnina, L., Kamoun, S., and Wu, C.h. (2019). Dude, where is my mutant? Nicotiana benthamiana meets forward genetics. New Phytologist 221, 607–610.

Derevnina, L., Petre, B., Kellner, R., Dagdas, Y.F., Sarowar, M.N., Giannakopoulou, A., De la Concepcion, J.C., Chaparro-Garcia, A., Pennington, H.G., Van West, P., and Kamoun, S. (2016). Emerging oomycete threats to plants and animals. Philosophical Transactions of the Royal Society B: Biological Sciences 371, 20150459.

Dodds, P.N., and Rathjen, J.P. (2010). Plant immunity: towards an integrated view of plant-pathogen interactions. Nat Rev Genet 11, 539–548.

Drozdetskiy, A., Cole, C., Procter, J., and Barton, G.J. (2015). JPred4: a protein secondary structure prediction server. Nucleic Acids Res. 43, W389–W394.

Du, Y., Mpina, M.H., Birch, P.R., Bouwmeester, K., and Govers, F. (2015). Phytophthora infestans RXLR effector AVR1 interacts with exocyst component Sec5 to manipulate plant immunity. Plant Physiology 169, 1975–1990.

Enright, A.J., Van Dongen, S., and Ouzounis, C.A. (2002). An efficient algorithm for large-scale detection of protein families. Nucleic acids research 30, 1575–1584.

Evangelisti, E., Gogleva, A., Hainaux, T., Doumane, M., Tulin, F., Quan, C., Yunusov, T., Floch, K., and Schornack, S. (2017). Time-resolved dual transcriptomics reveal early induced Nicotiana benthamiana root genes and conserved infection-promoting Phytophthora palmivora effectors. BMC Biol. 15, 39.

Fisher, M.C., Henk, D.A., Briggs, C.J., Brownstein, J.S., Madoff, L.C., McCraw, S.L., and Gurr, S.J. (2012). Emerging fungal threats to animal, plant and ecosystem health. Nature 484, 186–194.

Gao, C., Luo, M., Zhao, Q., Yang, R., Cui, Y., Zeng, Y., Xia, J., and Jiang, L. (2014). A unique plant ESCRT component, FREE1, regulates multivesicular body protein sorting and plant growth. Curr. Biol. 24, 2556–2563.

Gawehns, F., Cornelissen, B.J., and Takken, F.L. (2013). The potential of effector-target genes in breeding for plant innate immunity. Microbial biotechnology 6, 223–229.

Geldner, N., Dénervaud-Tendon, V., Hyman, D.L., Mayer, U., Stierhof, Y.D., and Chory, J. (2009). Rapid, combinatorial analysis of membrane compartments in intact plants with a multicolor marker set. The Plant Journal 59, 169–178.

Giannakopoulou, A., Schornack, S., Bozkurt, T.O., Haart, D., Ro, D.-K., Faraldos, J.A., Kamoun, S., and O’Maille, P.E. (2014). Variation in capsidiol sensitivity between Phytophthora infestans and Phytophthora capsici is consistent with their host range. PLoS One 9.

Goodin, M.M., Zaitlin, D., Naidu, R.A., and Lommel, S.A. (2008). Nicotiana benthamiana: its history and future as a model for plant–pathogen interactions. Mol. Plant-Microbe Interact. 21, 1015–1026.

Götz, S., García-Gómez, J.M., Terol, J., Williams, T.D., Nagaraj, S.H., Nueda, M.J., Robles, M., Talón, M., Dopazo, J., and Conesa, A. (2008). High-throughput functional annotation and data mining with the Blast2GO suite. Nucleic Acids Res. 36, 3420–3435.

Haas, B.J., Kamoun, S., Zody, M.C., Jiang, R.H., Handsaker, R.E., Cano, L.M., Grabherr, M., Kodira, C.D., Raffaele, S., Torto-Alalibo, T., Bozkurt, T.O., Ah-Fong, A.M., Alvarado, L., Anderson, V.L., Armstrong, M.R., Avrova, A., Baxter, L., Beynon, J., Boevink, P.C., Bollmann, S.R., Bos, J.I., Bulone, V., Cai, G., Cakir, C., Carrington, J.C., Chawner, M., Conti, L., Costanzo, S., Ewan, R., Fahlgren, N., Fischbach, M.A., Fugelstad, J., Gilroy, E.M., Gnerre, S., Green, P.J., Grenville-Briggs, L.J., Griffith, J., Grunwald, N.J., Horn, K., Horner, N.R., Hu, C.H., Huitema, E., Jeong, D.H., Jones, A.M., Jones, J.D., Jones, R.W., Karlsson, E.K., Kunjeti, S.G., Lamour, K., Liu, Z., Ma, L., Maclean, D., Chibucos, M.C., McDonald, H., McWalters, J., Meijer, H.J., Morgan, W., Morris, P.F., Munro, C.A., O’Neill, K., Ospina-Giraldo, M., Pinzon, A., Pritchard, L., Ramsahoye, B., Ren, Q., Restrepo, S., Roy, S., Sadanandom, A., Savidor, A., Schornack, S., Schwartz, D.C., Schumann, U.D., Schwessinger, B., Seyer, L., Sharpe, T., Silvar, C., Song, J., Studholme, D.J., Sykes, S., Thines, M., van de Vondervoort, P.J., Phuntumart, V., Wawra, S., Weide, R., Win, J., Young, C., Zhou, S., Fry, W., Meyers, B.C., van West, P., Ristaino, J., Govers, F., Birch, P.R., Whisson, S.C., Judelson, H.S., and Nusbaum, C. (2009). Genome sequence and analysis of the Irish potato famine pathogen Phytophthora infestans. Nature 461, 393–398.

He, J., Ye, W., Choi, D.S., Wu, B., Zhai, Y., Guo, B., Duan, S., Wang, Y., Gan, J., and Ma, W. (2019). Structural analysis of Phytophthora suppressor of RNA silencing 2 (PSR2) reveals a conserved modular fold contributing to virulence. Proceedings of the National Academy of Sciences 116, 8054–8059.

Heard, W., Sklenář, J., Tome, D.F., Robatzek, S., and Jones, A.M. (2015). Identification of regulatory and cargo proteins of endosomal and secretory pathways in Arabidopsis thaliana by proteomic dissection. Molecular & Cellular Proteomics 14, 1796–1813.

Howden, A.J.M., and Huitema, E. (2012). Effector-triggered post-translational modifications and their role in suppression of plant immunity. Frontiers in plant science 3, 160.

Ivanov, S., Austin, J., Berg, R.H., and Harrison, M.J. (2019). Extensive membrane systems at the host–arbuscular mycorrhizal fungus interface. Nature Plants 5, 194–203.

Jiang, R.H., Tripathy, S., Govers, F., and Tyler, B.M. (2008). RXLR effector reservoir in two Phytophthora species is dominated by a single rapidly evolving superfamily with more than 700 members. Proceedings of the National Academy of Sciences 105, 4874–4879.

Jones, D.T., Taylor, W.R., and Thornton, J.M. (1992). The rapid generation of mutation data matrices from protein sequences. Bioinformatics 8, 275–282.

Kamoun, S., Furzer, O., Jones, J.D.G., Judelson, H.S., Ali, G.S., Dalio, R.J.D., Roy, S.G., Schena, L., Zambounis, A., Panabières, F., Cahill, D., Ruocco, M., Figueiredo, A., Chen, X.-R., Hulvey, J., Stam, R., Lamour, K., Gijzen, M., Tyler, B.M., Grünwald, N.J., Mukhtar, M.S., Tomé, D.F.A., Tör, M., Van Den Ackerveken, G., McDowell, J., Daayf, F., Fry, W.E., Lindqvist-Kreuze, H., Meijer, H.J.G., Petre, B., Ristaino, J., Yoshida, K., Birch, P.R.J., and Govers, F. (2015). The Top 10 oomycete pathogens in molecular plant pathology. Mol. Plant Pathol. 16, 413–434.

Katoh, K., Kuma, K.-i., Toh, H., and Miyata, T. (2005). MAFFT version 5: improvement in accuracy of multiple sequence alignment. Nucleic Acids Res. 33, 511–518.

Kumar, S., Stecher, G., Li, M., Knyaz, C., and Tamura, K. (2018). MEGA X: molecular evolutionary genetics analysis across computing platforms. Mol. Biol. Evol. 35, 1547–1549.

Kwon, C., Neu, C., Pajonk, S., Yun, H.S., Lipka, U., Humphry, M., Bau, S., Straus, M., Kwaaitaal, M., Rampelt, H., Kasmi, F.E., Jürgens, G., Parker, J., Panstruga, R., Lipka, V., and Schulze-Lefert, P. (2008). Co-option of a default secretory pathway for plant immune responses. Nature 451, 835–840.

Larkin, M.A., Blackshields, G., Brown, N.P., Chenna, R., McGettigan, P.A., McWilliam, H., Valentin, F., Wallace, I.M., Wilm, A., and Lopez, R. (2007). Clustal W and Clustal X version 2.0. Bioinformatics 23, 2947–2948.

Lee, A.H.Y., Petre, B., and Joly, D.L. (2013). Effector wisdom. New Phytologist 197, 375–377.

Lee, S.-J., and Rose, J.K.C. (2010). Mediation of the transition from biotrophy to necrotrophy in hemibiotrophic plant pathogens by secreted effector proteins. Plant Signaling & Behavior 5, 769–772.

Lindbo, J.A. (2007). TRBO: a high-efficiency tobacco mosaic virus RNA-based overexpression vector. Plant physiology 145, 1232–1240.

Lu, Y.J., Schornack, S., Spallek, T., Geldner, N., Chory, J., Schellmann, S., Schumacher, K., Kamoun, S., and Robatzek, S. (2012). Patterns of plant subcellular responses to successful oomycete infections reveal differences in host cell reprogramming and endocytic trafficking. Cellular microbiology 14, 682–697.

Maqbool, A., Hughes, R.K., Dagdas, Y.F., Tregidgo, N., Zess, E., Belhaj, K., Round, A., Bozkurt, T.O., Kamoun, S., and Banfield, M.J. (2016). Structural Basis of Host Autophagy-related Protein 8 (ATG8) Binding by the Irish Potato Famine Pathogen Effector Protein PexRD54. J Biol Chem 291, 20270–20282.

Mellacheruvu, D., Wright, Z., Couzens, A.L., Lambert, J.-P., St-Denis, N.A., Li, T., Miteva, Y.V., Hauri, S., Sardiu, M.E., Low, T.Y., Halim, V.A., Bagshaw, R.D., Hubner, N.C., al-Hakim, A., Bouchard, A., Faubert, D., Fermin, D., Dunham, W.H., Goudreault, M., Lin, Z.-Y., Badillo, B.G., Pawson, T., Durocher, D., Coulombe, B., Aebersold, R., Superti-Furga, G., Colinge, J., Heck, A.J.R., Choi, H., Gstaiger, M., Mohammed, S., Cristea, I.M., Bennett, K.L., Washburn, M.P., Raught, B., Ewing, R.M., Gingras, A.-C., and Nesvizhskii, A.I. (2013). The CRAPome: a contaminant repository for affinity purification–mass spectrometry data. Nature Methods 10, 730–736.

Mistry, J., Finn, R.D., Eddy, S.R., Bateman, A., and Punta, M. (2013). Challenges in homology search: HMMER3 and convergent evolution of coiled-coil regions. Nucleic Acids Res. 41, e121–e121.

Mukhtar, M.S., Carvunis, A.-R., Dreze, M., Epple, P., Steinbrenner, J., Moore, J., Tasan, M., Galli, M., Hao, T., and Nishimura, M.T. (2011). Independently evolved virulence effectors converge onto hubs in a plant immune system network. science 333, 596–601.

Nelson, B., Cai, X., and Nebenführ, A. (2007). A multi-color set of in vivo organelle markers for colocalization studies in organelle markers for co-localisation studies in Arabidopsis and other plants: Fluorescent organelle markers. (Plant J.), pp. 1126–1136.

Oh, S.-K., Young, C., Lee, M., Oliva, R., Bozkurt, T.O., Cano, L.M., Win, J., Bos, J.I., Liu, H.-Y., van Damme, M., Morgan, W., Choi, D., Van der Vossen, E.A.G., Vleeshouwers, V.G.A.A., and Kamoun, S. (2009). In planta expression screens of Phytophthora infestans RXLR effectors reveal diverse phenotypes, including activation of the Solanum bulbocastanum disease resistance protein Rpi-blb2. The Plant Cell 21, 2928–2947.

Oliveira-Garcia, E., and Valent, B. (2015). How eukaryotic filamentous pathogens evade plant recognition. Current opinion in microbiology 26, 92–101.

Pais, M., Win, J., Yoshida, K., Etherington, G.J., Cano, L.M., Raffaele, S., Banfield, M.J., Jones, A., Kamoun, S., and Saunders, D.G. (2013). From pathogen genomes to host plant processes: the power of plant parasitic oomycetes. Genome biology 14, 211.

Pais, M., Yoshida, K., Giannakopoulou, A., Pel, M.A., Cano, L.M., Oliva, R.F., Witek, K., Lindqvist-Kreuze, H., Vleeshouwers, V.G., and Kamoun, S. (2018). Gene expression polymorphism underpins evasion of host immunity in an asexual lineage of the Irish potato famine pathogen. BMC Evol. Biol. 18, 93.

Pandey, P., Leary, A.Y., Tümtas, Y., Savage, Z., Dagvadorj, B., Tan, E., Khandare, V., Duggan, C., Yusunov, T., Madalinski, M., Mirkin, F.G., Schornack, S., Dagdas, Y., Kamoun, S., and Bozkurt, T.O. (2020). The Irish potato famine pathogen subverts host vesicle trafficking to channel starvation-induced autophagy to the pathogen interface. bioRxiv, 2020.03.20.000117.

Perez-Riverol, Y., Csordas, A., Bai, J., Bernal-Llinares, M., Hewapathirana, S., Kundu, D.J., Inuganti, A., Griss, J., Mayer, G., and Eisenacher, M. (2019). The PRIDE database and related tools and resources in 2019: improving support for quantification data. Nucleic Acids Res. 47, D442–D450.

Petre, B., Win, J., Menke, F.L.H., and Kamoun, S. (2017). Protein-Protein Interaction Assays with Effector-GFP Fusions in Nicotiana benthamiana. Methods Mol Biol 1659, 85–98.

Petre, B., Saunders, D.G., Sklenar, J., Lorrain, C., Win, J., Duplessis, S., and Kamoun, S. (2015). Candidate effector proteins of the rust pathogen Melampsora larici-populina target diverse plant cell compartments. Molecular Plant-Microbe Interactions 28, 689–700.

Petre, B., Saunders, D.G., Sklenar, J., Lorrain, C., Krasileva, K.V., Win, J., Duplessis, S., and Kamoun, S. (2016). Heterologous Expression Screens in Nicotiana benthamiana Identify a Candidate Effector of the Wheat Yellow Rust Pathogen that Associates with Processing Bodies. PLoS One 11, e0149035.

Qiao, Y., Shi, J., Zhai, Y., Hou, Y., and Ma, W. (2015). Phytophthora effector targets a novel component of small RNA pathway in plants to promote infection. Proceedings of the National Academy of Sciences 112, 5850–5855.

Rausche, J., Stenzel, I., Stauder, R., Fratini, M., Trujillo, M., Heilmann, I., and Rosahl, S. (2020). A phosphoinositide 5-phosphatase from Solanum tuberosum is activated by PAMP-treatment and may antagonize phosphatidylinositol 4, 5-bisphosphate at Phytophthora infestans infection sites. New Phytol.

R Core Team (2020). R: A language environment for statistical computing. R Foundation for Statistical Computing, Vienna, Austria. URL http://www.R-project.org/.

Ren, Y., Armstrong, M., Qi, Y., McLellan, H., Zhong, C., Du, B., Birch, P.R., and Tian, Z. (2019). Phytophthora infestans RXLR effectors target parallel steps in an immune signal transduction pathway. Plant Physiol. 180, 2227–2239.

Rey, T., and Schornack, S. (2013). Interactions of beneficial and detrimental root-colonizing filamentous microbes with plant hosts. Genome biology 14, 121.

Rietman, H. (2011). Putting the Phytophthora infestans genome sequence at work: multiple novel avirulence and potato resistance gene candidates revealed.

Robinson, D.G., Jiang, L., and Schumacher, K. (2008). The endosomal system of plants: charting new and familiar territories. Plant Physiol. 147, 1482–1492.

Rovenich, H., Boshoven, J.C., and Thomma, B.P. (2014). Filamentous pathogen effector functions: of pathogens, hosts and microbiomes. Current opinion in plant biology 20, 96–103.

Salomon, S., Grunewald, D., Stüber, K., Schaaf, S., MacLean, D., Schulze-Lefert, P., and Robatzek, S. (2010). High-throughput confocal imaging of intact live tissue enables quantification of membrane trafficking in Arabidopsis. Plant Physiol. 154, 1096–1104.

Saunders, D.G., Breen, S., Win, J., Schornack, S., Hein, I., Bozkurt, T.O., Champouret, N., Vleeshouwers, V.G., Birch, P.R., Gilroy, E.M., and Kamoun, S. (2012). Host protein BSL1 associates with Phytophthora infestans RXLR effector AVR2 and the Solanum demissum immune receptor R2 to mediate disease resistance. The Plant Cell 24, 3420–3434.

Schindelin, J., Arganda-Carreras, I., Frise, E., Kaynig, V., Longair, M., Pietzsch, T., Preibisch, S., Rueden, C., Saalfeld, S., and Schmid, B. (2012). Fiji: an open-source platform for biological-image analysis. Nature methods 9, 676–682.

Sharpee, W.C., and Dean, R.A. (2016). Form and function of fungal and oomycete effectors. Fungal Biology Reviews 30, 62–73.

Song, J., Win, J., Tian, M., Schornack, S., Kaschani, F., Ilyas, M., van der Hoorn, R.A.L., and Kamoun, S. (2009). Apoplastic effectors secreted by two unrelated eukaryotic plant pathogens target the tomato defense protease Rcr3. Proceedings of the National Academy of Sciences 106, 1654–1659.

Tian, M., Win, J., Song, J., van der Hoorn, R., van der Knaap, E., and Kamoun, S. (2007). A Phytophthora infestans cystatin-like protein targets a novel tomato papain-like apoplastic protease. Plant Physiol. 143, 364–377.

Tomczynska, I., Stumpe, M., and Mauch, F. (2018). A conserved RxLR effector interacts with host RABA-type GTPases to inhibit vesicle-mediated secretion of antimicrobial proteins. The Plant Journal 95, 187–203.

Toruño, T.Y., Stergiopoulos, I., and Coaker, G. (2016). Plant-pathogen effectors: cellular probes interfering with plant defenses in spatial and temporal manners. Annual review of phytopathology 54, 419–441.

Toufexi, A., Duggan, C., Pandey, P., Savage, Z., Segretin, M.E., Yuen, L.H., Gaboriau, D.C., Leary, A.Y., Khandare, V., Ward, A.D., Botchway, S.W., Bateman, B.C., Pan, I., Schattat, M., Sparkes, I., and Bozkurt, T.O. (2019). Chloroplasts navigate towards the pathogen interface to counteract infection by the Irish potato famine pathogen. bioRxiv, 516443.

Turnbull, D., Yang, L., Naqvi, S., Breen, S., Welsh, L., Stephens, J., Morris, J., Boevink, P.C., Hedley, P.E., Zhan, J., Birch, P.R., and Gilroy, E.M. (2017). RXLR effector AVR2 up-regulates a brassinosteroid-responsive bHLH transcription factor to suppress immunity. Plant Physiology 174, 356–369.

van Schie, C.C., and Takken, F.L. (2014). Susceptibility genes 101: how to be a good host. Annual review of phytopathology 52, 551–581.

van West, P., de Jong, A.J., Judelson, H.S., Emons, A.M.C., and Govers, F. (1998). The ipiO Gene of Phytophthora infestans Is Highly Expressed in Invading Hyphae during Infection. Fungal Genet. Biol. 23, 126–138.

Vleeshouwers, V.G., Raffaele, S., Vossen, J.H., Champouret, N., Oliva, R., Segretin, M.E., Rietman, H., Cano, L.M., Lokossou, A., Kessel, G., Pel, M.A., and Kamoun, S. (2011). Understanding and exploiting late blight resistance in the age of effectors. Annual review of phytopathology 49, 507–531.

Vleeshouwers, V.G.A.A., Rietman, H., Krenek, P., Champouret, N., Young, C., Oh, S.-K., Wang, M., Bouwmeester, K., Vosman, B., Visser, R.G.F., Jacobsen, E., Govers, F., Kamoun, S., and Van der Vossen, E.A.G. (2008). Effector Genomics Accelerates Discovery and Functional Profiling of Potato Disease Resistance and Phytophthora Infestans Avirulence Genes. PLOS ONE 3, e2875.

Voigt, B. (2008). Actin cytoskeleton and endomembrane system dependent cell growth (Universitäts-und Landesbibliothek Bonn).

Voigt, B., Timmers, A.C., Šamaj, J., Hlavacka, A., Ueda, T., Preuss, M., Nielsen, E., Mathur, J., Emans, N., Stenmark, H., Nakano, A., Baluška, F., and Menzel, D. (2005). Actin-based motility of endosomes is linked to the polar tip growth of root hairs. Eur. J. Cell Biol. 84, 609–621.

Vossen, J.H., Nijenhuis, M., Arens-De Reuver, M.J.B., Van Der Vossen, E.A.G., Jacobsen, E., and Visser, R.G.F. (2011). Cloning and exploitation of a functional R-gene from Solanum chacoense. US Patent Application No. WO2011034433A1.

Wang, S., McLellan, H., Bukharova, T., He, Q., Murphy, F., Shi, J., Sun, S., van Weymers, P., Ren, Y., Thilliez, G., Wang, H., Chen, X., Engelhardt, S., Vleeshouwers, V., Gilroy, E.M., Whisson, S.C., Hein, I., Wang, X., Tian, Z., Birch, P.R.J., and Boevink, P.C. (2018). Phytophthora infestans RXLR effectors act in concert at diverse subcellular locations to enhance host colonization. Journal of Experimental Botany 70, 343–356.

Wang, X., Boevink, P., McLellan, H., Armstrong, M., Bukharova, T., Qin, Z., and Birch, P.R. (2015). A host KH RNA-binding protein is a susceptibility factor targeted by an RXLR effector to promote late blight disease. Molecular plant 8, 1385–1395.

Wei, P., Wong, W.W., Park, J.S., Corcoran, E.E., Peisajovich, S.G., Onuffer, J.J., Weiss, A., and Lim, W.A. (2012). Bacterial virulence proteins as tools to rewire kinase pathways in yeast and immune cells. Nature 488, 384–388.

Weßling, R., Epple, P., Altmann, S., He, Y., Yang, L., Henz, S.R., McDonald, N., Wiley, K., Bader, K.C., and Gläßer, C. (2014). Convergent targeting of a common host protein-network by pathogen effectors from three kingdoms of life. Cell host & microbe 16, 364–375.

Whisson, S.C., Boevink, P.C., Moleleki, L., Avrova, A.O., Morales, J.G., Gilroy, E.M., Armstrong, M.R., Grouffaud, S., Van West, P., and Chapman, S. (2007). A translocation signal for delivery of oomycete effector proteins into host plant cells. Nature 450, 115–118.

Win, J., Kamoun, S., and Jones, A.M. (2011). Purification of effector–target protein complexes via transient expression in Nicotiana benthamiana. In Plant Immunity (Springer), pp. 181–194.

Win, J., Krasileva, K.V., Kamoun, S., Shirasu, K., Staskawicz, B.J., and Banfield, M.J. (2012a). Sequence divergent RXLR effectors share a structural fold conserved across plant pathogenic oomycete species. PLoS Path. 8, e1002400.

Win, J., Chaparro-Garcia, A., Belhaj, K., Saunders, D., Yoshida, K., Dong, S., Schornack, S., Zipfel, C., Robatzek, S., Hogenhout, S., and Kamoun, S. (2012b). Effector Biology of Plant-Associated Organisms: Concepts and Perspectives. Cold Spring Harb. Symp. Quant. Biol., pp. 235–247.

Yang, L., McLellan, H., Naqvi, S., He, Q., Boevink, P.C., Armstrong, M., Giuliani, L.M., Zhang, W., Tian, Z., and Zhan, J. (2016). Potato NPH3/RPT2-like protein StNRL1, targeted by a Phytophthora infestans RXLR effector, is a susceptibility factor. Plant physiology 171, 645–657.

Yoshida, K., Schuenemann, V.J., Cano, L.M., Pais, M., Mishra, B., Sharma, R., Lanz, C., Martin, F.N., Kamoun, S., Krause, J., Thines, M., Weigel, D., and Burbano, H.A. (2013). The rise and fall of the Phytophthora infestans lineage that triggered the Irish potato famine. Elife 2, e00731.

Yun, H.S., and Kwon, C. (2017). Vesicle trafficking in plant immunity. Curr Opin Plant Biol 40, 34–42.

Zess, E.K., Jensen, C., Cruz-Mireles, N., De la Concepcion, J.C., Sklenar, J., Stephani, M., Imre, R., Roitinger, E., Hughes, R., Belhaj, K., Mechtler, K., Menke, F.L.H., Bozkurt, T.O., Banfield, M.J., Kamoun, S., Maqbool, A., and Dagdas, Y.F. (2019). N-terminal β-strand underpins biochemical specialization of an ATG8 isoform. PLoS Biol. 17, e3000373.

Zheng, X., McLellan, H., Fraiture, M., Liu, X., Boevink, P.C., Gilroy, E.M., Chen, Y., Kandel, K., Sessa, G., Birch, P.R., and Brunner, F. (2014). Functionally redundant RXLR effectors from Phytophthora infestans act at different steps to suppress early flg22-triggered immunity. PLoS Pathog 10, e1004057.

