## Supplementary material for "Host-interactor screens of *Phytophthora infestans* RXLR proteins reveal vesicle trafficking as a major effector-targeted process": Petre et al Supplemental Figures and Tables 24Sep2020 .pdf

#### Figure S1

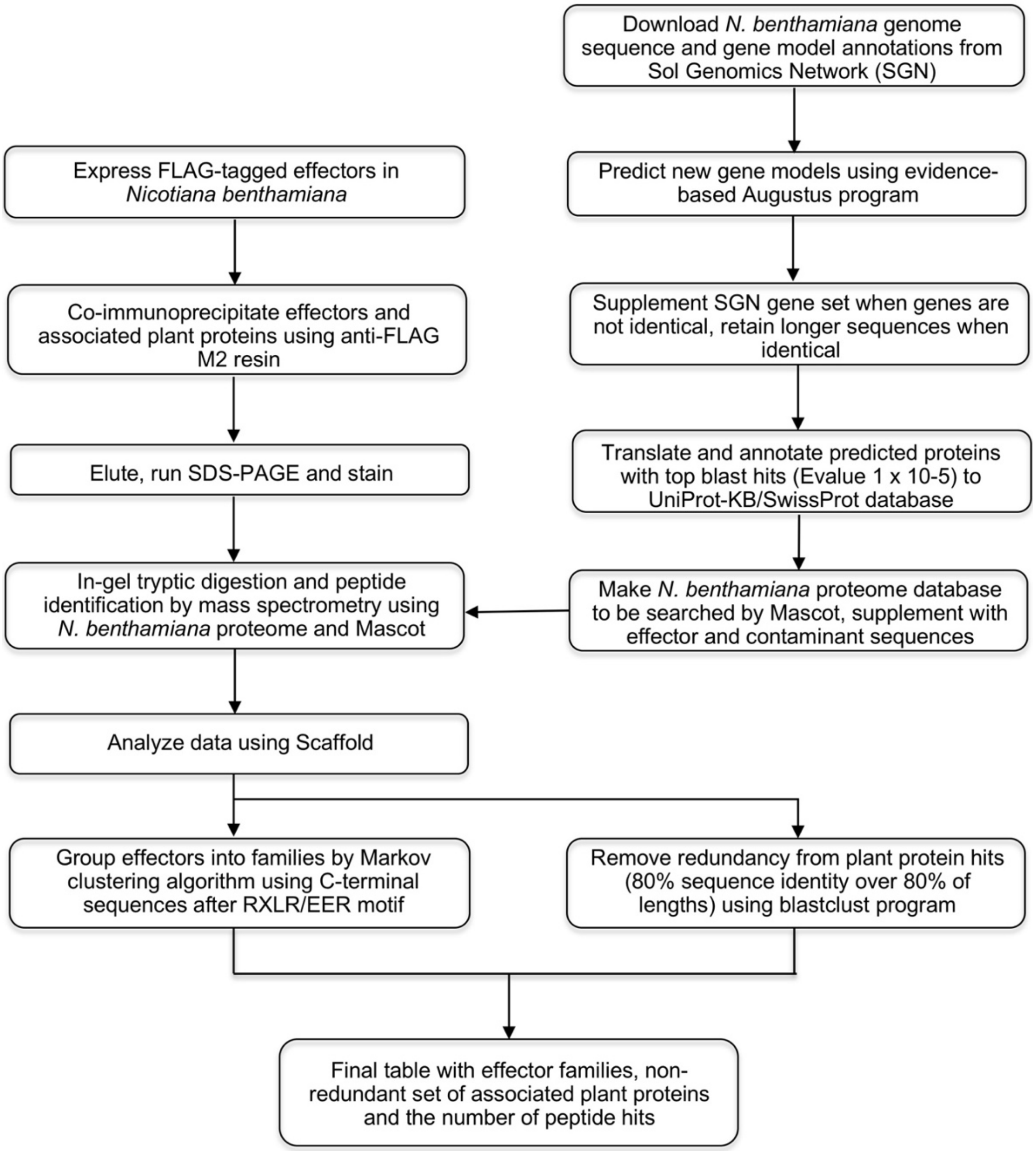


**Figure S1.** Strategy to screen for effector-associated plant proteins in *Nicotiana benthamiana*

#### Figure S2

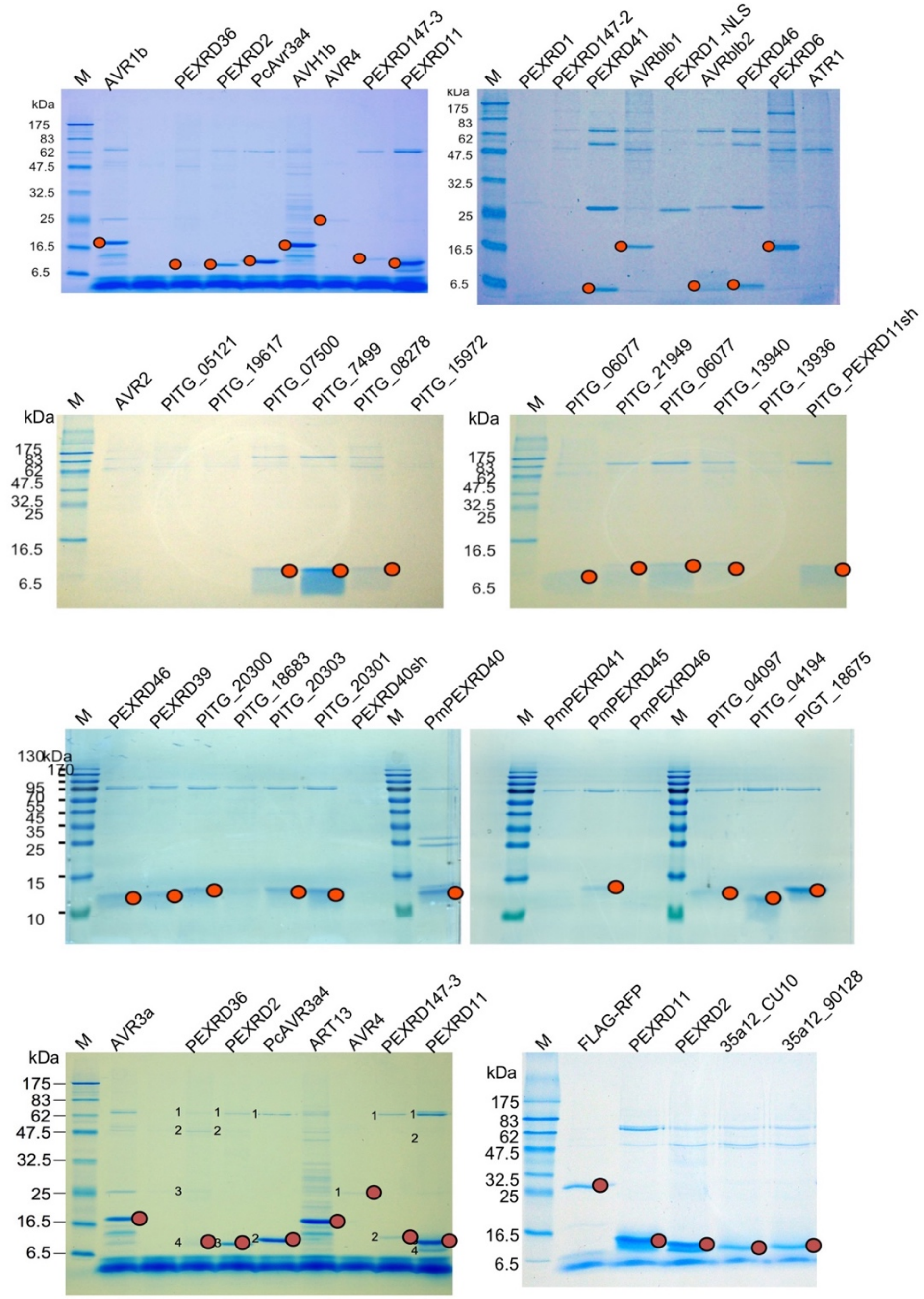


**Figure S2.** Anti-FLAG immunoprecipitation efficiently purifies the effector fusions

Protein mixtures isolated by anti-FLAG immunoprecipitation were reduced and denatured in a Laemmli buffer, then subjected to SDS-PAGE and Coomassie blue staining. Approximate location of the band signals matching the expected size of the FLAG-tagged effector fusion are indicated with red filled circles. M = PageRuler Plus standard molecular weight marker. Numbers indicate the size of marker bands in kilodalton.

### Figure S3

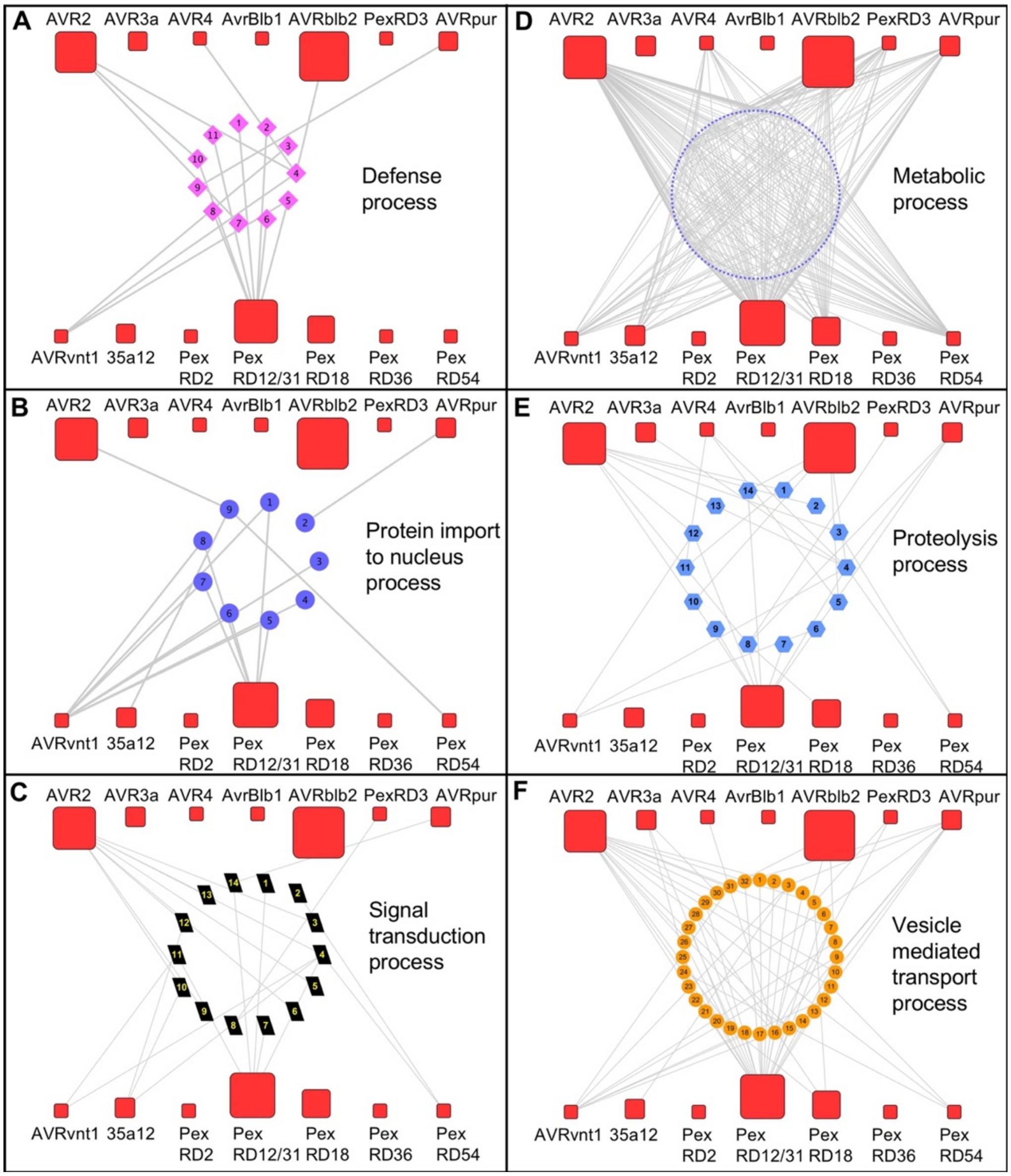


**Figure S3.** Sub-networks of the RXLR interactome organized by GO terms

(**A**) Sub-network with host proteins involved in defense processes.
(**B**) Sub-network with host proteins involved in protein import to nucleus processes.
(**C**) Sub-network with host proteins involved in signal transduction processes.
(**D**) Sub-network with host proteins involved in metabolic processes.
(**E**) Sub-network with host proteins involved in proteolysis processes.
(**F**) Sub-network with host proteins involved in vesicle mediated transport processes.

Effector families are depicted in rounded red squares and the sizes of the squares correspond to the sizes of the effector families.

#### Figure S4

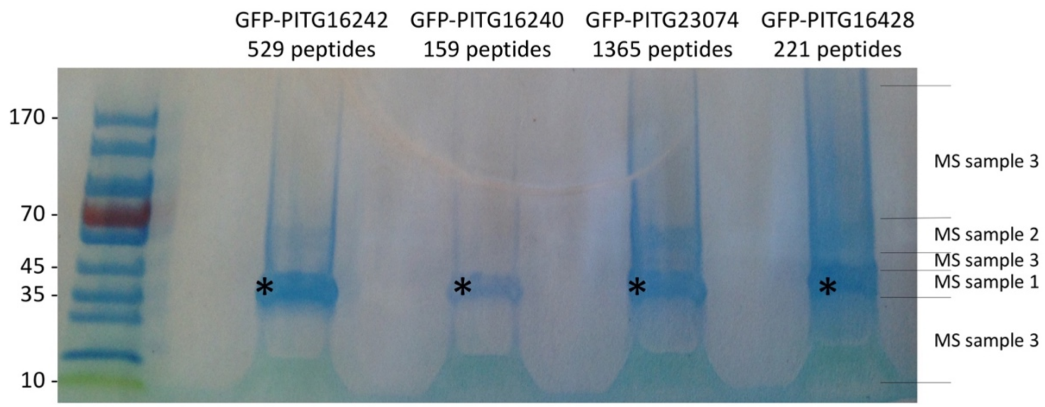


**Figure S4.** Anti-GFP coimmunoprecipitation efficiently purifies GFP-tagged fusions

Protein mixtures isolated by anti-GFP immunoprecipitation were reduced and denatured in a Laemmli buffer, then subjected to SDS-PAGE/CCB staining assays. Trypsin-digested peptides were processed by LC-MS/MS and collected peaks were used to search a database containing the sequence of the GFP fusions. For each GFP fusion, the number of peptides identified by LC-MS/MS and matching their sequence is indicated. The size of selected bands of the page ruler is indicated in kilodalton. Black asterisks indicate the band signal matching the expected size of the GFP fusion. For each lane, the area of gel cut and processed as an independent sample (numbered 1 to 3) for LC-MS/MS is indicated on the right-hand side.

### Figure S5

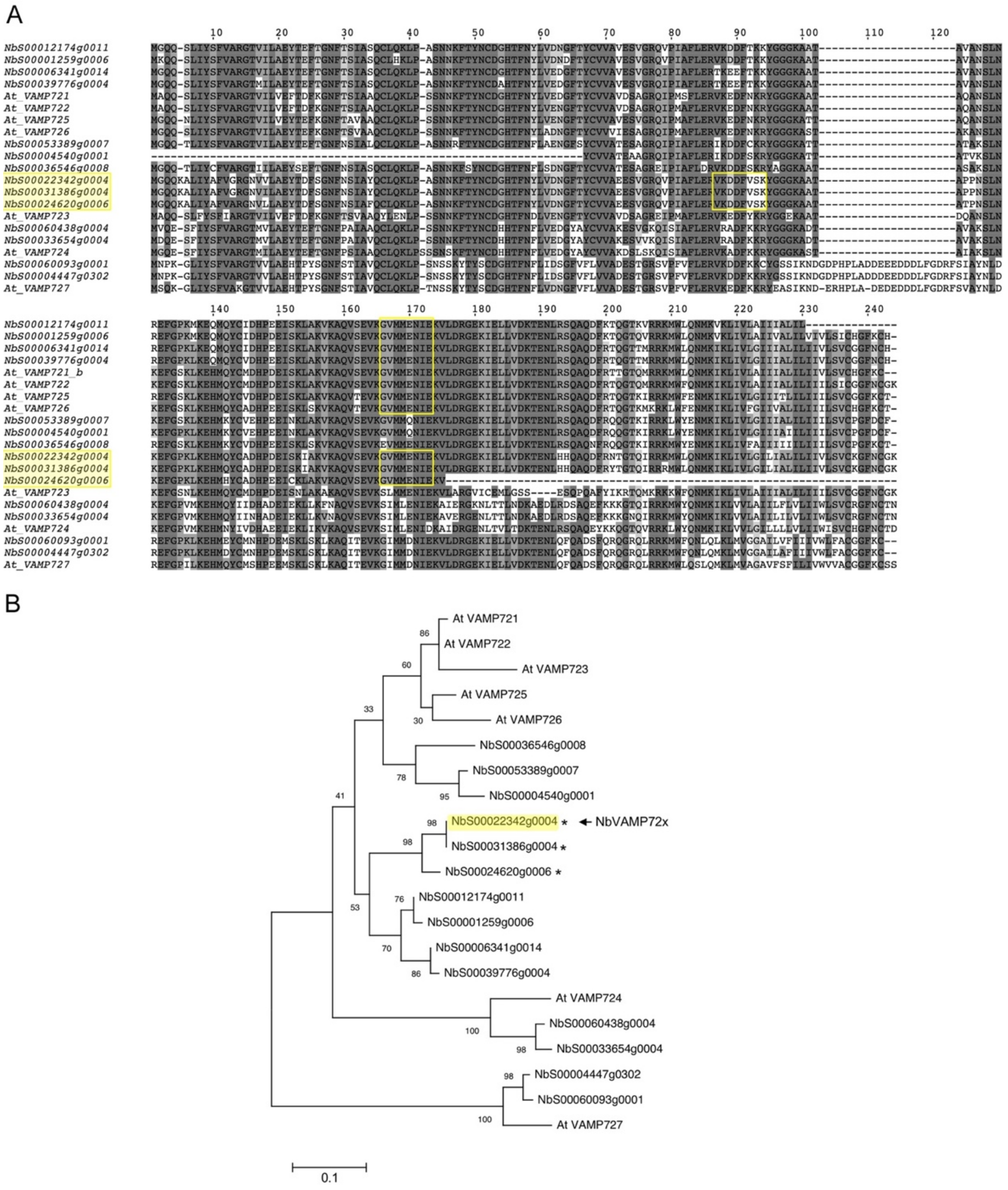


**Figure S5.** NbVAMP associated with PexRD12/31 effectors belongs to VAMP72 family in *N. benthamiana*

**(A)** Multiple sequence alignment of VAMP 72 family from *N. benthamiana* (SGN database accession numbers were shown) and *A. thaliana* (Accession numbers of *A. thaliana* VAMPs: At_VAMP721_b = NP_171967.1; At_VAMP722 = P47192.2; At_VAMP723 = NP_850201.1; At_VAMP724 = O23429.2; At_VAMP725 = O48850.2; At_VAMP726 = Q9MAS5.2; At_VAMP727 = NP_001078283.1); *N. benthamiana* VAMPs identified by two peptide hits in MS searches are highlighted in yellow; peptide hits were shown in yellow rectangle box.

**(B)** Phylogenetic tree of VAMP72 family from *N. benthamiana* and *A. thaliana*. The phylogeny was inferred by using the Maximum Likelihood method and JTT matrix-based model. The tree with the highest log likelihood (-930.02) is shown. The percentage of trees in which the associated taxa clustered together is shown next to the branches. Initial tree(s) for the heuristic search were obtained automatically by applying Neighbor-Join and BioNJ algorithms to a matrix of pairwise distances estimated using the JTT model, and then selecting the topology with superior log likelihood value. The tree is drawn to scale, with branch lengths measured in the number of substitutions per site. This analysis involved 21 amino acid sequences. All positions containing gaps and missing data were eliminated (complete deletion option). There were a total of 88 positions in the final dataset. Evolutionary analyses were conducted in MEGA X. Cloned NbVAMP is highlighted in yellow; * = VAMP identified by two peptide hits.

**Figure S6**

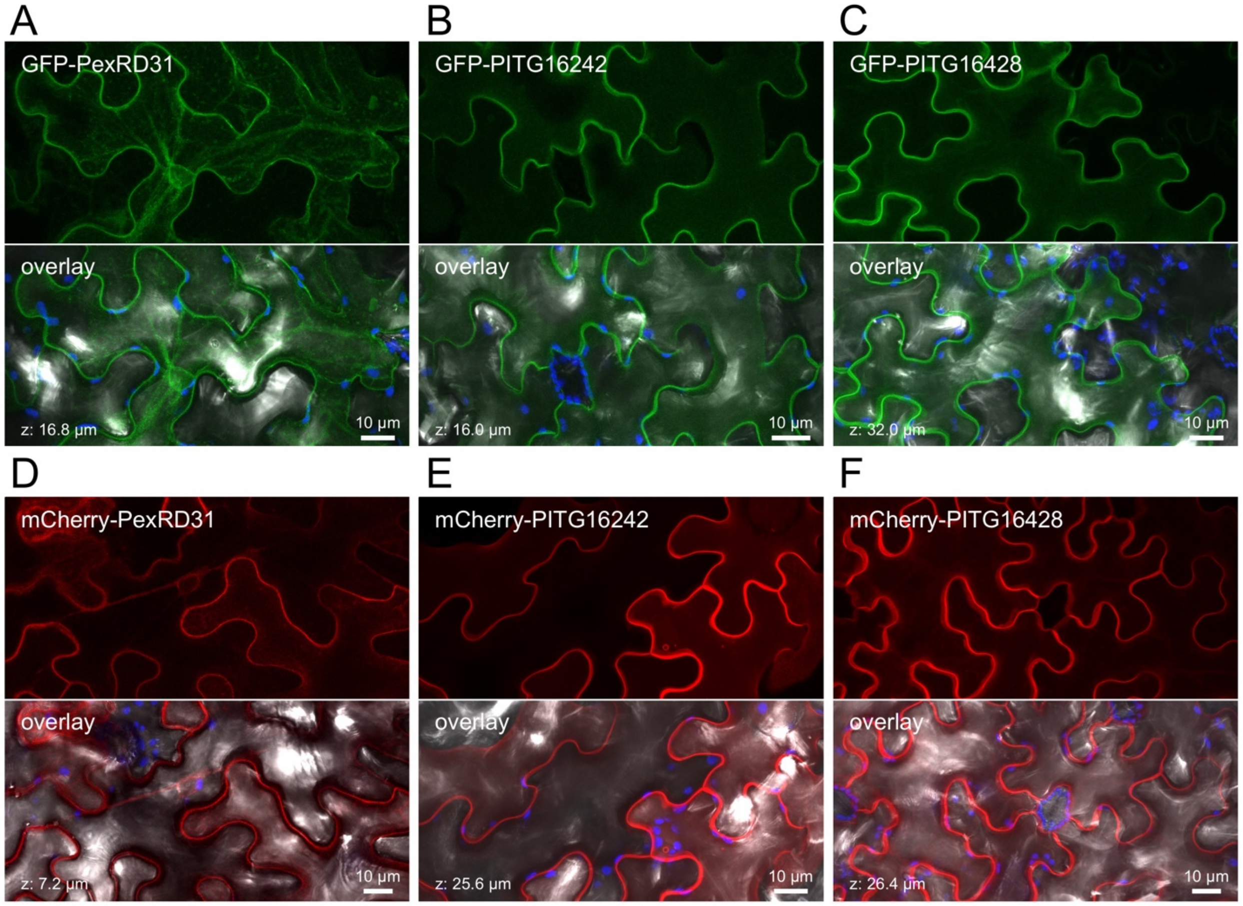


**Figure S6.** PexRD12/31 effectors accumulate mainly at the cell periphery

**(A)** Live-cell imaging of a green fluorescent protein (GFP)-PexRD31 fusion in *Nicotiana benthamiana* leaves.
**(B)** Live-cell imaging of GFP-PITG16242.
**(C)** Live-cell imaging of GFP-PITG16428.
**(D)** Live-cell imaging of mCherry-PexRD31.
**(E)** Live-cell imaging of mCherry-PITG16242.
**(F)** Live-cell imaging of mCherry-PITG16428.

Proteins were expressed in leaf cells by agroinfiltration. Live-cell imaging was performed with a laser-scanning confocal microscope three days after infiltration. GFP and chlorophyll were excited at 488 nm; mCherry was excited at 561 nm. GFP (green), mCherry (red), and chlorophyll (blue) fluorescence were collected between 505 and 525 nm, 580 and 620 nm, and 680 and 700 nm, respectively. Images are maximal projections of up to 40 optical sections (maximal z-stack of 32.0 µm). The overlay panels combine fluorescent protein (GFP or mCherry), chlorophyll, and bright field images.

### Figure S7

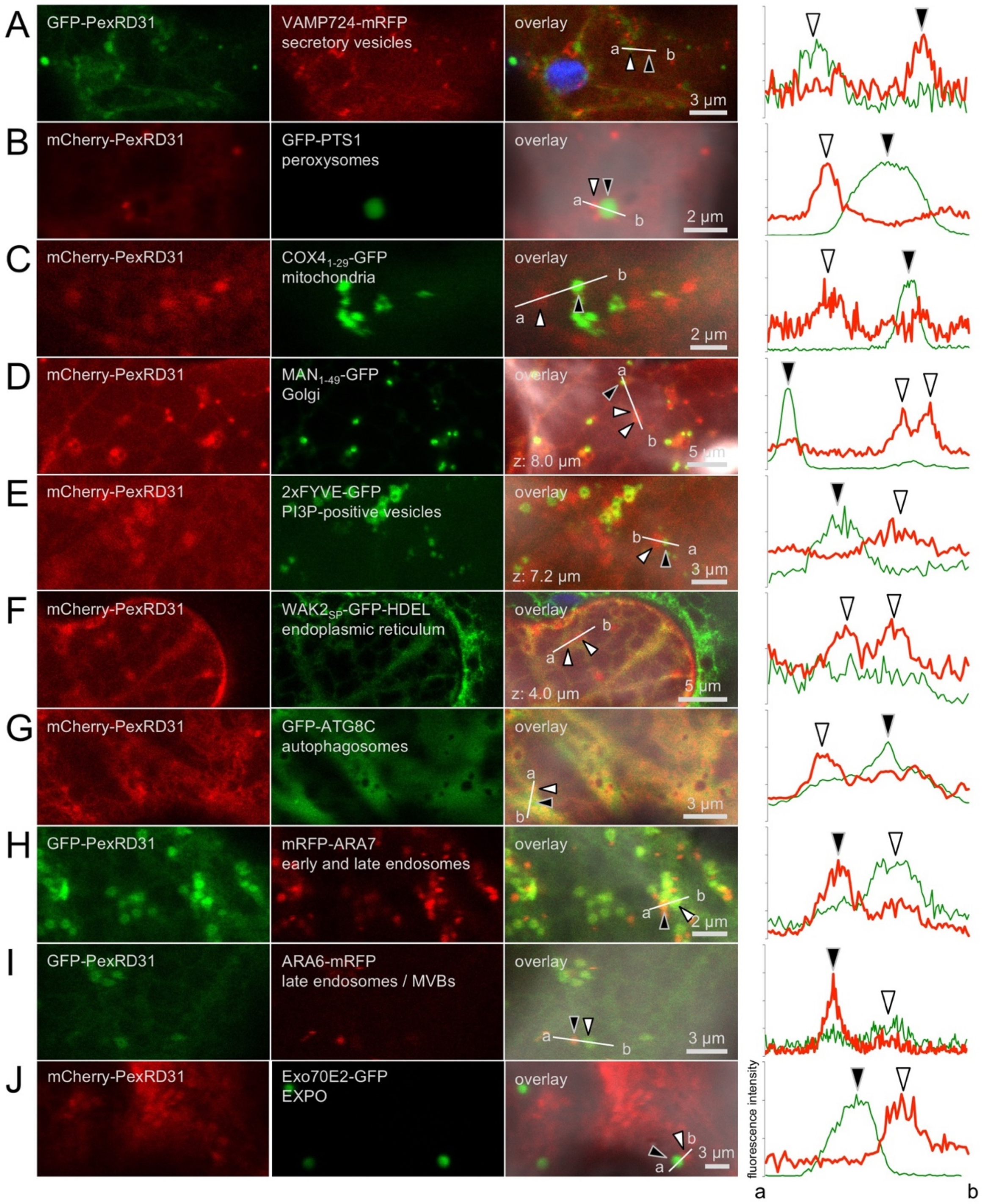


**Figure S7.** PexRD31 does not accumulate in the main cell compartments

Live-cell imaging of
**(A)** green fluorescent protein (GFP)-PexRD31 and NBVAMP72X-monomeric red fluorescent protein (mRFP, secretory vesicles marker) in *N. benthamiana* leaves.
**(B)** mCherry-PexRD31 and GFP-PTS1 (peroxysomes marker) in *N. benthamiana* leaves.
**(C)** mCherry-PexRD31 and COX4$_{1-29}$-GFP (mitochondria marker) in *N. benthamiana* leaves.
**(D)** mCherry-PexRD31 and MAN1$_{1-49}$-GFP (Golgi marker) in *N. benthamiana* leaves.
**(E)** mCherry-PexRD31and 2xFYVE-GFP (Phosphatidylinositol-3-Phosphate [PI3P]-positive vesicles marker) in *N. benthamiana* leaves.

**(F)** mCherry-PexRD31 and $WAK2_{signal\ peptide}$-GFP-HDEL (endoplasmic reticulum [ER] marker) in *N. benthamiana* leaves.
**(G)** mCherry-PexRD31 and GFP-ATG8C (selective autophagosomes marker) in *N. benthamiana* leaves.
**(H)** GFP-PexRD31 and mRFP-ARA7 (early and late endosomes [EE\LE] marker) in *N. benthamiana* leaves.
**(I)** GFP-PexRD31 and ARA6-mRFP (late endosomes and multivesicular bodies [LE\MVB] marker) in *N. benthamiana* leaves.
**(J)** mCherry-PexRD31 and EXO70E2-GFP (Exocyst-positive organelle [EXPO] marker) in *N. benthamiana* leaves.

Proteins were expressed in leaf cells by agroinfiltration. Live-cell imaging was performed with a laser-scanning confocal microscope three days after infiltration. GFP and chlorophyll were excited at 488 nm; mCherry was excited at 561 nm. GFP (green), mCherry (red), and chlorophyll (blue) fluorescence were collected between 505 and 525 nm, 580 and 620 nm, and 680 and 700 nm, respectively. Images are single optical sections of 0.8 µm or maximal projections of up to 10 optical sections (max. z-stack: 8.0 µm). The overlay panel combines GFP, mCherry, chlorophyll, and bright field channels. The right-hand side panel show relative fluorescence intensity plots of the GFP and the mCherry along the line from a to b depicted in the corresponding overlay panel. In the intensity plots, white and black arrowheads indicate fluorescence peaks as visible in the overlay panel.

**Figure S8**

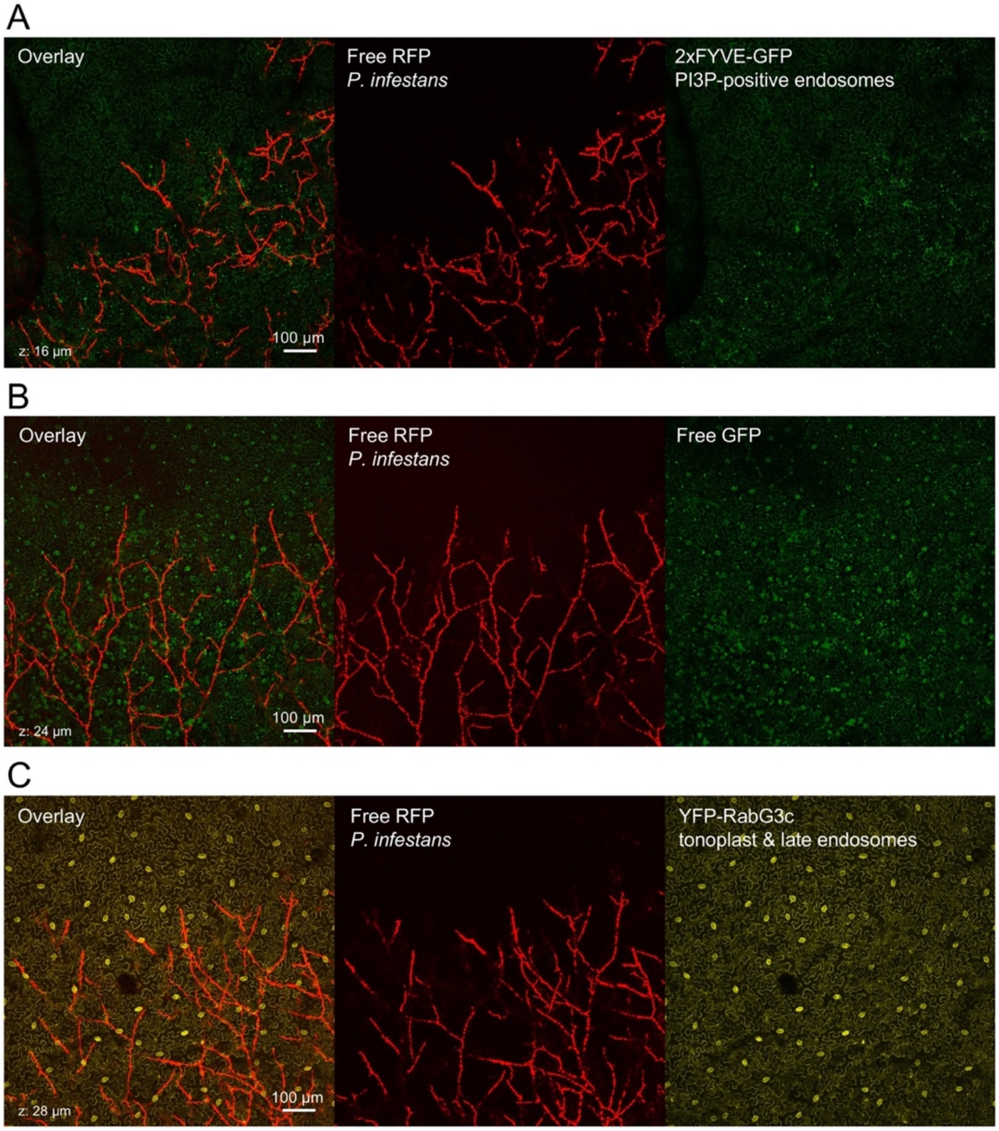


**Figure S8.** *P. infestans* biotrophic colonization does not trigger the formation of punctate signals in the nucleocytosolic, late endosomal, and tonoplast compartments

Live cell imaging of
**(A)** a 2xFYVE-GFP fusion (marker of PI3P-positive endosomes)
**(B)** a free GFP (marker of the nucleus and cytosol), and
**(C)** a YFP-RabG3c fusion (marker of late endosomes and tonoplast) in *N. benthamiana* leaf cells colonized by *P. infestans* isolate 88069td. Leaves of stable transgenic *N. benthamiana* plants were drop inoculated by zoospores of *P. infestans* isolate 88069td. Live-cell imaging was performed with a laser-scanning confocal microscope three days after inoculation. GFP, YFP, and RFP were excited at 488 nm, 514 nm, and 561 nm, respectively. GFP (green), YFP (yellow), and RFP (red) fluorescence were collected between 505 and 525 nm, 525 and 550 nm, and 580 and 620 nm, respectively. Images are maximal projections of up to 35 optical sections (max. z-stack of 28 µm). The overlay panels on the left-hand side combines either **(A-B)** the GFP and RFP channels or **(C)** the YFP and RFP channels. Note the presence of GFP puncta in **(A)** (positive control) but not in **(B-C)**.

#### Figure S9

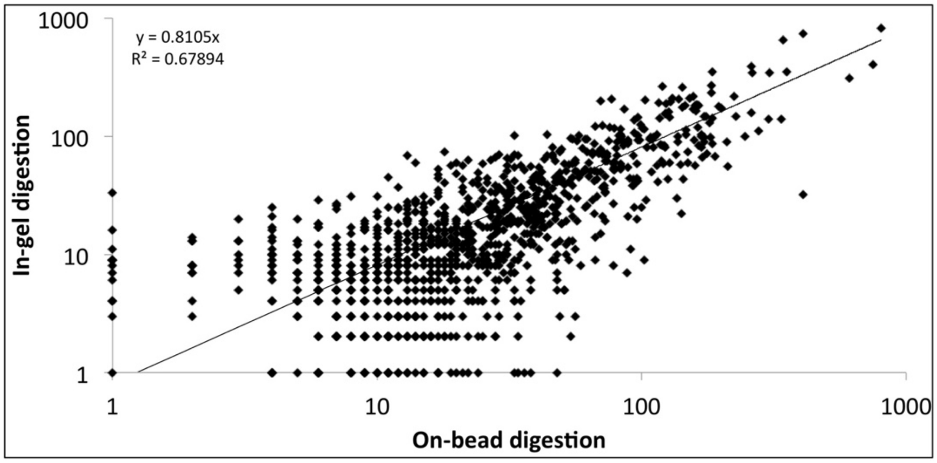


**Figure S9.** In-gel and on-beads trypsin digestion methods yield similar data

Total spectrum count values (all samples merged, added of one unit) from in-gel and on-beads trypsin digestion methods were plotted using Microsoft Excel 'Marked Scatter' graph function. Linear regression and associated parameters were calculated using the same software. Both axes have a Log scale.

### SUPPLEMENTAL TABLES

#### **Table S1.** List of experimentally validated effector-host protein associations

| Effector | Plant protein | Number of associated effector families | References |
|---|---|---|---|
| **AVRblb2** | Cysteine proteinase RD21a (C14) | 3 | Bozkurt et al., 2011 |
| **AVR2** | BSL1 serine/threonine protein phosphatase | 1 | Saunders et al., 2012 |
| **AVR2** | BSL3 serine/threonine-protein phosphatase | 1 | Turnbull et al., 2019 |
| **AVR3a** | Dynamin 2B | 5 | Chaparro-Garcia et al., 2015 |
| **PexRD54** | ATG8 | N/A | Maqbool et al., 2016 |
| **PexRD54** | Ras-related protein RabE1c | 4 | Pandey et al., 2020 |
| **PexRD12/31 (7 effectors)** | Vesicle associated membrane protein 7B (NbVAMP72x) | 2 | This study |

**Table S2.** Biological processes targeted by *Phytophthora infestans* effectors

| Biological process[1] | Effector family[2] | | | | | | | | | | | | | |
|---|---|---|---|---|---|---|---|---|---|---|---|---|---|---|
| | AVR2 | AVR3a | AVR4 | AVRblb1 | AVRblb2 | PexRD3 | AVRpur | AVRvnt1 | PexRD12 | PexRD18 | PexRD2 | PexRD36 | PexRD54 | 35a12 |
| ATP hydrolysis coupled proton transport | 3 | | 1 | | 2 | | 1 | 4 | 5 | 1 | | 2 | | |
| ATP synthesis coupled proton transport | 7 | 1 | 5 | | 5 | | 2 | 1 | 4 | 2 | | | | 1 |
| Actin cytoskeleton organization | | | | | 1 | 1 | 2 | 2 | 2 | 1 | | | | |
| Apoptotic process | | | | | | | | | 1 | | | | | |
| Cell development | 2 | | 1 | | 1 | 1 | 1 | 1 | 1 | | | | | |
| Cell redox homeostasis | 4 | | | | | | | | | | | | | |
| Cell wall biogenesis | 1 | | | | 1 | 1 | 1 | 1 | 5 | 1 | | | | |
| Cell wall modification | 2 | | | | | | | | | | | | | |
| Cellular response to reactive oxygen Species | 1 | | | | | | | | | | | | | |
| Chloroplast organization | 1 | | | | | | | | 1 | | | | | |
| Cytokinesis | | | | | 1 | | | | 3 | | | | | |
| DNA repair | | | | | | 1 | 1 | | 2 | | | | | |
| Defense response | 3 | | 1 | | 1 | | 1 | 3 | 7 | | | | | |
| Detection of biotic stimulus | 1 | | | | | | | | | 1 | | | | |
| Electron transport | 2 | | | | | | | | 1 | | | | | |
| Metabolic process | 67 | | 15 | | 2 | 23 | 23 | 19 | 66 | 2 | | 1 | | 16 |
| Methylation | 1 | | | | | | | | | | | | | |
| Microtubule cytoskeleton organization | 2 | | 1 | | 1 | | 2 | 2 | 3 | 1 | | | | |
| Microtubule-based movement | 2 | | 1 | | 1 | | 2 | 3 | 3 | | | | | 1 |
| Negative regulation of peptidase activity | 2 | | | | 2 | 1 | | 1 | | | | | | |
| Nucleosome assembly | 3 | 2 | 1 | | 3 | | 1 | | 1 | 1 | | | | |
| Oxidation-reduction process | 3 | | | | 1 | | | | 1 | | | | | |
| Phagocytosis | 1 | | | | | | | | | | | | | |
| Photosynthesis | 27 | | 7 | 1 | 12 | 4 | 4 | 7 | 12 | 8 | 1 | 2 | | 8 |
| Proteasome-mediated ubiquitin-dependent protein catabolic process | 13 | | 2 | | 2 | 2 | 14 | 11 | 28 | 2 | | | | 1 |
| Protein folding | 25 | 4 | 6 | 4 | 1 | 3 | 14 | 1 | 2 | 7 | 4 | 2 | | 7 |
| Protein Import into chloroplast stroma | 1 | | | | | 1 | | | 2 | | | | | |
| Protein import into nucleus | 1 | | | | | | 1 | 7 | 5 | | | | | 1 |
| Protein phosphorylation | | | | | | | | | 1 | | | | | |
| Protein stabilization | | | | | | | | | 1 | | | | | |
| Protein targeting to chloroplast | 1 | | | | | | 1 | | 1 | | | | | |
| Protein targeting to mitochondrion | 1 | | | | | | | | | | | | | |
| Protein targeting to vacuole | 1 | | | | | | | | | | | | | |
| Proteolysis | 8 | 1 | 3 | | 5 | 1 | 3 | 3 | 8 | 1 | | | | |
| RNA processing | 5 | 1 | 2 | | 2 | 1 | 2 | 2 | 5 | 3 | | | | |
| RNA transcription | | | | | 2 | | | | | | | | | |
| RNA-dependent DNA replication | 1 | | | | | 1 | 1 | | 1 | | | | | |
| Regulation of cellular process | | | 1 | | 1 | | | | | | | | | |
| Removal of superoxide radicals | 1 | | | | | | | | | | | | | |
| Response to heat | | | | | | | 1 | | | | | | | |
| Response to stress | 1 | | | | | | | | | | | | | |
| Signal transduction | 7 | | | | | 1 | 1 | 2 | 5 | | | | | 2 |
| Thylakoid membrane organization | | | 1 | | 1 | | | | 3 | | | | | |
| Translation | 51 | | 4 | | 49 | 3 | 6 | 67 | 82 | 14 | 2 | 1 | | 11 |
| Transport | 1 | | 4 | | 6 | 5 | 3 | 8 | 26 | 3 | 1 | | | |
| Unassigned | 19 | | 2 | | 3 | 8 | 4 | 4 | 18 | 4 | | | | |
| Vesicle-mediated transport | 14 | 3 | 1 | | 3 | 2 | 7 | 7 | 24 | 3 | | | | 1 |

[1]Biological processes were annotated according to terms described by Gene Ontology Consortium.
[2]The numbers represent number of plant proteins in a biological process that are associated with each effector family.

**Table S3.** N-terminally GFP-tagged PexRD12/31 effectors coimmunoprecipitate with a largely overlapping set of *Nicotiana benthamiana* vesicle trafficking proteins compared to FLAG-tagged effectors

| *N. benthamiana* protein annotation | Effectors used[1] | | | | | | | | | | | | | | | | | | Sequence identifier |
|---|---|---|---|---|---|---|---|---|---|---|---|---|---|---|---|---|---|---|---|
| | Anti-GFP IPs | | | | | | Anti-FLAG IPs | | | | | | | | | | | | |
| | GFP-PITG_16245 | GFP-PITG_16242 | GFP-PITG_23074 | GFP-PITG_16428 | Total no. peptides | Other GFP fusions[2] | FLAG-PITG_16245 | FLAG-PITG_16233 | FLAG-PITG_16235 | FLAG-PITG_16242 | FLAG-PITG_16409 | FLAG-PITG_16243 | FLAG-PITG 23069 | FLAG-PITG_23074 | FLAG-PITG_16428 | Total no. peptides | Other FLAG fusions[3] | No. effector families associated[4] | |
| Vesicle fusing ATPase[5] | | | 1 | 4 | 5 | | | | 5 | 4 | 3 | 1 | 2 | 2 | 10 | 27 | 4 | 2 | NbS00011575g0012 |
| Transmembrane emp24 domain-containing protein A[5] | | | | 6 | 6 | | | | | | 2 | | | | | 2 | | 1 | NICBE_138070 |
| Dynamin 2B | | 2 | 4 | 1 | 7 | 16 | 7 | | | 1 | | | | | | 8 | 7 | 5 | NICBE_074039 |
| Syntaxin[5] | | | 13 | 6 | 19 | | | | | | | | | | 5 | 5 | 2 | 1 | NbS00027157g0003 |
| ADP-ribosylation factor 2[5] | 9 | | 6 | 4 | 21 | 12 | | | | | | | | | 2 | 2 | 11 | 2 | NICBE_020747 |
| Vacuolar-sorting receptor 3[5] | | | 9 | 13 | 22 | | | | 1 | 2 | | | | | 2 | 5 | | 1 | NICBE_083936 |
| Putative phagocytic receptor 1b[5] | | 1 | 13 | 9 | 23 | | | | | | | | | | 1 | 1 | 9 | 3 | NICBE_351046 |
| Ras-related protein RABA1f[5] | | 1 | 9 | 16 | 26 | | | | | | | | | | 4 | 4 | 4 | 2 | NICBE_128666 |
| Vesicle associated membrane protein 7B[5] | | 5 | 2 | 19 | 26 | | | | | 1 | | | | | 2 | 3 | 4 | 2 | NbS00022342g0004 |
| Extended synaptotagmin-3[5] | | | 9 | 19 | 28 | | | | | | | | | | 1 | 1 | 2 | 1 | NICBE_418648 |
| Exocyst complex component 5[5] | | 15 | 21 | 2 | 38 | | 4 | | | | | | | | | 4 | | 1 | NICBE_108112 |
| Sec7 guanine nucleotide exchange factor[5] | | 7 | 21 | 12 | 40 | | | | | | | | | | 2 | 2 | 2 | 2 | NbS00049277g0005 |
| Ras related protein Rab 2 A[5] | | 6 | 34 | 5 | 45 | | | | 1 | 1 | | | 1 | | 2 | 5 | 9 | 3 | NbS00004361g0010 |
| Coatomer subunit delta[5] | 1 | 9 | 18 | 25 | 53 | | | | | | | | 2 | | 1 | 3 | 4 | 1 | NICBE_222296 |
| Coatomer alpha subunit protein | 7 | 9 | 19 | 27 | 62 | | 5 | | | | | | | | 3 | 8 | 4 | 5 | NbS00003584g0003 |
| Exocyst complex component 3[5] | 1 | 5 | 29 | 35 | 70 | | 2 | | | | | | | | | 2 | | 1 | NICBE_298622 |
| Probable exocyst complex component 4[5] | | 11 | 39 | 43 | 93 | | 2 | | | | | | 1 | 1 | | 4 | 2 | 1 | NICBE_251150 |
| Exocyst complex component SEC3A[5] | | 14 | 46 | | 105 | | 2 | | | | | | | | | 2 | | 1 | NICBE_053349 |
| Coatomer subunit alpha-1[5] | 15 | 33 | 49 | 36 | 133 | 23 | 7 | | | | | | | | 3 | 10 | 9 | 4 | NICBE_286236 |
| Coatomer subunit beta-2[5] | 16 | 24 | 43 | 53 | 136 | 21 | 1 | | | | | | | | 1 | 2 | 11 | 4 | NICBE_204349 |
| Coatomer subunit gamma[5] | 3 | 23 | 71 | 72 | 169 | 16 | 13 | 5 | 1 | | | | 2 | | 7 | 28 | 7 | 3 | NbS00000812g0012 |
| ARF guanine nucleotide exchange factor 2[5] | | 52 | 88 | 57 | 197 | | 1 | | | | | | | | | 1 | 2 | 1 | NbS00006288g0001 |
| Pattern formation protein EMB30[5] | 1 | 14<br>5 | 42 | 23 | 211 | | 2 | | | | | | | 1 | 2 | 5 | | 1 | NICBE_346430 |
| | 12 | 40 | 92 | 16<br>4 | 308 | 14 | 8 | | | | | | 1 | | 2 | 11 | 7 | 5 | NICBE_323561 |
| SEC1 family transport protein SLY1[5] | | | | | | | 3 | | | | | | | | | 3 | | 1 | NICBE_170643 |
| SNAP25 homologous protein SNAP33 | | | | | | | | | | | | | 1 | | 2 | 3 | 1 | 1 | NICBE_352850 |
| Annexin D1[5] | | | | | | 4 | 4 | | | | | | 3 | | | 7 | 9 | 2 | NICBE_369402 |
| Dynamin related protein 1E[5] | | | | | | 13 | 1 | | | | | | | | | 1 | 6 | 2 | NbS00056353g0008 |
| Ras-related protein RABE1c[5] (Rab8a) | | | | | | 12 | | | 1 | 1 | 1 | 1 | 2 | 1 | 1 | 8 | 15 | 4 | NICBE_214722 |
| CASP protein | | | | | | | 6 | 1 | | | | | | | 6 | 13 | 19 | 6 | NICBE_048655 |

[1]Numbers shown in the table refer to the number of peptide hits against *N. benthamiana* protein identified by coIP/MS of effectors.
[2]Other anti-GFP coIP/MS datasets extracted from Petre *et al*., 2015, 2016 (negative controls). In total, 36 experiments are included.
[3]Other anti-FLAG coIP/MS datasets from this study (negative controls).
[4]Number of associated effector families was acquired from Dataset 1 and 2.
[5]Plant protein that shows association with less than five effector families.

**Table S4**. Primers used in this study

| Primer ID | Primer sequence (5’ to 3’) |
|---|---|
| **PITG16242F** | CACCGAAGACACAATGTGGCTTAGTCGTGTTACAAATTG |
| **PITG16242R** | CTACGAAGACGGAAGCTTAGTGTTTGTTCCGCCATTCGG |
| **PITG16428F** | CACCGAAGACACAATGCTGATCAAAACCTTCGAGTC |
| **PITG16428R** | CTACGAAGACGGAAGCTTAGAAACGTTTTAGTCGGTTGTTGC |
| **PITG23074F** | CACCGAAGACACAATGATTTTTCGCCGATTTACGGATTG |
| **PITG23074R** | CTACGAAGACGGAAGCTTATAGTTTGTCCCGCCACTTTTTCG |
| **PITG16245F** | CACCGAAGACACAATGCTGCTTAATGGTATGACAG |
| **PITG16245R** | CTACGAAGACGGAAGCTTACTGGTTCTTCCACCACTTC |
| **NbVAMP72x_F** | CACCGGTCAGCAGAAGGCTTTGATCT |
| **NbVAMP72x_R** | TTACTTTCCACAATTGAATCCCT |

**Table S5.** Coding nucleotide sequences of the fusion proteins used in this study

| Protein insert identifier | GenBank Accession | Coding sequence of insert |
|---|---|---|
| Inserts for GoldenGate plasmid vectors[1] | | |
| PITG23074$_{66-119}$ | XP_002897462 | A**ATG**ATTTTTCGCCGATTTACGGATTGGATCAAGTACCTTTTTAACAAGATGAATC<br>CGAAACAGTTACACATCTACTTGGGCTTAGATGGCCTTGGCGAAACGGCCTATC<br>AGCACAAGAACTACCCGATCTATCTGATGAAGTCGAAAAAGTGGCGGGACAAAC<br>TA**TAA**GCTT |
| PITG16245$_{61-115}$ | XP_002897644 | A**ATG**CTGCTTAATGGTATGACAGATTTTTTCAAGTACCACGCTGGAAAGATGAAT<br>CCCGAGCAGCTTTACAAGTACTTAAACTTAAAAGGACTTGGTCAAGAAGCCTACA<br>AACACAAGAACTACGCTAGTTACATTAAGAAGTCGAAGAAGTGGTGGAAGAACC<br>AG**TAA**GCTT |
| PITG16428$_{74-142}$ | XP_002897468 | A**ATG**CTGATCAAAACCTTCGAGTCTTGGGTGAGGTACAAGCTATTGCAGCTACA<br>CCCCAAATTCAGGCAAGTCAAGCTACATCCCGGTCGAGTCTTTACGCTCTTAGG<br>CTTAGATGGACTTCACGAAATGGCCATCTATCACCCTGGCTGGAAGAGGTACGT<br>GGACTACTCGCAGAAGTGGCGAAGCAACAACCGACTAAAACGTTTC**TAA**GCTT |
| PITG16242$_{75-129}$ | XP_002897642 | A**ATG**TGGCTTAGTCGTGTTACAAATTGGATCAAGTACAAGAGGGGCAAGATGAA<br>TCCCAAGCAATTACACACATACTTGGGCTTAGATGGACTGGGCCAAAGCGCTCG<br>CGATAGCTCGAACTTCCAGAAGTACTTGAAGAAGTCGGCAGAATGGCGGAACAA<br>ACAC**TAA**GCTT |
| AtEXO70E2 (At5g61010) | NP_200909 | A**ATG**GCAGAGTTTGATTCCAAGGTTCCTGTTTCCGGAATGAACAATCATGTCTTT<br>GAGGCATGTCACCATGTTGTTAAGGCACTGAGGGCATCTGATAACAACCTGGAT<br>GCCAATTTGAGAAAGCTCCTAAGTGACCTAGAGATGCATTTGTCAACATTTGGGA<br>TTGCTGATACCAAAGTTGAAGATGCAGGATTCTCTGAGATCAAGAAGAGATTCAA<br>AGAAGCTGTGAAGAGAATCCGCAGTTGGGAGACAAACCAGTCGACGATGTTTGA<br>AGCTGGTCTGTCTGAAGCTGATCAGTTCTTTCAAGCCCTGTATGACGTTCAAACA<br>GTTCTTGTGGGTTTCAAAGCTTTGCCCATGAAAACTAACCAGATGGAGAAAGATG<br>TTTACAACCAAGCTACGGTTGCTCTTGACATCGCGATGTTGAGGCTTGAGAAAG<br>AGCTCTGTGATGTTCTGCATCAGCACAAACGGCATGTACAGCCTGATTACTTGG<br>CTGTTAGTTCCCGTAGAAAAGATATTGTGTACGACGAGTCCTTTGTCTCCCTAGA<br>TGATGAAGTTATTGTCGAAGCTTCTTCACATGAAGATGATGAACAGATATCGGAC<br>TTTTATAATTCTGATTTGGTTGATCCCATCGTACTTCCTCACATCAAAGCTATTGC<br>AAACGCCATGTTTGCTTGCGAGTATGATCAACCATTCTGTGAAGCTTTCATCGGC<br>GTTCAAAGAGAAGCTCTGGAGGAGTATATGGTTACTCTCGAAATGGAAAGATTC<br>AGCTGCGTTGATGTTCTCAGAATGGACTGGGAGGATTTGAATGGTGCAATGAGA<br>AAATGGACAAAGGTTGTTAAGATCATTACTCAGGTCTATCTCGCCAGTGAAAAAC<br>AGCTATGTGATCAGATTTTGGGAGACTTTGAGTCAATTTCTACAGCTTGCTTCAT<br>TGAAATCTCAAAAGATGCAATTCTATCACTCCTCAACTTTGGAGAAGCTGTGGTG<br>CTGAGATCTTGCAAGCCAGAAATGCTTGAGCGCTTCCTTAGTATGTATGAGGTTT<br>CAGCAGAGATTCTCGTGGATGTCGATAACCTCTTCCCCGATGAAACAGGCTCGT<br>CTTTAAGAATTGCATTTCACAACCTGTCAAAAAAACTAGCTGATCATACAACCACA<br>ACCTTCTTAAAGTTCAAAGACGCAATAGCCTCAGATGAATCCACACGCCCCTTTC<br>ATGGAGGCGGGATTCATCACCTGACCAGGTATGTAATGAACTACTTGAAGCTTC<br>TCCCGGAGTATACCGACTCACTAAACTCACTGCTTCAGAACATACACGTTGACG<br>ACTCCATCCCCGAGAAAACAGGAGAAGATGTCCTCCCTTCGACATTCTCTCCAA<br>TGGCTAGACACCTCAGGTCAATCGTCACAACTCTAGAATCCAGCCTTGAGCGAA<br>AAGCTCAGCTGTACGCAGACGAAGCACTGAAATCCATTTTCCTGATGAACAACTT<br>CCGTTACATGGTCCAGAAGGTAAAGGGATCAGAGCTGAGACGCTTGTTTGGAGA<br>CGAATGGATCAGGAAGCACATTGCAAGCTACCAATGCAATGTCACTAACTACGA<br>GAGATCCACATGGAGCTCCATACTCGCTTTGCTCAGAGACAACAACGACTCTGT<br>AAGAACCCTGAGAGAAAGATGCAGACTTTTCAGCCTTGCATTTGATGATGTTTAC<br>AAGAACCAAACCCGTTGGTCAGTCCCTGATTCAGAGCTCCGCGATGATCTTCAC<br>ATCTCGACCTCTGTCAAGGTTGTTCAGTCTTACAGAGGATTTCTTGGAAGAAATG<br>CAGTTCGCATAGGCGAAAAACACATCCGATACACTTGTGAGGACATTGAGAATA<br>TGCTCCTTGATCTCTTTGAATGTTTGCCATCTCCAAGATCACTGCGCAGCTCTCG<br>TAAGAGAggTTCG |
| Inserts for pTRBO plasmid vector[2] | | |
| AVR2 | QBB68791 | ATGGACTACAAGGACGACGATGACAAAGTCAAGCTTCTCGAGAATTCCGGTTTT<br>TCTCTTAAGGATACTTTGAAGAAGCTTAATCCTATTAAAGCAGCTGTTAAAGCTAA<br>AGATAAGGCAAAAGAAGTTACTGAAAAAATTACAGACGCTGATTGGAAGAAACTT<br>GTGGAGCATTTGAAGATCAAAGGAGACAAAAGATCTTGA |
| PexRD11 | ACX46536 | ATGGACTACAAGGACGACGATGACAAAGTCAAGCTTCTCGAGAATTCCCTTCCT<br>GCAGATGCGGGTAAAGTCATTCATGAAAACGGTGCTGACACTCGTATTCCGACC<br>CACGTGCATGACCAGCGCTTACTGCGTCGTGTCAGAAATGACGAAGGTGAGCT<br>CACGGAAGAAAGAACCGGAGGCTTACTGGATAAGATAAAGTCTGTGGTGAAGAA<br>AATCACACCCGAAAAGGCCGTGACAAAGTTTAAGGAGAAGGACATTACGAATCC<br>CGAATGGTTGAAGATCATAAAGCACAAGGTTCGTGAAGCAAAGGGACAGGGGTA<br>CAAATAGGCGGCCGC |

| | | |
|---|---|---|
| PexRD11sh | ACX46536 | ATGGACTACAAGGACGACGATGACAAAGTCAAGCTTCTCGAGAATTCCACCGGA<br>GGCTTACTGGATAAGATAAAGTCTGTGGTGAAGAAAATCACACCCGAAAAGGCC<br>GTGACAAAGTTTAAGGAGAAGGACATTACGAATCCCGAATGGTTGAAGATCATA<br>AAGCACAAGGTTCGTGAAGCAAAGGGACAGGGGTACAAATGA |
| PITG_05121 | XP_002998822 | ATGGACTACAAGGACGACGATGACAAAGTCAAGCTTCTCGAGAATTCCATTGTTT<br>CTTGGCCTGAGGTTCTTAAGAAGGTTACTGCTGCTAGATCTGCAAAGATGAAGG<br>TGAGAAAAGTTACTGAAAAATTCGAGGATCCTGTGTCCAAGAAGCTTGTTGAACA<br>TTTGAGGGTGTATAGGGATAAGGGTTCTTGA |
| PITG_06077 | XP_002998270 | ATGGACTACAAGGACGACGATGACAAAGTCAAGCTTCTCGAGAATTCCACTATT<br>GGTAATGCAATTCTTGAAGCTGCAAAGAAGTTGGATCCTGTTGAAGCTGTTAAAA<br>AGGCTAAAGAAGCAGCCAAGAGAAAAAAGTCTATCTTGGAGACTATGAAACTTC<br>AAGCTTGGCTTGAGAAGATGAGAGAGACAATAATGAAAGATTGA |
| PITG_07500 | XP_002904502 | ATGGACTACAAGGACGACGATGACAAAGTCAAGCTTCTCGAGAATTCCGGTTTT<br>GGTGGAGCTCTCGTTGATGGTGTTAAGAAACTTAATCCTATTACTGCAGCTAAGA<br>AAGCAAAGGAGAAAGCCGAAAAGATTAAACAACATGTGAAGGAGCTTGGAAAAT<br>ATGAAGATTGGTTGAAGGAAGTTAGAGAAGCTATCGATAAAGACTGA |
| PITG_08278 | XP_002903684 | ATGGACTACAAGGACGACGATGACAAAGTCAAGCTTCTCGAGAATTCCGGTTTC<br>GGAGGTGCACTCGCTGATGGACTTAAGAAACCTAATCCTGCAAAGGCTGCTAAA<br>AAGGCCAAAGAAAAGGCTGCTAAAATTAAGCAAGATCTTAAAGAGATTGGCGAA<br>CATGCTGCTTGGTTGGAAAAGATGAGAGAGACTATCGGTAAGGACTGA |
| PITG_13936 | XP_002899599 | ATGGACTACAAGGACGACGATGACAAAGTCAAGCTTCTCGAGAATTCCACTGGT<br>GGTTTTATGGATAAGCTTAAATCTGTTGCTGAGAAGATTAAACCTTCTAAGGCTG<br>TTGAAAAAGTGAAGGAAATGGTTGTGACTAAACCTGAGTGA |
| PITG_13940 | XP_002899603 | ATGGACTACAAGGACGACGATGACAAAGTCAAGCTTCTCGAGAATTCCACTAAG<br>GGTTTTAAAGAAAAGCTTAAAGCTGTTCTTGAAAAGATTACTCCTAAAAAGTCTCT<br>TGACAAGCTTAAAGAAACAGTTGTGACTACTTTGGATTGGGTTAGAATTGTGAAG<br>CATAAAACCGATGAGGCTAAGAGATTGGGATTCAAATGA |
| PITG_15972 | XP_002898185 | ATGGACTACAAGGACGACGATGACAAAGTCAAGCTTCTCGAGAATTCCGGTTTT<br>GGTGGAGCTCTTGCAGATGGTCTTAAGAAGTTGAATCCTGCAAAAGCTGCTAAG<br>AAAGCCAAGGAAAAATCTTGTGAGAATTGA |
| PITG_19617 | XP_002996938 | ATGGACTACAAGGACGACGATGACAAAGTCAAGCTTCTCGAGAATTCCGGTTTC<br>GGTGGAGCTCTTGCAGATGGACTTAAGAAATTGAATCCTGCTAAGGCAGCCAAA<br>AAGGCTAAAGAAAAGGCTGCCAAAATTAAGCAAGATTTGAAGGAAATTGGTGAG<br>CATGCTGCTTGGCTTGAGAAGATGAGAGAAACAATCGGAAAGGACTGA |
| PITG_21949 | XP_002996876 | ATGGACTACAAGGACGACGATGACAAAGTCAAGCTTCTCGAGAATTCCACTAAG<br>GGTTTCAAGGAAAAACTTAAGGCTGTTGTTGAAAAGATTACTCCTAAGAAGTCTC<br>TTGATAAGCTTAAGGAGACAGTGGTTACTACCCTCGATTGGGTGAAGATTGTTAA<br>GCATAAGACAGACGAGGCTAAGAGATTGGGATTTAAGTGA |
| PITG_22975 | XP_002899618 | ATGGACTACAAGGACGACGATGACAAAGTCAAGCTTCTCGAGAATTCCGCTAAG<br>GGTTTTAAAGAGAAGCTTAAAGCTGTTGTTGAAAAGATCACTCCTAAAAAGTCTT<br>TGGAAAAACTTAAGGAAACTGTGATTTCTACACCTGATTGGGCTAAGATCGTTAA<br>ACATAAGACTGATGAGGCAATGAGACTCGGATATAAATGA |
| AVR3a | E2DWQ7 | ATGGACTACAAGGACGACGATGACAAAGTCAAGCTTCTCGAGAATTCCATCGAC<br>CAAACCAAGGTCCTGGTGTATGGGACGCCAGCTCACTACATACACGATTCAGCC<br>GGCAGAAGACTTCTTCGCAAGAACGAAGAGAATGAAGAAACGTCTGAGGAGCG<br>AGCCCCAAATTTCAATTTGGCGAATCTAAATGAGGAGATGTTTAATGTGGCTGCG<br>TTGACGAAGAGAGCAGATGCCAAAAAGCTAGCGAAACAGCTTATGGGTAATGAT<br>AAGCTGGCGGATGCTGCATACATTTGGTGGCAGCACAACAGGGTTACGCTAGA<br>CCAGATTGACACGTTCCTGAAGCTTGCAAGCCGCAAGACGCAAGGCGCAAAGT<br>ACAATCAGATCTACAATAGCTACATGATGCACCTGGGGCTCACTGGATATTAG |
| Pex147-2 | XP_002898841 | ATGGACTACAAGGACGACGATGACAAAGTCAAGCTTCTCGAGAATTCCGTCGAC<br>CAAACCAAGGTTCTGATGTATGGGTCGCCAGCTCACTACATACACGATTCAGCC<br>GGCAGAAGACTTCTTCGCAAGAACGAAGAGAGTGAAGAAACGTCTGAGGAGCG<br>AGCCCCTAATTTCAATTTGGCGACTTTAAATGAGGAGATGTTTGATGTGGCTGCG<br>TTGACGAAGAAAGCAGATGCCAAAAAGCTAGCTAAACAGCTTATGGGTAATGGT<br>AAGCTGGCGGATGCTGCATACATTTGGTGGCAGCACAAGAGGCTTACGCTAGA<br>CCAGATTGACGCGTTCCTCAAGCTCGCAAGCAGCAAGACACAAGGCGCGAGAT<br>ACAATCGGATCTACAATAGCTGCTATATGATGCACCTGGGGCTCACTGGATTTTA<br>GGCGGCCGC |
| Pex147-3 | XP_002898843 | ATGGACTACAAGGACGACGATGACAAAGTCAAGCTTCTCGAGAATTCCATCGAC<br>CAAACCAAGGTTCTGATGTATGGGACGCCAGCTCACTACATACACGATTCGGCC<br>GGCAGAAGATTTCTTCGCAAGAACGAAGAGAATGAAGAAACGTCTGAGGAGCGA<br>GCCCCAAATTTCAATTTGGCGAATTTAAATGAGGAGATTTTTAATGTGGCTGCGT<br>TGACGAAGAAAGCAGATGCCAAAAAGCTAGCGAAACAGCTCATGGGTAATGATA<br>AGATGGCGAAAGCAGCATACGTTTGGTGGCAGCACAATGGTGTGACGCCAAGC<br>CAGATTGACACGTTCCTGAAGCTCGCAAGCGGCAAGACGCAAGGCGCAAGATA<br>CAATGAGATCTACAATAGCTACCTGATGCACCTGGGGCTCACTGCATATTAGGC<br>GGCCGC |
| AVR4 | XP_002904419 | ATGGACTACAAGGACGACGATGACAAAGTCAAGCTTCTCGAGAATTCCGCGAAA<br>GCTGATTCTTTAGCTCGTACCGTCAGCGTTGTTGACAACGTCAAAGTAAAAAGCA<br>GATTTCTGAGGGCTCAAACGGACGAGAAGAACGAAGAGAGAGCAACGATAACG<br>CTTGGAGACAGGGTTGTTTCCGACAAGGCGGCGACAAAAGATCTGCTACAGCA<br>GCTTCTTGCACTGGGCACGCCACTGGAAAAAGTCCAGAAGCAATTCCTGAACAT<br>ACCGCAGATGAAAACATTTGCGGAGTTGAGCAAACACCCGAACTGGAAAGCGCT<br>TGACAAATATGAACGGATGCAGTGGCAGAAGCTAAAGGAGGGCGAAACACTGA<br>CATTTATGCGTCTTGGCGATCGATTATACTCTAAAGAGAAAGCGCAAGAACAGCT |

| | | |
|---|---|---|
| | | CCTTAGGTGGGTTGCGCAGAAAAAACCTGTGGAGAGTGTATATGATGACCTACA<br>AGTGGCAGGCTTTGCACATAATACTGTTGCTGCTCGCCAGAACTGGAGAGCATA<br>TATTATGTACGACAAGTGGTTTACGGCGGCCTCACAAATGCAGAGGAACCCGCA<br>GCAGTATGCCAAGTTCGGCACGGGATATCATTCGGAGCAAAAGACGACGGAGT<br>TGTTCGAGAAGTGGGCCATGGAGGGAACCCATATAAAAAGTGTCATCACGACGC<br>TTAAACTCAACGGTAAGTCGGCGTCTGAGATGGCAAATAACGAGAATTTTCCCG<br>CGCTCCTGAAGTATGTCAAGTTGTATCTTGATTTTAAACCAGTCAGGGACCTTAA<br>CGCAAAATCCCGTCTCCAAGCTAGACGGCCCATATCTTAGGCGGCCGC |
| AVRblb1 | XP_002895051 | ATGGACTACAAGGACGACGATGACAAAGTCAAGCTTCTCGAGAATTCCGTTTCA<br>TCCAATCTCAACACCGCCGTGAATTACGCTTCCACATCCAAGATTCGCTTTCTGT<br>CGACTGAGTACAACGCCGATGAAAAAAGAAGCTTGCGAGGTGACTACAACAATG<br>AGGTCACAAAAGAGCCCAACACGTCTGACGAAGAGCGGGCGTTTTCTATCTCAA<br>AGTCTGCGGAATACGTGAAGATGGTACTTTATGGATTCAAACTTGGATTTTCTCC<br>TCGCACTCAGTCCAAGACGGTGTTGCGATACGAAGATAAACTGTTTACGGCTCT<br>CTATAAATCCGGAGAGACGCCGAGAAGCCTAAGGACCAAGCATCTCGATAAGG<br>CTTCCGCTAGCGTATTTTTCAACAGATTCAAAAAATGGTACGATAAAAACGTTGG<br>CCCTAGCTAGGCGGCCGC |
| PexRD39 | ACX46588 | ATGGACTACAAGGACGACGATGACAAAGTCAAGCTTCTCGAGAATTCCTTCCCA<br>ATCCCCGACGAGTCTCGCCCCTTGTCGAAGACATCTCCTGACACTGTGGCCCCA<br>AGATCGCTTCGGATCGAGGCCCAAGAAGTTATTCAGAGCGGCCGGGGAGACGG<br>ATATGGTGGGTTCTGGAAAAACGTAGCCCAGAGTACTAACAAGATCGTCAAGAG<br>GCCGGATATCAAGATAAGCAAACTTATCGCGGCGGCCAAGAAGGCAAAAGCAAA<br>AATGACGAAGTCCTGAGCGGCCGC |
| PexRD39sh | ACX46588 | ATGGACTACAAGGACGACGATGACAAAGTCAAGCTTCTCGAGAATTCCATTGAA<br>GCTCAAGAAGTTATCCAATCTGGTAGAGGTGATGGATATGGTGGATTTTGGAAG<br>AATGTTGCTCAATCTACTAATAAGATAGTGAAGAGACCTGATATTAAGATTTCTAA<br>ACTTATCGCTGCTGCTAAAAAGGCTAAAGCTAAGATGACTAAATCTTGA |
| PexRD40 | ACX46596 | ATGGACTACAAGGACGACGATGACAAAGTCAAGCTTCTCGAGAATTCCTTCCCA<br>ATCCCCGACGTGTCTCGCCCCTTGTCGAAGACATCTCCTGACACTGTGGCCCCA<br>AGATCGCTTCGGGTCGAGGCCCAAGAAGTTATTCAGAGCGGCCGGGGAGACGG<br>ATATGGAGGGTTCTGGAAAAACGTAGTCCAGAGTACTAACAAGATCGTCAAGAA<br>GCCGGATATCAAGATAGGCAAACTTATCGAGGCGGCCAAGAAGGCAAAAGCAA<br>AAATGACGAAGTCCTGAGCGGCCGC |
| PexRD40_34aa | ACX46596 | ATGGACTACAAGGACGACGATGACAAAGTCAAGCTTCTCGAGAATTCCGAAGCT<br>CAAGAAGTTATTCAATCTGGTAGAGGTGATGGTTATGGAGGTTTTTGGAAGAATG<br>TTGTGCAATCTACTAATAAGATTGTTAAGAAACCTGATATCTGA |
| PexRD40sh | ACX46596 | ATGGACTACAAGGACGACGATGACAAAGTCAAGCTTCTCGAGAATTCCGTTGAA<br>GCTCAAGAAGTTATTCAATCTGGAAGAGGTGATGGTTATGGAGGTTTTTGGAAG<br>AATGTGGTTCAATCTACTAATAAGATTGTGAAGAAACCTGATATCAAAATAGGTAA<br>GCTTATTGAGGCTGCAAAAAAGGCTAAAGCTAAGATGACTAAATCTTGA |
| PexRD41 | ACX46573 | ATGGACTACAAGGACGACGATGACAAAGTCAAGCTTCTCGAGAATTCCGCGTTT<br>CTGAATCCCGACGAAACTCGGCTCTTATCAGACACTTTTACCAAAAGATCCCTTC<br>GGGTCGCAGGCCAAGAAGTTGCCCGGGGCGACCGGGGCGAAGAGATTGTGAG<br>AGTTATAGTCCAGAGTACTAACAAAATCTTCAAGAGACCGGCGGAAAAAGACAT<br>GAGCAAACTGATTGCAGCGGCTAAGATTGCGATGTTGGAGAAAAAGATGGCTAA<br>GCTCTCATTCGTCGGTAAGGAGGCAGCGAAGTAGGCGGCCGC |
| PexRD45 | ACX46578 | ATGGACTACAAGGACGACGATGACAAAGTCAAGCTTCTCGAGAATTCCATCCCG<br>AATCACACCACCGAGTCTCAACTCTTGTCGAAGGCATCCCCTGACCCTGCGGCG<br>AAAAGATCTCTCCGGAACGCAGGCCAACAAGTTGTTCAGAGCCGCCCGACGGA<br>CGGGAACGGTGGAGTCTTCAAAGCTTTCAGTGGTACAAACAAACTCATAAAGCT<br>GCCAGACATGAAAATAAGCAACGTTCTCGAAGCTGCCAAGAAGGTGAAAAAGTT<br>GAAAGAGATGGACAAGTTGAAAAAGTTGATTAAGTCCTCAAAGTAGGCGGCCGC |
| PexRD46 | ACX46579 | ATGGACTACAAGGACGACGATGACAAAGTCAAGCTTCTCGAGAATTCCGCGCTT<br>CTGAATCCCGACGAAACTCGGCTCTTATCAGACACTTTTACCAAAAGATCCCTTC<br>GGGTCGCAGGCCAAGAAGTTGCCCGGGGCGACCGGGGCGAAGAGATTGTGAG<br>AGTTATAGTCCAGAGTACTAACAAAATCTTCAAGAGACCGGCGGAAAAAGACAT<br>GAGCAAACTGATTGCAGCGGCTAAGATTGCGATGTTGGAGAAAAAGATGGCTAA<br>GCTCTCATTCGTCGGTAAGAAGGCAGCGAAGTAGGCGGCCGC |
| PITG_04086 | XP_002905796 | ATGGACTACAAGGACGACGATGACAAAGTCAAGCTTCTCGAGAATTCCGTTGAA<br>GCTCAAGAAGTTATTCAATCTGGAAGAGGTGATGGTTATGGAGGTTTTTGGAAG<br>AATATTATCCCTTCTACTAATAAGATAATTAAGAAACCTGATATCAAGATTTCTAA<br>GCTTATAGAGGCTGCAAAAAAGGCTAAAAAGAAAATGACTAAGTCTTGA |
| PITG_04097 | XP_002905808 | ATGGACTACAAGGACGACGATGACAAAGTCAAGCTTCTCGAGAATTCCGTTGAA<br>GGTCAAGAAGTTGTGCAAGGTGGTACTCTTGATGGTAATGGTGGTGTTTGGAAG<br>GCTATTGCTCATACTACTAATAAGATTGTGAAGAAGCCTGAAATCGATGTTTCTA<br>AACTTATTGATGTGGCTAAGAAAGCTAAGAAGGTTAAGAAGCTTAAAAATCTTAT<br>GAAACTTAAGAAGTCTTCATCTTGA |
| PITG_04194 | XP_002905878 | ATGGACTACAAGGACGACGATGACAAAGTCAAGCTTCTCGAGAATTCCATTGCT<br>GGTCAAGAAGTTGTGCAATCTAGACCTCTTGATGGTAATGGTGGTGTTTGGAAAT<br>CAATCGTGCAAACTACTAATAAAGTTGCTAAGAATCCTGAAATAGATCTTTCTGAT<br>CTTGTGAAAGTTGCTAAGAAAGCTAAGGATCTTAAGAAGATTAAGAATCTTATTTC<br>AAGAAAGAAAACTTCTTCTTGA |
| PITG_18675 | XP_002997305 | ATGGACTACAAGGACGACGATGACAAAGTCAAGCTTCTCGAGAATTCCGTTGAA<br>GGTCAAGAAGTGGTTCAATCTGGTAAGCTTGATGGTAAAGGTGGTGTTTGGAAG<br>ACTATTACTCATACTACTAATAAAATTGTTCCTAAGCCTGATATCGATGTGTCTAA |

| | | |
|---|---|---|
| | | AGTTCTTGATCTTGCTAAGAAAGTTATGGAAGCTGAAAGACTTAAGAAACTTATTA<br>AGCTTAAGAAGTCTTCATCCTGA |
| PITG_18683 | XP_002997312 | ATGGACTACAAGGACGACGATGACAAAGTCAAGCTTCTCGAGAATTCCGTTGAA<br>GCTCAAGAAGTTATTCAATCTGGTAGAGGTGATGGTTATGGAGGTTTTTGGAAGA<br>ATATTATTCCTTCTACTAATAAGATTATCAAAAAGCCTGATATCAAAATTGGAAAG<br>CTTATAGAGGCTGCAAAGAAGGCTAAAAAGAAAATGACTAAATCTTGA |
| PITG_20300 | XP_002895918 | ATGGACTACAAGGACGACGATGACAAAGTCAAGCTTCTCGAGAATTCCATTGAA<br>GCTCAAGAAGTTATTCAATCTGGTAGAGGTGATGGATATGGTGGATTTTGGAAG<br>AATGTTGCTCAATCTACTAATAAGATAGTGAAGAGACCTGATATCAAAATTGGTA<br>AACTTATTGAGGCAGCTAAGAAGGCTAAAGCAAAGATGACTAAATCTTGA |
| PITG_20301 | XP_002895919 | ATGGACTACAAGGACGACGATGACAAAGTCAAGCTTCTCGAGAATTCCGTTGAA<br>GCTCAAGAAGTTATTCAATCTGGTAGAGGTGATGGTTATGGAGGTTTTTGGAAGA<br>ATGTGTTTCCTTCTACTAATAAGATTATCAAGAAACCTGATATTAAGATATCTAAA<br>CTTATTGCTGCTGCAAAAAAGGCTAAAGCTAAGATGACTAAATCTTGA |
| PITG_20303 | XP_002895919 | ATGGACTACAAGGACGACGATGACAAAGTCAAGCTTCTCGAGAATTCCGTTGAA<br>GCTCAAGAAGTTATTCAATCTGGTAGAGGTGATGGTTATGGAGGTTTTTGGAAGA<br>ATGTGTTTCCTTCTACTAATAAGATTATCAAAAAGCCTGATATTAAAATATCTAAG<br>CTTATTGCTGCAGCTAAAAAGGCTAAAGCTAAGATGACTAAATCTTGA |
| PITG_07689 | XP_002904634 | ATGGACTACAAGGACGACGATGACAAAGTCAAGCTTCTCGAGAATTCCAATGAC<br>ATTGTACTGACTGGTGTAATGTCGATGGGCGTTTTGCACCTCGTCAGTGCTGAC<br>CAAAGCACCATCGACTATGCTCGATTCCTGAGAGTGAACAAAGCTGTCGGTGAA<br>AACAACGAAGAGAGGGCTGGGGAAGGTGCTCAAAAGGCAAAGGCCCTTGTCAG<br>CCAGATGTTCAAGTCAAACTCTTTTTCAGCCCATGGACGAAAAGCTGAAATCGGT<br>GTTCAACTTCGCTAA |
| PITG_16275 | XP_002897665 | ATGGACTACAAGGACGACGATGACAAAGTCAAGCTTCTCGAGAATTCCACCGGG<br>ACCGACGTTTCACTAGCTGGTGTTTTGTCGCTGGGCTTGCTACACCTCGTCAGC<br>GCTGACCAAAGTTTTACTGAGCCGCAGCGGTTGCTGAGCGGCAACCACAAGGT<br>CGGCGAAGGTATTAAGGAAGAGAGAGGGTTCAAGGAAGTTGCCGAGGAGGCGA<br>TGGCAAAGTTTGTTGTCGACAAGGTGTATAGGACAAACTCTTTTTCGGCCCTCGA<br>GAAACTAGAGAAAACGCTCAATTTGCCGGTGCTCGAAAAAGCTTTGGTATTGGC<br>AGATGATAAAATGAAAGGGGCGTTCAAGTTCGCTGAGGAAGCGAAGATGAATCC<br>TGCAGATATGGCCAAAATTGTGAAAGATTTCCCAGATTTCACGGATGCTGAGAA<br>GATGAAGACAGTGCTAGAATACACCAACTACTTAAAGACAATGGGCAAATTGGA<br>CTAG |
| PITG_16294 | XP_002897362 | ATGGATTACAAAGATGATGATGATAAAGTGAAGTTGCTTGAAAATTCTACCTCTA<br>CACAGAACCTTGGAAATAGTTTGATGAGAGTGTTCTCTAAGGAAGCTACTAGAAA<br>ATATTACCTTGATTTGTTTAAGAGGGCTGATTTCACAGCAAATCTTCCTAAACTCG<br>CAAAGAAAGGTGGACCAGATAGACTCAACGATGCTCTTAAGAAATTGAGGAAGG<br>CAGGTATTTCAGAAGAGAAGTTCGCTGAGTTGAAAGGAGCTGCAGCTAAATACG<br>CAGATGATTGGTACAGAATCTACGGTAAACTCGATCCTATCAGGGCATAA |
| PITG_18880 | XP_002997157 | ATGGATTACAAGGATGATGATGATAAGGTGAAACTCTTGGAAAACTCTATCTCTA<br>TTAAAAACATTGGAAACTTCTTGAAAAGGATTTTCAGTAAGGAAGCTAGAAAGAA<br>ACATTATCTTAATAAGTTCAAACAAGCTGATATCGAGTCTAATCTTCCTAACTTGG<br>CAAAGAAAGGTGGACCAAATAGGTTGAACGATGCATTACAAAAGCTCAAGAAAA<br>CTGGTATCTCTGAAGAGCAGCTTAAAGAATTGGAGTCAGCTGCAAAAGGTTATA<br>GTAAGGAATGGTTCAGGAAGTTCGGAAAGTTAGATCCAGTTAGAGCATAG |
| PITG_18880-4 | XP_002997157 | ATGGATTACAAAGATGATGATGATAAGGTGAAACTCCTCGAAAACTCTATCTCTA<br>TTAAGAACATTGGAAACTTCCTCAAAAGAATTTTCAGTAAGGAAGCTAGAAAGAA<br>ACATTATCTTAAGAAGTTTAAGCAAGCTGATATTGAGTCTAATCTTCCTAACTTGG<br>CAAAGAAAGGTGGACCAAATAGGCTTAACGATGCATTGCACGTTCAAAAGTTGA<br>AGAAAACTGGTATCTCTGAAGAGCAGTTAAAAGAACTCGAGTCAGCTGCAGAGG<br>TTTATTCAAAAGAGTGGTTTAGGAAGTTCGGAAAGTTAGATCCAGTTAGGGCATA<br>G |
| NUK10 | XP_002898730 | ATGGACTACAAGGACGACGATGACAAAGTCAAGCTTCTCGAGAATTCCCTATCT<br>GCCCATCGGGCGCAGATAATGAACGTCGCGACGTCAGATCTCATCTCACCGATC<br>GAGTCTACAGTCCAAGACGACAACTACGACAGACAGTTGCGGGGGTTCTACGCT<br>ACAGAAAATACAGACCCTGTTAACAATCAAGACACTGCGCATGAGGATGGCGAG<br>GAGAGGGTCAATGTCGCCACGGTGCTTGGAAAGGGGGATGAAGCTTGGGACGA<br>TGCACTGATGCGCTTGGCCTATCAGCACTGGTTCGACGGAGGCAAAACTAGTGA<br>CGGCATGCGATTAATAATGGACCTTCCAGCGAAAGGTGAAGCACTCCGACACCC<br>GAATTGGGGGAAATACATTAAATACTTAGAGTTCGTTAAGGAGAAAAAGAAGGA<br>GGCTGCAGACGCTGCGGCAGTCGCGGCGCTCAAGCGAAGGCGGACTTACAGG<br>GGATGGTATGTCGACGGGAAAACGGAGAAAGACGTACGCAAGATTTTCGGACTT<br>CCGGCAACAGGAAAAGCCAAGAACCACCCAAACTGGGCAGATTTTCAGGAATAC<br>TTAAACGTCGTAAGAGAATACTCAAAAGTAGTTTTTAAATGAGCGGCCGC |
| NUK10-NLS | XP_002898730 | ATGGACTACAAGGACGACGATGACAAAGTCAAGCTTCTCGAGAATTCCCTATCT<br>GCCCATCGGGCGCAGATAATGAACGTCGCGACGTCAGATCTCATCTCACCGATC<br>GAGTCTACAGTCCAAGACGACAACTACGACAGACAGTTGCGGGGGTTCTACGCT<br>ACAGAAAATACAGACCCTGTTAACAATCAAGACACTGCGCATGAGGATGGCGAG<br>GAGAGGGTCAATGTCGCCACGGTGCTTGGAAAGGGGGATGAAGCTTGGGACGA<br>TGCACTGATGCGCTTGGCCTATCAGCACTGGTTCGACGGAGGCAAAACTAGTGA<br>CGGCATGCGATTAATAATGGACCTTCCAGCGAAAGGTGAAGCACTCCGACACCC<br>GAATTGGGGGAAATACATTAAATACTTAGAGTTCGTTAAGGAGACTTACAGGGG<br>ATGGTATGTCGACGGGAAAACGGAGAAAGACGTACGCAAGATTTTCGGACTTCC<br>GGCAACAGGAAAAGCCAAGAACCACCCAAACTGGGCAGATTTTCAGGAATACTT<br>AAACGTCGTAAGAGAATACTCAAAAGTAGTTTTTAAATGAGCGGCCGC |

| | | |
|---|---|---|
| PexRD12B | XP_002895188 | ATGGACTACAAGGACGACGATGACAAAGTCAAGCTTCTCGAGAATTCCCTTCTTA<br>ATGGTATGACTGATTTTTTTAAGTATCATGCTGGTAAGATGTCTCCTGAACAACTT<br>TATAAGTATCTTAATCTTAAGGGTCTTGGTCAAGAAGCTTATAAGCATAAGAATTA<br>TGCTTCTTATATTAAGAAGTCTAAGAAGTGGTGGAAGAATCAATGA |
| PexRD31 | XP_002897647 | ATGGACTACAAGGACGACGATGACAAAGTCAAGCTTCTCGAGAATTCCAGGCAA<br>ACAGCAGCGAATATTATGTACCCAGTTCTCGATGGAGAACAGAACGTCCTTGGC<br>AAGCGATCCCTAAGAACCGACCACATGAGAGTTAGCAATGTTGAGGACCGAGAA<br>GGTGATGAAGAACGAATTTTTCGCCGATTTACGGATTGGATCAAGTACATTTTTA<br>ACAAGATGAATCCGAAACAGTTACACACCTACTTGGGCTTAGATGGCCTTGGCG<br>AAACGGCCTATCAGCACAAGAACTACCCGATCTATCTGATGAAGTCGAAAAAGT<br>GGCGGGACAAACTATAGGCGGCCGC |
| PITG_16233 | XP_002897640 | ATGGACTACAAGGACGACGATGACAAAGTCAAGCTTCTCGAGAATTCCCTTAAT<br>GGTATGACAGATTTTTTCAAGTACCACGCTGGAAAGATGAGTCCCGAGCAGCTT<br>TACAAGTACTTAAACTTAAAAGGACTTGGTCAAGAAGCCTACAAACACAAGAACT<br>ACGCTAGTTACATTAAGAAGTCGAAGAAGTGGTGGAAGAACCAGTAA |
| PITG_16235 | XP_002897637 | ATGGACTACAAGGACGACGATGACAAAGTCAAGCTTCTCGAGAATTCCCTTAGT<br>CGTGTTACAAATTGGATCAAATACAAGAGGGGCAAGATGAATCCCAAGCAATTA<br>CACACATACTTGGGCTTAGATGGACTGGGCCAAAGCGCTCGCGATAGCTCGAA<br>CTTCCAGAAGTACTTGAAGAAGTCGGCAGAATGGCGGAACAAACACTAA |
| PITG_16242 | XP_002897642 | ATGGACTACAAGGACGACGATGACAAAGTCAAGCTTCTCGAGAATTCCTTGACG<br>ACGAAGGCTAACGCCGACACGACGCAGACGACAACCGATATTTCGCCCGCATT<br>GCTCGCAGGAGTACAGGACAAAGACCTTGCCATGCGAGCTCTTAGGACAGGTC<br>GCTCGAGTGGTGGCAATGTTGAGCACAAGGACCATGACGAAGAGCGGTGGCTT<br>AGTCGTGTTACAAATTGGATCAAGTACAAGAGGGGCAAGATGAATCCCAAGCAA<br>TTACACACATACTTGGGCTTAGATGGACTGGGCCAAAGCGCTCGCGATAGCTCG<br>AACTTCCAGAAGTACTTGAAGAAGTCGGCCGAATGGCGGAACAAACACTAA |
| PITG_16243 | XP_002897643 | ATGGACTACAAGGACGACGATGACAAAGTCAAGCTTCTCGAGAATTCCATTTTTA<br>GAAGATTTACTGATTGGATTAAGTATCTTTTTAATAAGATGAATCCTAAGCAACTT<br>CATACTTATCTTGGTCTTGATGGTCTTGGTGAAACTGCTTATAAGCATAAGTCTTA<br>TCCTATTTATCTTATGAAGTCTAAGAAGTGGAGAGATAAGCTTTGA |
| PITG_16248 | XP_002897647 | ATGGACTACAAGGACGACGATGACAAAGTCAAGCTTCTCGAGAATTCCATTTTTA<br>GAAGATTTACTGATTGGATTAAGTATATTTTTAATAAGATGAATCCTAAGCAACTT<br>CATACTTATCTTGGTCTTGATGGTCTTGGTGAAACTGCTTATCAACATAAGAATTA<br>TCCTATTTATCTTATGAAGTCTAAGAAGTGGAGAGATAAGCTTTGA |
| PITG_16409 | XP_002897637 | ATGGACTACAAGGACGACGATGACAAAGTCAAGCTTCTCGAGAATTCCTGGCTT<br>TCTAGAGTTACTAATTGGATTAAGTATAAGAGAGGTAAGATGAATCCTAAGCAAC<br>TTCATACTTATCTTGGTCTTGATGGTCTTGGTCAATCTGCTAGAGATTCTTCTAAT<br>TTTCAAAAGTATCTTAAGAAGTCTGCTGAATGGAGAAATAAGCATTGA |
| PITG_16427 | XP_002897644 | ATGGACTACAAGGACGACGATGACAAAGTCAAGCTTCTCGAGAATTCCCTTCTTA<br>ATGGTATGACTGATTTTGTTAAGTATCATGCTGGTAAGATGAATCCTGAACAATT<br>GTATAAGTATCTTAAGTTGCAAGGTAGAGGTCAAGAAGCTTATAAGCATAAGAAT<br>TATGCTTCTTATATTAAGAAGTCTAAGAAGTGGTGGAAGAATCAATGA |
| PITG_16428 | XP_002897468 | ATGGACTACAAGGACGACGATGACAAAGTCAAGCTTCTCGAGAATTCCCTTATTA<br>AGACTTTTGAATCTTGGGTTAGATATAAGCTTCTTCAACTTCATCCTAAGTTTAGA<br>CAAGTTAAGCTTCATCCTGGTAGAGTTTTTACTCTTCTTGGTCTTGATGGTCTTCA<br>TGAAATGGCTATTTATCATCCTGGTTGGAAGAGATATGTTGATTATTCTCAAAAGT<br>GGAGATCTAATAATAGACTTAAGAGATTTTGA |
| PITG_20336 | XP_002897640 | ATGGACTACAAGGACGACGATGACAAAGTCAAGCTTCTCGAGAATTCCTTGACC<br>ACGACTGTGGCTGACACGGCCCAGACGGCAACCAGCATTCTAACTCCTGTTCTA<br>GCTGGGGAGCCGAACAAACACGTTGCAACGCGATCTTTGAGAACGCATCCGAT<br>AGACGACAGCGACGATGGCGAAGAGCGACTGCTTAATGGTATGACAGATTTTTT<br>CAAGTACCACGCTGGAAAGATTAA |
| PITG_23069 | XP_002897638 | ATGGACTACAAGGACGACGATGACAAAGTCAAGCTTCTCGAGAATTCCATTTTTA<br>GAAGATTTACTGATTGGATTAAGTATCTTTTTAATAAGATGAATCCTAAGCAACTT<br>CATACTTATCTTGGTCTTGATGGTCTTGGTGAAACTGCTTATAAGCATAAGAATTA<br>TCCTATTTATAGAATGATGTCTAAGAAGTGGAGAGATAAGCTTTGA |
| PITG_23074 | XP_002897462 | ATGGACTACAAGGACGACGATGACAAAGTCAAGCTTCTCGAGAATTCCATTTTTA<br>GAAGATTTACTGATTGGATTAAGTATCTTTTTAATAAGATGAATCCTAAGCAACTT<br>CATATTTATCTTGGTCTTGATGGTCTTGGTGAAACTGCTTATCAACATAAGAATTA<br>TCCTATTTATCTTATGAAGTCTAAGAAGTGGAGAGATAAGCTTTGA |
| PITG_05911 | XP_002998153 | ATGGACTACAAGGACGACGATGACAAAGTCAAGCTTCTCGAGAATTCCGGTTAT<br>CATCTTACTCTCAAGGGTGAAGTTAAAGCTCTCGCTAAGAAAATTATTGCTGATT<br>TTAACACTGCTGATGATGTTTATAAGAAATGGAACGAAAATGGTCATTCTCTTAAT<br>AAGATTGCTAATCTCCTTAAAGTTTCTGAAAAGAAGAAATATGCTCCTGTTTACAA<br>TGGTTACCTTGCTTATCTTAACAGAATTAATTCTTGA |
| PITG_05912 | XP_002998154 | ATGGACTACAAGGACGACGATGACAAAGTCAAGCTTCTCGAGAATTCCGGTTAT<br>CATCTTACTCTCAAGGGTGAAGTTAAGGCTCTCGCTAAAAAGATTATTGCTGATT<br>TTAATACTGCTGATGATGTTTACAAAAAGTGGAACGAAAATGGTCATTCTCTTAAC<br>AAAATTGCTAACCTCCTTAAGGTTTCTGAAAAGGAAAAATACGCTCCTGTTTATAA<br>TGGTTATCTTGCTTATCTTAATAGAATTAATTCTTGA |
| PITG_05918 | XP_002998156 | ATGGACTACAAGGACGACGATGACAAAGTCAAGCTTCTCGAGAATTCCGGTTTT<br>CATCTTACTACTAGAGCTGAAGTTAAGGCTCTCGCTAGAAAAGTTATTACTGATT<br>TTGATGATGCTAATGATGTTTATAGAAAATGGTACGAAAACGGTTACTCTAAGAA<br>CAAAATTGCTAATCTCCTCAAGGTTTCTGAAAAAGAAAAATATGCTCCTGTTTATA<br>ACGGTTATCTTGCTTATCTTAATAGAATTAATTCTTGA |

| | | |
|---|---|---|
| PITG_22089 | XP_002894699 | ATGGACTACAAGGACGACGATGACAAAGTCAAGCTTCTCGAGAATTCCGGTTAT<br>CATCTTACTCTCAAGGGTGAAGTTAAAGCTCTTGCTAAGAAAATTATTGCTGATTT<br>TGATACTGCTGATGATGTTTATAAAAAGTGGAATGAAAACGGTCATTCTCTCAAC<br>AAAATTGCTAATCTCCTTAAGGTTTCTGAAAAGGAAAAGTACGTTCCTGTTTACAA<br>CGGTTATCTTGCTTATCTTAATAGAATTAATTCTTGA |
| PexRD2 | XP_002894954 | ATGGACTACAAGGACGACGATGACAAAGTCAAGCTTCTCGAGAATTCCCTCTCG<br>ACGAACACGGGTGTTCAGGCTGCTAATTTGGTAGGCCCAGCCCAGCGTCTACT<br>GAGGAAACACTACACGGCAGCTGAAAACGACGATGACTCTGAAGCAAGGGCCC<br>TGAATACAGAGAAGATGAAAACGATGTTGAAAGCTGGGATGACTGTTGACGACT<br>ACGCTGCCAAGCTAAAACTTACCGACAAGATTGCAGCTGCAGCTAACTCTGCAA<br>GGGCGATGGAAAAGCTTGGCGAAACTCTCAAGATGAAGAAGCTCCTGCGGTAC<br>CTCAACTATGTGGCTGAACACACAGCAGTTTGAGCGGCCGC |
| PITG_14787 | XP_002898994 | ATGGACTACAAGGACGACGATGACAAAGTCAAGCTTCTCGAGAATTCCTCTCTTT<br>CTAAGGCTGAAATGAAAAGATTGTTTGAAGCTGGTAATTCTTTGGATGATTTTGC<br>TAAGCATCTTGGTATTGCTGATGATGTTGTTAGAGCTCAATCTTCTAATACTGTTC<br>TTCAAAGATTGATGCAAACTGATGAATATATGAAATATTCTACTTATTTGAATTTTC<br>TTTCTAAACAAAATAAAAAGAAGAAGCCTCCTACTTTTTATCATCTTTGA |
| PexRD3 | ACX46525 | ATGGACTACAAGGACGACGATGACAAAGTCAAGCTTCTCGAGAATTCCCCCGGT<br>GCGGACGCTGTACTAACTGGTGCTGTGTCGCTGGGATTTTTGCAACTCGTTGGT<br>GCTGACCAAAGTGTCATCGAGCAGCCTCGTTTTCTGAGAGACGGTAAGATTGCC<br>GAAGGTGACAACGAAGAGAGAGTGAATGCTCAAAAGGAGGCGGCAGCAAAAGT<br>CCTGGACCAAGTATTCAAGACGAAGTCTTTATCGCCCCTCGACAAACTCGAGAA<br>AACGTCTAATCTGGCGGTGATTCGTCACGTTGCTGCAATGGTAGACGACAAGGT<br>GGATAAAATATTCGCGTTTGCTGACGCTGTGGGAATGGGTCGTGCGTCAATGCT<br>CAAAATGTTGAAGGGTGATAAACAATTCACGGATGCTGAAAGGTTGAAAACTGTC<br>AAGAGGTATGTAAAATTCTTGATCAAGAAGGAACAAAACAACAAGGCTTAA |
| 35a12_90128 | N/A | ATGGATTATAAGGATGATGATGATAAGGTTAAGCTTCTTGAAAATTCTGATGGAC<br>TTCTTTCTATTGTTGAAGAAGTTGATGAACTTATTGAACAACATAATCATATTAGA<br>GAACTTTTTGGAGAATTTTGTCTTGAAGGAAAGACTCCAGAAGAAATTGCTAAGG<br>GAGATACTCATAATAAGAAGACTGTTGAACTTTATAAGAAGTTTGATGCTTATCAT<br>TCTGAACATCATGCTAATGATCAAAAGCAATCTAGAATTGAATGTAAGCTTTCTAA |
| 35a12_Cu10 | N/A | ATGGATTATAAGGATGATGATGATAAGGTTAAGCTTCTTGAAAATTCTGATGGAC<br>TTCTTTCTATTGTTGAAGAAGTTGATGAACTTATTGAACAACATAATCATATTAGA<br>GAACTTTTTGGAGAATTTTGTCTTGAAGGAAAGACTCCAGAAGAAATTGCTAAGG<br>GAGATACTCATGATAAGAAGGCTGTTGAACTTTATAAGAAGTTTTATGCTTATCAT<br>TCTGAACATCATGCTAATGATCAAAAGCAATCTAGAATTGAATGTAAGTTTTCTAA |
| PexRD36 | ACX46558 | ATGGACTACAAGGACGACGATGACAAAGTCAAGCTTCTCGAGAATTCCGCTTGT<br>GCCAAGTCCCGACACTTAAGAGCGAATGGCAAAGACGCCTTGTGGAACTACGA<br>CACAAGCGGCGGTATCAACAGCATCGTAGCGGATGACGAAGAGCGCGTCGTCA<br>ATTTTTCGGGTATCAAGCGATGGCTCAAGGAGCTGTTTAAGAACTGGTCTCAAA<br>GGAACAAGAAGATTCCAGAAGGCACTGAATACGATTTTTTCACCGGAAACTATCA<br>ACAAAATGCTCAAAAGCAGACACGTTCGGCGTGAGCGGCCGC |
| PexRD54 | XP_002903599 | ATGGACTACAAGGACGACGATGACAAAGTCAAGCTTCTCGAGAATTCCGTTGGT<br>CCCTCTTGGCTAGCGAAAGTCGACGGTCTGATGCATAAAATGGTCACGTCCAGT<br>TTGTCTGCGGAAGAAGCGCAGCTCAAAGTGTGGATACAGTCCCAAATACATCCA<br>CGCGAGTTATTCGGGGTCTTAAGTCTTGGCAAGCGTGCAGCAAAGTTGGATGAT<br>AATCCGGATTTCGTGCAGTGGCTCAGGCTTGTAAAGGACTTCCGAGCTAATAAC<br>GGGAACCAGGCGTTTTCAGATCTGGACATTTACTATCTGCTTCTGAAGACAAATA<br>GTCCAGAGCAGTTGAAATTACTTTTCGAAACTCTTCGGCATACTCCCGGAATGAC<br>CAAGATCGGGGCGAGTATGGAGAAGTCTTTATCTGGAAATTGGATTCGCAAAGC<br>CCTAGAACAGGATACGTATCCAACGATCGTTTACAATACGCTTCGTCTTAAAGAC<br>GCTGGCACAAAGCTGGATGATACGCCAATGTTCCGTCAGTGGCTGGAGTATGTA<br>GAGAAGTACTGGAACAAGAATGCGGGCGCTTTCTTCGGGGACACACAAATGTTG<br>ACGTTGTTTCAGAAGACAATGACAGAAGAAGAAGACATCATAAAACTAGTTCACA<br>TGCTTCGAAACAACCCGGGGATGAAGAGTCACGCCGATAAATTGGAGAGGTATT<br>TGCTTCTGACGTCTGAATCGAGCCACAAAACGATGGCTGACGTGTGGCTGAAGG<br>CCCGAGAAACTCCAGAGGAAGTGTTTCGTATTCTACGTTTGGCAGAAAAGCAAA<br>CTGCCGCCGCTGATGACAACCGAATGTTGAATCTGTGGCTCAGGTACACCCAAA<br>CGTACCGGGACAAAATTGACAAGAACGCCTTCTCCGACGCGGAAGCACTGCAG<br>TTTTTCAGGAAGGCCAAACCGCTGGATTTCGACTGGGAAATTGTGTAA |
| PexRD6 | AAA21423 | ATGGACTACAAGGACGACGATGACAAAGTCAAGCTTCTCGAGAATTCCGTTTCA<br>TCCAATCTCAACACCGCCGTGAATTACGCTTCCACATCCAAGATTCGCTTTCTGT<br>CGACTGAGTACAACGCCGATGAAAAAAGAAGCTTGCGAGGTGACTACAACAATG<br>AGGTCACAAAAGAGCCCAACACGTCTGACGAAGAGCGGGCGTTTTCTATCTCAA<br>AGTCTGCGGAATACGTGAAGATGGTACTTTATGGATTCAAACTTGGATTTTCTCC<br>TCGCACTCAGTCCAAGACGGTGTTGCGATACGAAGATAAACTGTTTACGGCCCT<br>CTATAAATTAGGAGAGACGCCGATAAGCCTAAGGACCAAGCATCTCGATAAGGC<br>TTCCGGTAGCGTATTTTTCAACAGATTCAAAAACTGGTACGATAAAAACGTTGGC<br>CCTAGCTAG |
| PexRD8 | XP_002898614 | ATGGACTACAAGGACGACGATGACAAAGTCAAGCTTCTCGAGAATTCCGCCGCC<br>GACGCCTCGGAGCCCATGCCGAATATCGCCAAGTACGCCTCGCCTGAGGTGTC<br>GGTGCACCTTGGCGCTGAGCGTGAGAAGCGTCTCCTCCGTTTCGACTCGAACG<br>ACTACCGTGACGACGACGACGAGGAGGAACGTGCTAACGCTGCTAACCTCTTC<br>AACGTGGACAAGCTCACGGTGTACGTGAACAAGGCTCAGAAGCGTACGGCTAA<br>CAACGTGTCGGGTTCGCTCCTCAACTACTTCAAGCGTCTCGAGGCTTACGGTTA |

| | | |
|---|---|---|
| | | CAACCCGGTGAAGCTCGGTAACATCATCCCGGACGAGGAGTACGACAACCTCC<br>GTATGCTCTACCGTTCGTGGTACTACCACAACAAGTGA |
| Additional pTRBO constructs used to study their effects on 2xFYVE-GFP-labelled endosomes[2] | | |
| AVR1b (*Phytophthora sojae*) | AAM20935 | ATGGACTACAAGGACGACGATGACAAAGTCAAGCTTCTCGAGAATTCCACTGAG<br>TACTCCGACGAAACCAATATCGCCATGGTGGAATCTCCAGATCTCGTCCGTCGC<br>TCGCTCAGGAACGGCGACATTGCCGGTGGAAGATTTCTTCGAGCTCATGAAGAG<br>GACGATGCGGGGGAGCGGACCTTCAGCGTGACTGACCTGTGGAACAAGGTGG<br>CGGCCAAAAAGTTGGCCAAGGCGATGCTGGCGGACCCTTCAAAGGAGCAGAAA<br>GCGTACGAGAAGTGGGCAAAGAAGGGGTACAGCCTGGATAAGATCAAGAACTG<br>GTTGGCAATCGCGGACCCCAAGCAGAAGGGGAAGTACGACCGTATCTACAACG<br>GATACACCTTTCACCGGTATCAGAGCTGA |
| PITG_16737 | XP_002896958 | ATGGACTACAAGGACGACGATGACAAAGTCAAGCTTCTCGAGAATTCCGCTATT<br>TCTATCAATTTTTCTTCCCTTGAAAAGATCTTCAAAAAGGTTACTTCTGCAAAAAC<br>TACAGAGCTCCAAGGTATGCTTAAGGCAGACGAGGCTCTCGGATCCGCATTTAA<br>AACTCTCAAGTTGGGTACAATGAGAATAGGAAAAGATGGTTCTGTTGATCCTAAG<br>ATGGTGGCTAAATTCCTTTCCTCTAGAAATTTTAAGATATGGTCCCAACATGCTG<br>TTAAAATTAACAAGGACGATCCTTATGGAGAAATGTTGAAGGCATTAACTAATGT<br>GTTTGGGGAGAAAAACGTTGCAATGATGATCCTTGTGGGCAATTTGTCTAGGAA<br>CTCCAGGGATGTTGCTAAGAAACTTGAAAAGGCCCAGTTCTATAAATGGTACTTT<br>GTGGACAAGTATAAGACCGCCGATGAGGTTTTCACTAATGTGCTTAAAGCTGATA<br>GAAATAGGATACATGGTTACGGAAGAGAAAAGGAAATTTGGGGTGATTATGCTA<br>AATACGTTACAACTACCGTGATGAAGTATTGA |
| PITG_20144 | XM_002895883 | ATGGACTACAAGGACGACGATGACAAAGTCAAGCTTCTCGAGAATTCCCTATCC<br>ACTATCAAACAGCGAGCACCCGAATCCCCGATGGCTTCGCTCGTTAATTCATATA<br>TTATCAGTCGAAGAATGCTTCGAACAAGCGAGATGGCCGTCTCAGACAACGAAG<br>AGAGGGGAGCGATCGACACCCTGCGCACTAAGCTGAGTAAGCTATTTGAAAAAG<br>TGTTGACGTGGCTTAAGAGAAAGGAAGTCGCAGCTGCAGAGAAAGCGAAGACC<br>AAGAAGCTTGTGGAATTTGTGAAAACTCAAGCTCGCTCCGAAAACCTCGACGAC<br>GCTGTGAAGAATTTGCAGAAGGAAGGCATCACTCATGGCCAGGTCAAGGAATG<br>GCTAAGACTCGACAAGGAACAGAATCAGTGGAAGTTTTTGGACCGCGATAGCCC<br>GCTCTATCTCATGAGTACCAACTTTTTGACAAGCTCAGGGATGCAGCTGTAA |
| NbVAMP72x insert for pENTR vector[3] | | |
| NbVAMP72x | NbS00022342g0004 | <u>CACC</u>GGTCAGCAGAAGGCTTTGATCTATGCGTTTGTGGGTCGTGGGAATGTGGT<br>ATTGGCTGAATATACAGATTTCAGTGGTAACTTCAACTCCATAGCTTATCAGTGC<br>CTTCAGAAGCTCCCTGCTTCCAATAACAAGTTTACTTACAACTGTGATGGTCACA<br>CCTTCAATTACCTTGTCGATAATGGCTTCACATACTGTGTGGTTGCTGAGGAGTC<br>CGTTGGAAGACAAGTTCCAATAGCATTTTTGGAGCGTGTTAAGGATGATTTTGTG<br>TCGAAATATGGGGGTGGGAAGGCTGCCACCGCTCCTCCAAACAGCCTGAACAA<br>GGAATTTGGACCTAAGTTGAAGGAGCATATGCAGTATTGTGCTGACCATCCTGA<br>TGAAATTAGCAAAATTGCGAAGGTGAAAGCCCAGGTTTCAGAAGTTAAAGGTGTT<br>ATGATGGAAAACATTGAGAAGGTTCTTGATCGTGGAGAAAAGATAGAGCTTCTTG<br>TGGATAAGACGGAGAACCTTCATCACCAGGCACAAGACTTCAGGAATACTGGAA<br>CCCAAATCCGGAGGAAAATGTGGCTGCAAAACATGAAGATCAAGCTGATAGTGT<br>TAGGTATCCTGATTGCTTTGATCCTCATCATTGTTCTCTCAGTTTGCAAGGGATT<br>CAATTGTGGAAAGTAA |

[1]Residues in red color indicate SNPs compared with the predicted gene model in *P. infestans* T30-4 or A. thaliana Col-0; Golden gate overhangs are underlined; start/stop codons are in bold when present.
[2]Coding nucleotide sequences are full open reading frames from start to stop codons (inclusive) incorporating the FLAG tag sequences. CDS were codon-optimized for expression in *Nicotiana benthamiana*. They were synthesized and cloned into pTRBO using PacI (5'-end) and NotI (3'-end) restriction sites by GenScript Ltd.
[3]Nucleotide sequence added for Topo cloning is underlined.
